# Interfacial water confers transcription factors with dinucleotide specificity

**DOI:** 10.1101/2023.10.03.560647

**Authors:** Ekaterina Morgunova, Yimeng Yin, Fangjie Zhu, Tianyi Xiao, Ilya Sokolov, Alexander Popov, Charles Laughton, Jussi Taipale

**Author notes:** Corresponding author: Jussi Taipale (*).

## Abstract

Transcription factors bind to DNA by recognizing specific bases within their binding motifs. Binding to each DNA mononucleotide within the motif often contributes independently to total binding energy. However, some transcription factors (TFs) can bind to DNA more specifically than predicted by this model, by directly recognizing DNA dinucleotides. To understand this process, we have solved the structures of the basic helix-loop-helix protein MYF5, and the homeodomain protein BARHL2 together with DNA containing a set of dinucleotides that have different affinities to the proteins at high resolution (< 1 Å). We observe that dinucleotides can be recognized either enthalpically by an extensive water network that connects the adjacent bases to the TF, or entropically by formation of a hydrophobic patch that maintains water mobility at the protein-DNA interface. The two distinct thermodynamic signatures of the two equally optimal sites also confer differential temperature sensitivity to the optimal sites, with implications for thermal regulation of gene expression. Our results uncover the enigma of how TFs can recognize more complex local features than mononucleotides, and demonstrate that water-mediated recognition is important in predicting affinities of macromolecules from their sequence.

## INTRODUCTION

Proteins can bind to DNA in a non-sequence specific manner by binding to its backbone, or recognize specific sequence by direct interactions with the DNA bases. Specific hydrogen bonding or hydrophobic interactions between amino-acids and DNA bases result in recognition of individual mononucleotides, with total binding energy being approximately additive^1–6^. The additive model holds very well for bases that are distal to each other, but often performs less well in predicting energy contributions for adjacent bases. Due to the influence of adjacent bases on each other, TF binding to DNA can be better approximated by models that use dinucleotide instead of mononucleotide features^7,8^. It is well established that dinucleotide content affects the local dynamics and structure of the DNA backbone, and the relative orientation of the bases^9^. For example, nucleosomes bend DNA, and preferentially bind to DNA sequences where particular dinucleotides are located at specific positions relative to the direction of the DNA bending (Reviewed in ^10^). Some transcription factors can also affect the conformation of the DNA backbone ^11–18^, and their specificity towards dinucleotides could similarly be explained by the contribution of the dinucleotides to the structure and flexibility of the DNA backbone. However, the structural distortion caused by histone octamer or DNA-bending TFs is expected to result in relatively weak dinucleotide preferences, and thus cannot account for highly specific recognition of dinucleotides observed for TFs. Furthermore, many TFs having preferences for particular dinucleotides can bind to DNA without inducing major changes to its canonical B-form, suggesting the existence of additional mechanisms for recognition of dinucleotides.

Previously, we have shown that binding of TFs to two distinct sequence optima can be caused by partial independence of the two contributions to binding energy, ΔH and TΔS, with different DNA sequences representing entropic and enthalpic optima for binding^19^. We also proposed that such a thermodynamic mechanism affects many other macromolecular interactions. To investigate the molecular interactions that contribute to the entropic and enthalpic optima, and enable TFs to specifically recognize dinucleotides, we have in this work characterized the structures of MYF5 and BARHL2 bound to multiple different optimal and sub-optimal DNA sequences.

## RESULTS

### Mechanism of MYF5 binding to dinucleotides

To investigate how basic helix-loop-helix (bHLH) proteins can recognize dinucleotides flanking the E-box core sequences, and bind to two distinct DNA sequences with similar affinity^7,19–23^, we solved the structure of the bHLH domain of MYF5 homodimer bound to two different DNA fragments representing the entropic and enthalpic optima (^19^; **Fig. 1a,b**). One sequence is palindromic, and contains an E-box like core CAGCTG ^24^, flanked by the entropically optimal GT (**GT**CAGCTG**AC**;^19^, MYF5:DNA^GT^ hereafter), whereas the other sequence, **AA**CAGCTG**AC** (MYF5:DNA^AA^ hereafter) is non-palindromic, containing one enthalpic half-site (**AA**CAG), and one entropic half-site (CTG**AC**, the reverse complement of **GT**CAG).

**Figure 1.**
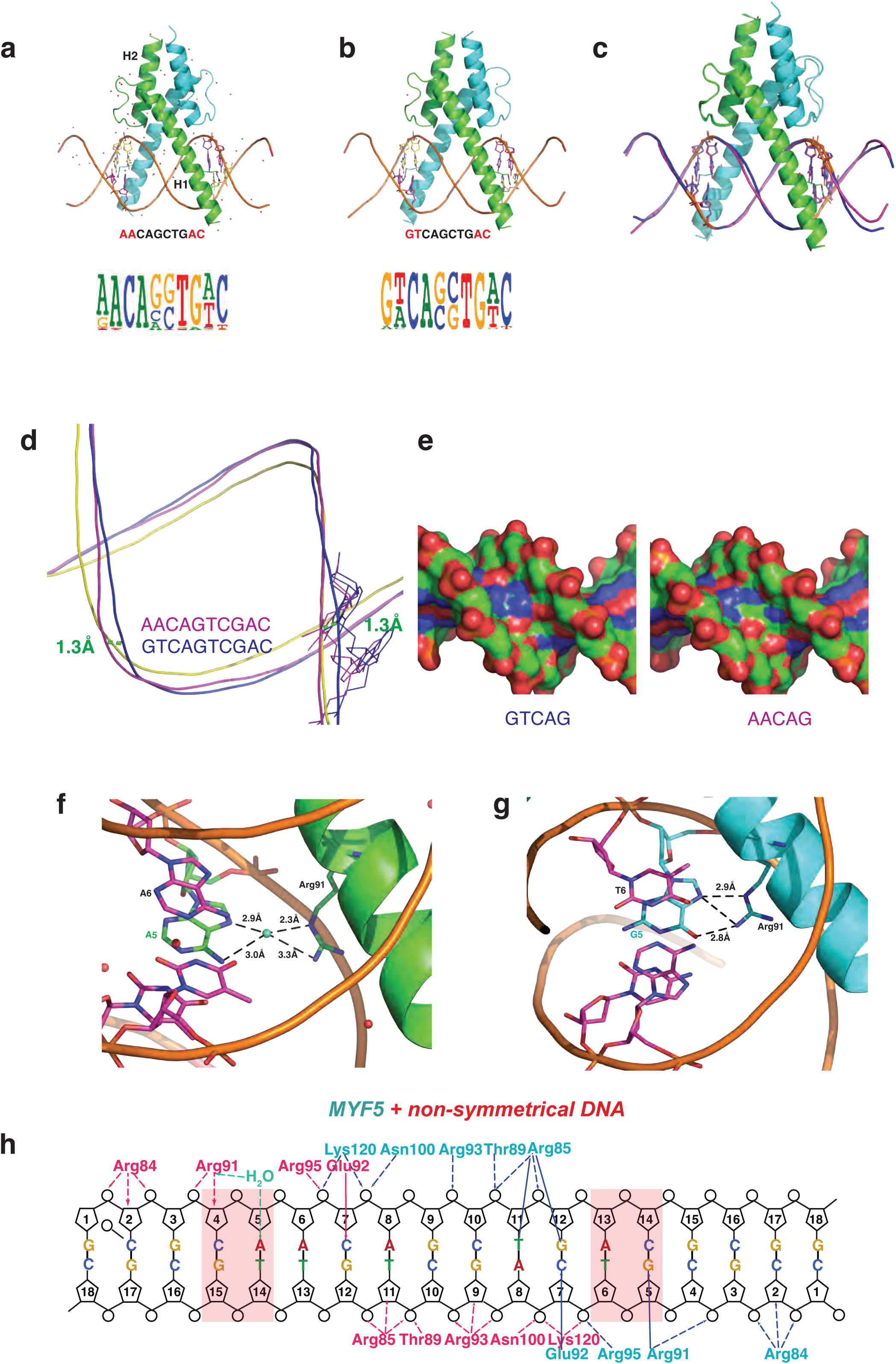
Structures of MYF5:DNA complexes. **a, b**, Overview of two structures of MYF5 DNA-binding domains bound to non-symmetrical DNA **AA**CAGCTG**AC** (**a**) and symmetrical DNA **GT**CAGCTG**AC** (**b**). Logos of the motifs of the homologous protein MYF6 are shown below the structures. Note that MYF6 prefers to bind to either AA or GT flank before the E-box. **c**, Structural alignment of two MYF5 structures containing different DNA sequences. The symmetrical DNA **GT**CAGCTG**AC** is in magenta, the non-symmetrical DNA **GT**CAGCTG**TT** in blue. **d**, Shape of DNA backbone near the divergent dinucleotide. Canonical B-shape DNA is in yellow. The measured distance between the corresponding phosphorus atoms is shown as green dash lines. **e**, Charge distribution on the surface of the region containing the divergent dinucleotide (blue is positive, red is negative). Note that charge distribution is very different at the region of the GT and AA dinucleotides. **f**, A water molecule mediates the contact between Arg91 and A_5_ of the AA dinucleotide. **g**, Direct contact is formed between Arg91 and G_5_ of the GT dinucleotide. **h**, Schematic representation of the contacts formed by MYF5 with non-symmetrical DNA. Left and right sides show contacts to AA and GT, respectively.

The structures were solved at 3.1 Å and 2.3 Å resolution using X-ray crystallography (see **Methods**). The overall fold of the bHLH domain is strikingly similar to that observed in other members of this family, including MyoD, Max and c-Myc (RMSD 1.2 Å; **Extended Data Fig. 1**). Each MYF5 monomer forms two long helices connected by a nine-residue loop. The first helix (H1) contains the basic region that binds to DNA, whereas the second helix (H2) is responsible for dimerization (Ref.^25^; **Fig. 1a**). As observed in all known bHLH-DNA complexes, the basic region fits into the major groove without significantly bending the DNA (**Fig. 1a-e; Extended Data Fig. 1**).

To determine whether changes in DNA backbone shape could explain the recognition of the two flanking dinucleotides, we performed comparative analysis of the DNA shape with Web 3DNA2.0 server^26^. This analysis revealed that the DNA shape in both the MYF5:DNA^GT^ and MYF5:DNA^AA^ is very similar to canonical B-DNA, with only minor differences (**Fig. 1d**). The major groove of the DNA^AA^ is wider in steps CA and TG but narrowed in the central step GC. The major groove of DNA^GT^ is less distorted although the central step GC is narrowed as well. Taken together, comparison of the shape (**Fig. 1d**) and electrostatic potential (**Fig. 1e**) of the two DNAs at the position of the variable dinucleotide shows a relatively small difference of 1.3 Å between backbone shapes; larger difference is observed between the electrostatic potentials in the major groove, suggesting that DNA backbone shape and electrostatic potential cannot explain the preference of MYF5 to flanking AA and GT dinucleotides.

We next analyzed details of DNA base contacts in the complexes. For both complexes, only two direct contacts are formed between the core E-box (CAGCTG) sequence and the MYF5 protein: the side chain oxygen of Glu92 of one MYF5 monomer hydrogen-bonds to the nitrogen 4 (N4) of the first C (C_7_), and a symmetric contact is made by the other MYF5 monomer to the C complementary to the last G of the E-box (**Fig. 1h**). In addition to the specific base contact, DNA affinity is increased by non-sequence specific hydrogen-bonds formed by Arg85, Thr89, Arg93 and 95, Asn100 and Lys120 with the ribose oxygens and phosphate moieties. The AA flank in DNA^AA^ is recognized by Arg91, which makes contact with the first A (A_5_) through a water molecule (**Fig. 1g**; **Extended Data Fig. 1c-e**). However, in DNA^GT^, Arg91 does not interact with water, but instead makes direct hydrogen bond contact to the G_5_ (**Fig. 1h**).

These results suggest that the dual specificity of MYF5 is not due to similarities in DNA shape or electrostatic potential between the recognized dinucleotides. Instead, the two high-affinity sites appear to result from the entropic and enthalpic optima^19^; in the entropic optima, the G in the GT dinucleotide is bound directly, freeing a water, whereas in the enthalpic optima, the first A of the AA dinucleotide is bound via a fixed water molecule. However, due to the relatively low resolution of the MYF5 structures, we were unable to determine why the G-bound state prefers an adjacent T and A-bound state prefers an adjacent A. This motivated us to look for an additional model representing another TF family, which would support the generality of our findings and enable the analysis of TF:DNA structures at higher resolution.

### Mechanism of BARHL2 binding to dinucleotides

To study TF binding to dinucleotides in more molecular detail, we performed an extensive structural analysis of the homeodomain TF BARHL2 bound to different DNA sequences. BARHL2 binds with high affinity to a sequence resembling a canonical homeobox (TAA**TT**G), but unlike most other homeodomain proteins, can also recognize a sequence TAA**AC**G that contains a different dinucleotide, AC, with even higher affinity. To investigate BARHL2 specificity at the region containing the dinucleotides, we solved BARHL2 structures bound to eight different DNAs, including the enthalpic (TAA**AC**G) and entropic (TAA**TT**G) optima, two more weakly bound sequences that are within one substitution from both optima (TAA**AT**G and TAA**TC**G), and four suboptimal sequences (TAAT**G**G, TAA**GT**G, TAA**GC**G, TAA**CC**G). The structures were of high to extremely high resolution (from 2.6 to 0.95 Å), enabling detailed analysis of the interactions contributing to the binding affinity of each sequence. For clarity, we refer to the complexes using the variable dinucleotides hereafter (e.g. TAA**AC**G is DNA^AC^ hereafter). One of the crystals of BARHL2:DNA**^AC^**complex diffracted to 0.95 Å which is so far the highest resolution achieved for TF:DNA complexes (for other high-resolution structures, see Refs ^27,28^). The resolution of this complex was so high that it revealed alternative conformations for most of the phosphates, sugars and bases of the DNA backbone (**Fig. 2a**; **Extended Data Fig.2**), hitherto only observed in two previous TF:DNA complexes^27,28^. Such high resolution allowed us to investigate the protein-DNA-water interactions in unprecedented molecular detail.

**Figure 2.**
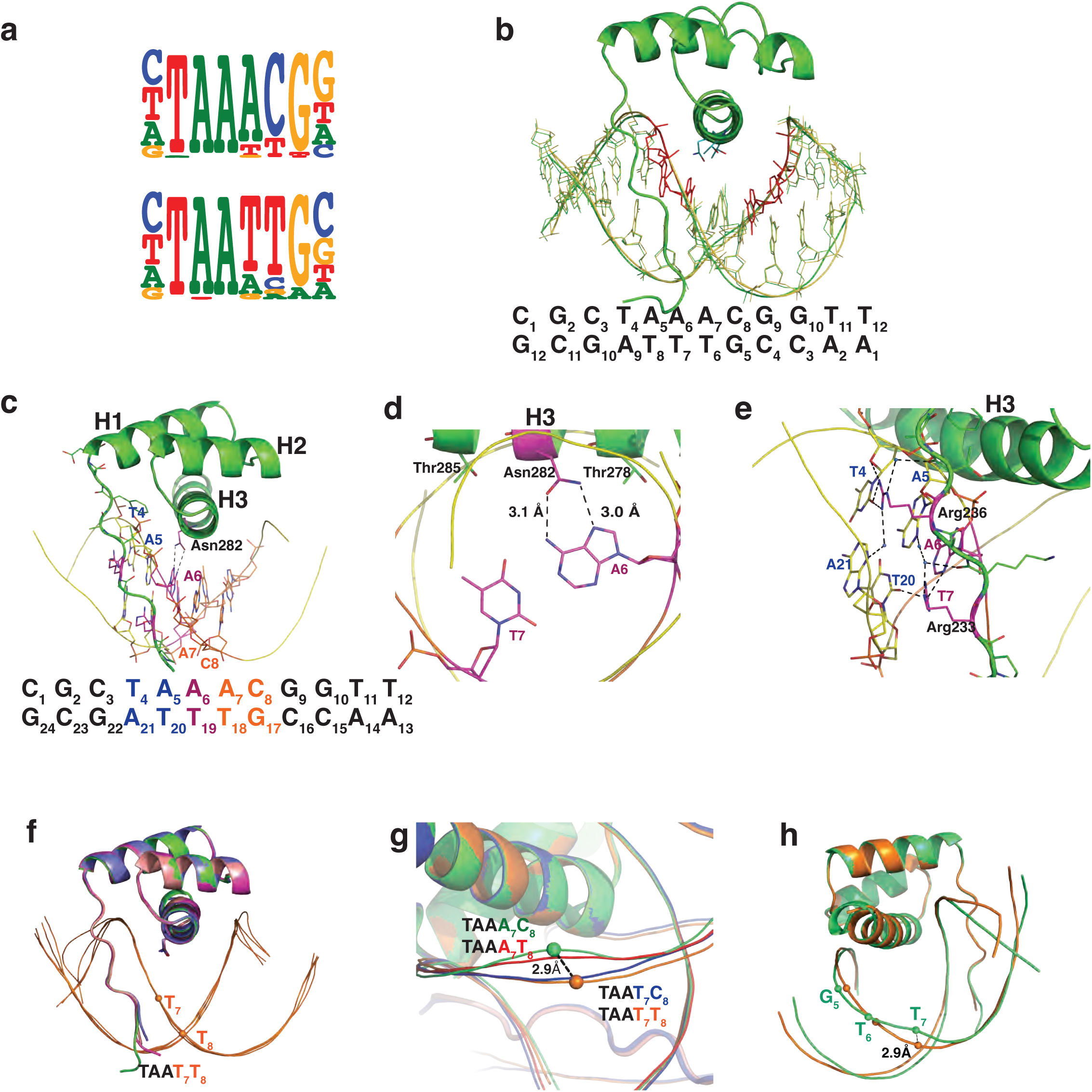
Structures of BARHL2:DNA complexes. **a**, Sequence logos for the two distinct DNA binding specificities of the BARHL2 protein. Note that BARHL2 binds to the dinucleotide after the TAA sequence, preferring either AC or TT. **b**, Overall structure of BARHL2 bound to the DNA C_1_G_2_C_3_T_4_A_5_A_6_**A_7_C_8_**G_9_G_10_T_11_T_12_ (BARHL2:DNA**^AC^**). The two conformations of the DNA backbone sugars and bases observed in the ultra-high-resolution structure (0.95 Å) are indicated in green and yellow. The bases that only show a single conformation are A_5_A_6_ of the direct strand and C_4_G_5_ of the complementary strand are presented as a ball-and-stick models in red. The sequence of the double stranded DNA is shown below. **c**, **d**, BARHL2 interacts with TAA part of its recognition sequence using a similar mechanism as other homeodomains. Close view of the interaction between A_6_ (magenta) and the canonical Asn282 is shown in (**d**). The divergent dinucleotides are in orange. **e**, Close view to the interactions via the minor groove. The T_4_A_5_ sequence (blue) is bound via minor groove contacts formed between the amino-terminal tail residues Arg233 and Arg236 (**e**). **f**, Structural alignment of four subunits of the complex BARHL2:DNA**^TT^**. The largest shift for DNA strands between 4 subunits is <1 Å. **g**, Alignment of four complexes with different DNAs. Colour codes are: red, DNA**^AT^**; blue, DNA**^TC^**; orange, DNA**^TT^**; green, DNA**^AC^**. The corresponding DNA sequences are written using the colour code of the figure. **h**, Structure of DNA backbone in the part of divergent dinucleotides of DNA**^TT^** and DNA**^AC^**is 2.9 Å.

Analysis of all the complexes confirmed that BARHL2 exhibits common homeobox features and encompasses a comparatively long, positively charged N-terminal peptide and three α-helixes connected by short loops (**Fig. 2b**). Like canonical homeodomain proteins, BARHL2 interacts with the DNA major groove using the third α-helix (H3, the recognition helix), and with the minor groove using its N-terminal peptide (**Fig. 2b-e**; **Extended Data Fig. 3**). In all solved homeodomain:DNA complexes, the last A of the TA**A** sequence is recognized by an asparagine (Asn282 in BARHL2), which forms a hydrogen bond with **A_6_** (T_4_A_5_**A_6_**). In BARHL2, Thr278 also forms a hydrophobic contact with carbon atom C8 of the aromatic ring of the same adenine. The TAA sequence is also recognized by Arg233 and Arg236 via the minor groove, forming direct and water-mediated hydrogen bonds with all three base pairs (**Fig. 2e**). These results indicate that, within the conserved TAA sequence, the interactions between BARHL2 and DNA are typical of those seen for the characterised anterior-type homeobox TFs.

We next analyzed the molecular basis of the sequence specificity of BARHL2 towards the dinucleotides following the TAA motif. First, we checked whether the high affinity of the two distinct optimal sequences DNA^AC^ and DNA^TT^ could be explained by a similarity in DNA backbone shape. However, all DNAs in all structures had almost canonical B-shape (e.g. DNA^TT^, **Fig. 2f**). Comparison of the DNAs between the DNA^AC^, DNA^TT^, DNA^AT^ and DNA^TC^ revealed that the largest shift from B-shape was observed at the position of the variable dinucleotide between DNA**^AC^** and DNA**^TT^** (2.9 Å; **Figs. 2g and h**). Thus, the two high affinity sequences were farther apart from each other than from the two lower affinity sequences, suggesting that the differences in affinity are unlikely to be caused by differences in DNA backbone shape.

To investigate whether the dinucleotides would instead be recognized by networks of water molecules, we studied the arrangement of water molecules in the protein-DNA interfaces. We first assessed whether differences in resolution would affect detection of the water molecules using the two BARHL2:DNA^AC^ structures solved at 0.95 Å and 1.3 Å. Inspection of the water content in the protein:DNA interface at both resolutions showed conserved positions of water molecules, indicating that at this range, resolution does not materially affect detection of water molecules contributing to protein-DNA recognition (**Extended Data Fig. 2g**).

We then compared the water arrangements and molecular interactions between BARHL2 and the eight different DNA sequences (**Fig. 3**). Analysis of the highest-affinity complex, BARHL2:DNA^AC^, revealed that the high affinity binding of BARHL to DNA^AC^ is due to a combination of indirect and direct recognition of DNA. Key amino-acids involved in recognition of the AC dinucleotide and its complementary bases are asparagine 282 and two threonines, 278 and 285. Thr278 connected to DNA by two water chains (**Fig. 3a**). The first water chain contains three water molecules which together connect the side chain oxygen of Thr278 with the side chain oxygen of Asn282 and N7 of the A of the AC dinucleotide (A_7_). On the other side, Thr278 is connected to N4 of the C of AC (C_8_) by a second chain of four water molecules. Thus, Thr278 orders water molecules in such a way that they contact both bases of the AC dinucleotide. The two water chains are also connected by one water molecule, which is within a hydrogen-bond distance from N6 of A_7_ and O6 of G_17_ opposite to C_8_ of AC. The T_18_ opposite to the A_7_ of the AC dinucleotide is also recognized directly by a hydrophobic contact to the methyl group (CG2) of Thr285 (**Fig. 3a**). In addition, the side-chain oxygen of Thr285 contacts N7 of the G_17_ opposite to the AC dinucleotide via a water molecule that is a part of the second water chain (**Fig. 3a**). Furthermore, the T_18_ opposite to the A_7_ of AC is connected via a water molecule to the side-chain oxygens of Asn282 and Thr285. This water molecule is in a network that includes another water which in turn is connected to N7 of G_17_ opposite to C_8_ of AC. Taken together, these results show that DNA^AC^ is bound by a combination of a direct hydrophobic interaction and an extensive network that forms in the major groove between the protein and DNA. The entrained water chain spans the interfacial contact surface between the recognition helix and DNA (**Fig. 3a**).

**Figure 3.**
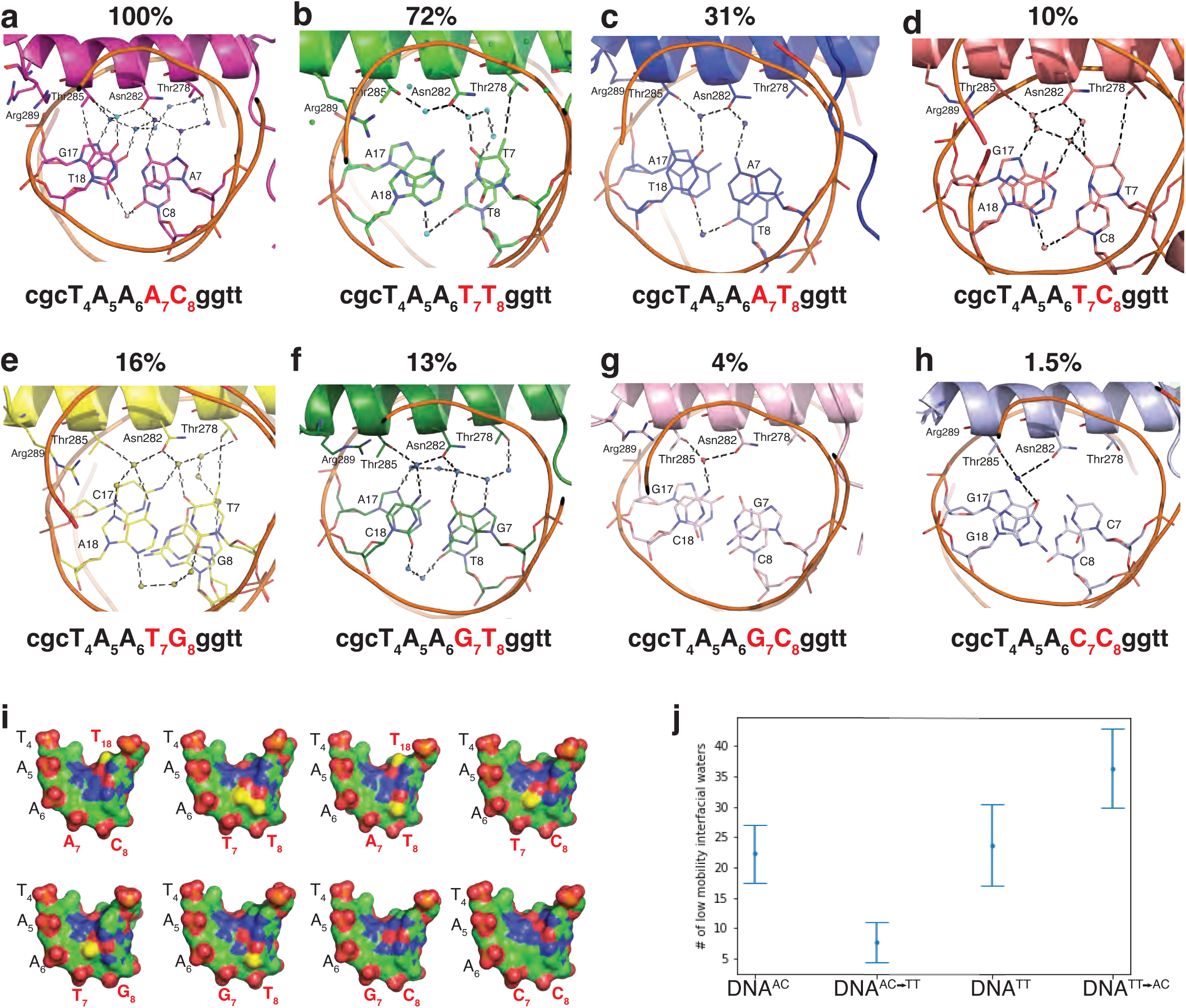
The role of water molecules in BARHL2:DNA complexes. The analysis of eight structures of BARHL2 containing the different dinucleotide. **a**, BARHL2:DNA**^AC^**; **b**, BARHL2:DNA**^TT^**; **c**, BARHL2:DNA**^AT^**; **d**, BARHL2:DNA**^TC^**; **e**, BARHL2:DNA**^TG^**; **f**, BARHL2:DNA**^GT^**; **g**, BARHL2:DNA**^GC^**; **h**, BARHL2:DNA**^CC^**; The enrichment of the respective 6-mer relative to TAAACG is shown above the panels. Panels **b**, **c**, **d**, **g**, and **h** show the complexes containing the largest number of water molecules of the four complexes of the respective asymmetric units. The structures on panels **a** and **f** contain only one complex in the asymmetric unit. **i**, Surface representation of the DNA binding sites. Carbon, oxygen, nitrogen and phosphorus atoms are green, red, blue, and orange, respectively, and the CH3 group of thymidine is in yellow. **j**, Analysis of interfacial water mobility using Molecular Dynamic simulation. Left: simulations run using the DNA**^AC^** lattice with or without *in silico* mutation of the AC dinucleotide to TT. Right: simulations run using the DNA**^TT^** lattice with or without *in silico* mutation of the TT dinucleotide to AC. Note that in both cases, AC dinucleotide has higher percentage of low mobility waters at the protein:DNA interface.

The molecular interactions observed in the structure of BARHL2 bound to DNA^TT^, which showed the next highest affinity in the SELEX experiments, were strikingly different from those found in the BARHL2:DNA^AC^ complex. The extensive water-network is lost due to a hydrophobic patch in the major groove, formed by the methyl-groups of the two Ts (**Fig. 3b**). The hydrophobic contact with Thr285 is replaced by hydrophobic contact between the methyl group (CG2) of Thr278 and C7 of the first T (T_7_) of TT. Only one water connecting the side-chain oxygen of Thr285 with the side-chain oxygen of Asn282 was observed in the area around Thr285. On the other side the Asn282 is connected to O4 of T_7_ via one water and to O4 of the second T (T_8_) of TT via a water chain containing three molecules. Thus, most of the contacts in the complex BARHL2:DNA^TT^ are shifted from the Thr285 area to the Thr278. An additional observation is that the side chain of Arg289 forms a hydrophobic contact with the methyl group of Thr285, in contrast to the BARHL2-DNA^AC^ where Arg289 shows two conformations, both of which are turned away from Thr285 (**Fig. 3b**).

The role of low-mobility water molecules in recognition of the BARHL2:DNA^AC^ compared to BARHL2:DNA^TT^ was also supported by molecular dynamics simulations. More low-mobility waters were observed in crystal lattice simulations of BARHL2:DNA^AC^ complex compared to the same complex where the DNA had been *in silico* mutated to DNA^TT^. The converse mutation, from TT to AC also led to a larger number of low-mobility waters in the BARHL2:DNA^TT^ complex crystal lattice (**Fig. 3j**; see **Methods** for details).

The ability of BARHL2 to specifically recognize dinucleotides is shown by the fact that it has the highest affinity to DNA^AC^ and DNA^TT^. If BARHL2 would interact with mononucleotides within a sequence motif independently, the highest affinity sequence would be one of the sequences (either DNA^AT^ or DNA^TC^) that is one base substitution (Hamming distance) away from both DNA^AC^ and DNA^TT^. We next investigated why DNA^AT^ has lower affinity to BARHL2 than the two optimal sequences. The DNA recognition pattern in BARHL2:DNA^AT^ complex is more similar to DNA^AC^ than DNA^TT^. Compared to DNA^AC^, the hydrophobic contact between Thr285 and T_18_ opposite to A_7_ of AC is retained, but the extensive water network connecting both Thr285 and Thr278 with the AC region is almost completely lost. There is only one water molecule connecting the side-chain oxygen of Thr285 with the side-chain oxygen of Asn282 and one water molecule connecting Asn282 with N6 of A_7_ of AC (**Fig. 3c**). In addition, the side chain of Arg289 has only one conformation, rotated away from the protein-DNA interface.

Another sequence, DNA^TC^ is also only one Hamming distance away from both local optima, but has even weaker affinity than DNA^AT^. The recognition pattern appears like a mixture of DNA^TT^ and DNA^AC^, with the hydrophobic contact of DNA^TT^ between C7 of T_7_ and the methyl group of Thr278 retained, but at a suboptimal distance (4.3 Å; **Fig. 3d**). The water molecules are arranged near Asn282 in a somewhat similar pattern to those observed in DNA^AC^, with the water-net connecting the side-chain oxygens of Thr285 and Asn282 and N7 and O6 of G_17_ (opposite to C of AC). Thus, DNA^TC^ combines aspects of recognition from DNA^TT^ and DNA^AC^, but both types of interactions are less optimal, explaining the relatively low affinity of the BARHL2:DNA^TC^ complex.

In addition to studying sequences between the two optimal sites, we also solved structures of BARHL2 bound to sequences that are close to TT and have relatively high affinity. Mutations of the second T (T_8_) to G_8_ decreased affinity less than the mutation to C. Overall, the recognition pattern in the DNA^TG^ complex is similar to that observed in the DNA^TC^ (**Fig. 3e**). However, the non-polar contact between Thr278 and T_7_ has a more optimal distance. The water network is also more extensive, with three more water molecules. Those water molecules connect side-chain oxygen of Thr278 to both O6 and N7 of one of the observed conformations of G_8_. The DNA^TG^ complex also shows very strong distortion of the DNA backbone, with a 3.7 Å shift between two conformations of the phosphate backbone at the position of the G nucleotide located after the TG sequence **(Extended Data Fig. 2f**).

The replacement of T_7_ with G led to a substantial loss of affinity; in BARHL2:DNA^GT^, neither Thr278 nor Thr285 makes direct hydrophobic contact with bases. Instead, water molecules are spread rather equally near both amino-acids. The side-chain oxygen atoms of Thr285, Asn282 and N4 of C_18_ are in contact through one water which is connected to the other one contacting N7 of A_17_ and simultaneously participating in a five-water chain connecting both O6 and N7 of G_7_ with the side-chain oxygen atoms of Asn282 and Thr278 (**Fig. 3f**).

Mutation of the AC sequence led to a much larger loss of affinity than mutation of TT. The AC sequence had high affinity due to a high enthalpic contribution of the elaborate water-network connecting amino-acids and bases. Consistently, mutating the A_7_ to either G or C in the BARHL2:DNA^GC^ and BARHL2:DNA^CC^ complexes resulted in a loss of most water-mediated contacts, and a very low affinity. Both complexes showed only one water molecule in the interface connecting the side-chain oxygens of Thr285 and Asn282 with N4 of C_18_ or O6 of G_18_, respectively (**Fig. 3g,h**). Furthermore, those complexes did not show any hydrophobic contacts involving either Thr278 or Thr285, further explaining their very low affinity.

### Role of hydrophobic interactions and water-mediated interactions

To determine the role of hydrophobic interactions and water-mediated bonds in BARHL2:DNA interactions, we mutated the two threonines 278 and 285 that contribute to the hydrophobic interactions and water-chain organization to residues found in other homeodomain proteins (**Extended Data Fig. 3**). The protein-DNA affinities were measured using SELEX. As expected, binding of BARHL2 to the TAATT sequence, which is commonly recognized by homeodomains, was not abolished by most mutations of the two threonines to residues present in other homeodomains (**Fig. 4a**). However, binding to the BARHL-family specific TAAAC sequence was very sensitive to mutation. Significant binding to TAAAC was retained only in two cases, where T278 was mutated to residues that retained the methyl but not hydroxyl group (isoleucine or valine, residues present in NKX1.2 and EMX1, respectively). These results support the importance of T278 and T285 in organizing the water network that recognizes the BARHL2-specific TAAAC sequence.

**Figure 4.**
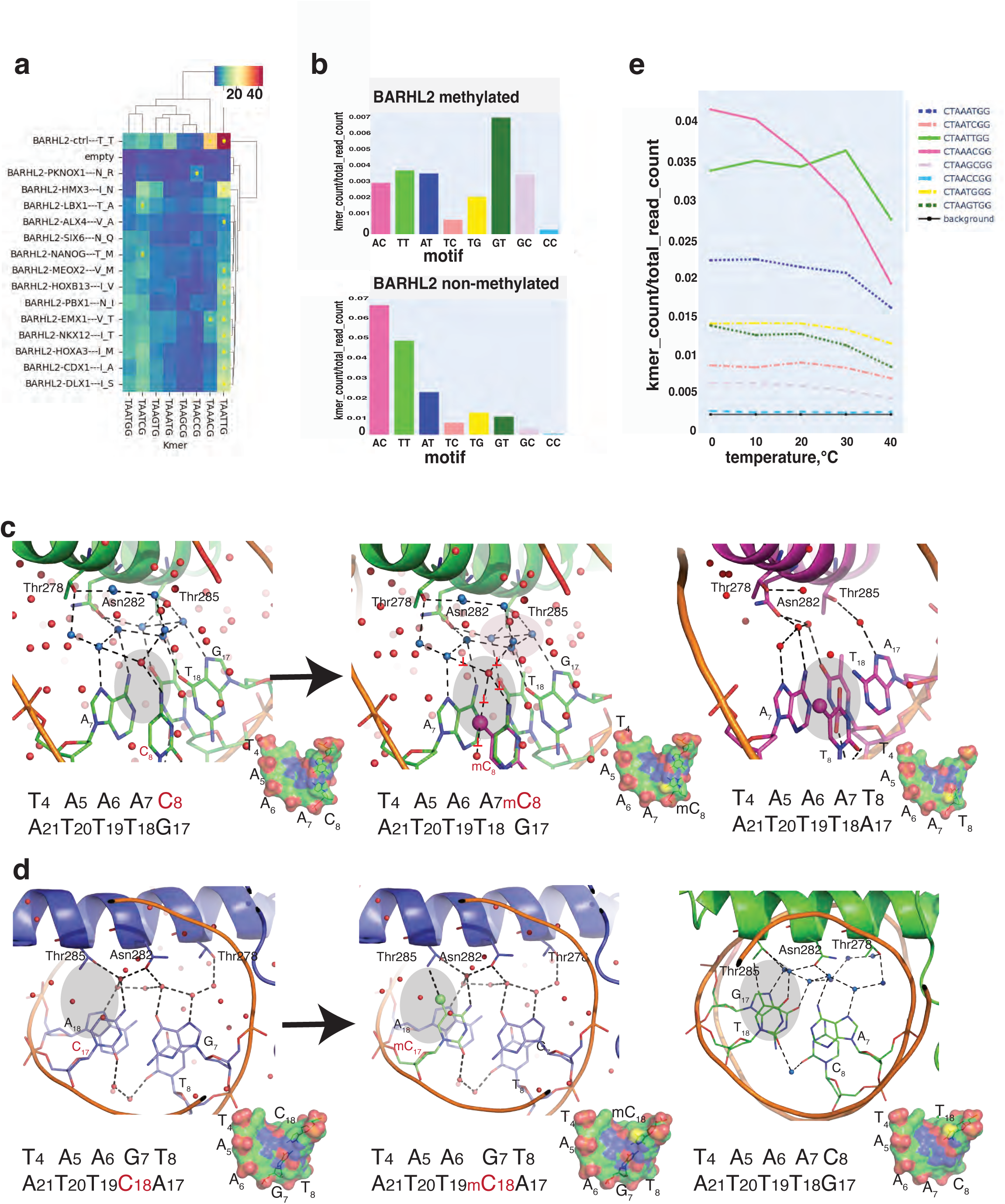
Effects of mutations, methylation and temperature changes on BARHL2 binding to DNA. **a**, The effect of mutation of Thr278 and Thr285 on binding of BARHL2 to the different homeobox-like sequences. Color indicates relative enrichment of the indicated sixmer in 4th SELEX cycle compared to the initial DNA library. Yellow circles indicate that the sixmer represents local maxima (enrichment is higher than that of related sequences). Labels indicate the amino-acids into which Thr278 and Thr285 were mutated, and the TF that contains those residues. **b**, The effect of methylation of the homeobox-like sequences on affinity. Top and bottom panels show relative enrichment of TAANNG sixmers containing the indicated dinucleotide in the NN position in the presence and absence of cytosine methylation, respectively. **c**, Effect of methylation of the C in BARHL2:DNA^AC^. The 5-methyl group of C_8_ is shown in ball and stick and colored magenta. Note that the addition of a methyl group to DNA^AC^ (left) makes the DNA^AmC^ (middle) similar to DNA^AT^ (right) and destabilizing the water-network between BARHL2 and DNA. **d**, BARHL2 has relatively low affinity to DNA**^GT^** (left); however, methylation of C complementary to the G (middle) increases the affinity, most likely because similar to the T complementary to the A of DNA**^AC^** (right), the 5-methyl group of C_18_ can form a hydrophobic contact to Thr285. **e**, Effect of temperature on BARHL2 affinity to different homeobox sequences. Note that temperature affects affinity of kmers containing AC (magenta) and TG (dark green) more than those containing TT (light green).

To further assess the role of hydrophobic contacts in BARHL2:DNA recognition, we tested the effect of methylation of cytosines on BARHL2:DNA affinity. Cytosine methylation introduces a hydrophobic methyl group to the 5 position of C, enabling methyl-C to take part in similar hydrophobic interactions as a T. In addition, the hydrophobic methyl group destabilizes local water networks. To test the effect of methylation of Cs at all positions and dinucleotide contexts, we directly introduced 5-methylcytosine to DNA using PCR, and then determined the enrichment of different sequences using SELEX. Two strong effects were detected: First, the methylation of C-base in DNA^AC^ reduced BARHL2:DNA affinity (**Fig. 4b**). The lower affinity was most likely due to the methyl group interfering with the elaborate water network of the BARHL2:DNA^AC^ complex (**Fig. 4b**). Second, methylation of the C on the complementary strand of DNA^GT^ greatly increased the affinity, to a level even higher than that observed in both unmethylated optimal sequences, DNA^AC^ and DNA^TT^. The high affinity is most likely due to the ability of mC but not unmethylated C to form a hydrophobic contact to Thr285 (**Fig. 4b-d**).

To further test the role of enthalpic water-mediated interactions in BARH2:DNA binding, we performed the HT-SELEX at a series of different temperatures (0°C, 10°C, 20°C, 30°C, 40°C). As the entropic contribution to binding depends on temperature, comparing binding at different temperatures can reveal which sequences are bound more entropically, and which depend more on enthalpic contribution to affinity. Comparison of the effect of temperature on relative binding affinity of the different sequences revealed that the affinity to the enthalpic TAAAC sequence decreased when temperature was increased; a similar but less dramatic trend was also observed for TAAGT. However, the sequence representing the entropic optima TAATT was less affected by temperature (**Fig. 4e**). Taken together, these results are consistent with the role for extensive water-networks on binding of BARHL2 to the TAAAC and methylated TAAGT sequences, and a dominant entropic contribution of binding to TAATT^19^.

## DISCUSSION

The currently established view of the binding specificity between protein molecules and DNA is that specific binding results from three main types of interactions: shape recognition, formation of specific hydrogen bonds between amino acids and bases and the formation of water-mediated contacts^29^. Direct interactions between individual amino-acids and DNA bases can be used to differentiate between mononucleotides, but because of their one-to-one nature, individual contacts are generally not very sensitive to neighbouring base content. The influence of sequence on DNA conformation, in turn, can be used to recognize dinucleotides, as the bending modulus of the DNA helix towards a particular direction is sensitive to local dinucleotide content^29,30^. Other geometric aspects of DNA, such as the width of the minor groove are also affected by dinucleotide content, and narrowing of the minor groove caused by homopolymeric stretches of A or T can be sensed by arginine residues^31^. It has been less clear how DNA-binding proteins that do not substantially distort DNA shape can preferentially bind to two distinct dinucleotides. In this work, we have used structural biology to determine how two TFs, MYF5 and BARHL2, that represent different structural families, can specifically recognize dinucleotides without dramatically bending the DNA. Our results show that dinucleotide recognition critically depends on water molecules at the protein-DNA interface.

Recently, computer simulation methods that provide spatially resolved maps of hydration thermodynamics in protein-ligand systems have been developed. Spatial decomposition of translational water-water correlation entropy in Factor FXa^32^, and Crambin^33^ showed that thermodynamic driving forces linked to hydration can play an important role in protein folding and ligand binding^34^. In the case of Crambin, more than half of the solvent entropy contribution came from induced water correlations^33^, highlighting the importance of water-water interactions. Because both free DNA and the interface between protein and DNA contain large numbers of water molecules, the binding reaction commonly leads to large trade-offs between entropy and enthalpy. We have shown earlier that individual TFs can bind to two distinct local sequence optima, one representing entropic optima, where water-molecules flow freely, and the other representing enthalpic optima, where water molecules at the protein-DNA interface are immobilized into a complex network^19^. We show here that the water correlations contribute to recognition of a dinucleotide within the BARHL2 recognition sequence (**Fig. 5**), which can be recognized entropically by formation of a local hydrophobic patch that repels water (BARHL2:DNA^TT^), or enthalpically by an extensive water-network that links the two DNA nucleotides to the same amino-acid (BARHL2:DNA^AC^). In both cases, the dinucleotides are preferred because of local collective action of the water-molecules. Sequences between TT and AC lead to compromises that weaken affinity, because water molecules have a strong influence on each other in the confined space between the protein and DNA. We also show here that the two modes of dinucleotide recognition lead to different temperature sensitivity of binding of the same TF to two different motifs. This effect will result in differential gene expression of target genes that contain the entropic and enthalpic sites, enabling organisms to directly sense their body temperature at the level of individual genes. This mechanism is likely to be particularly important in unicellular organisms, plants, and poikilothermic animal species.

**Figure 5.**
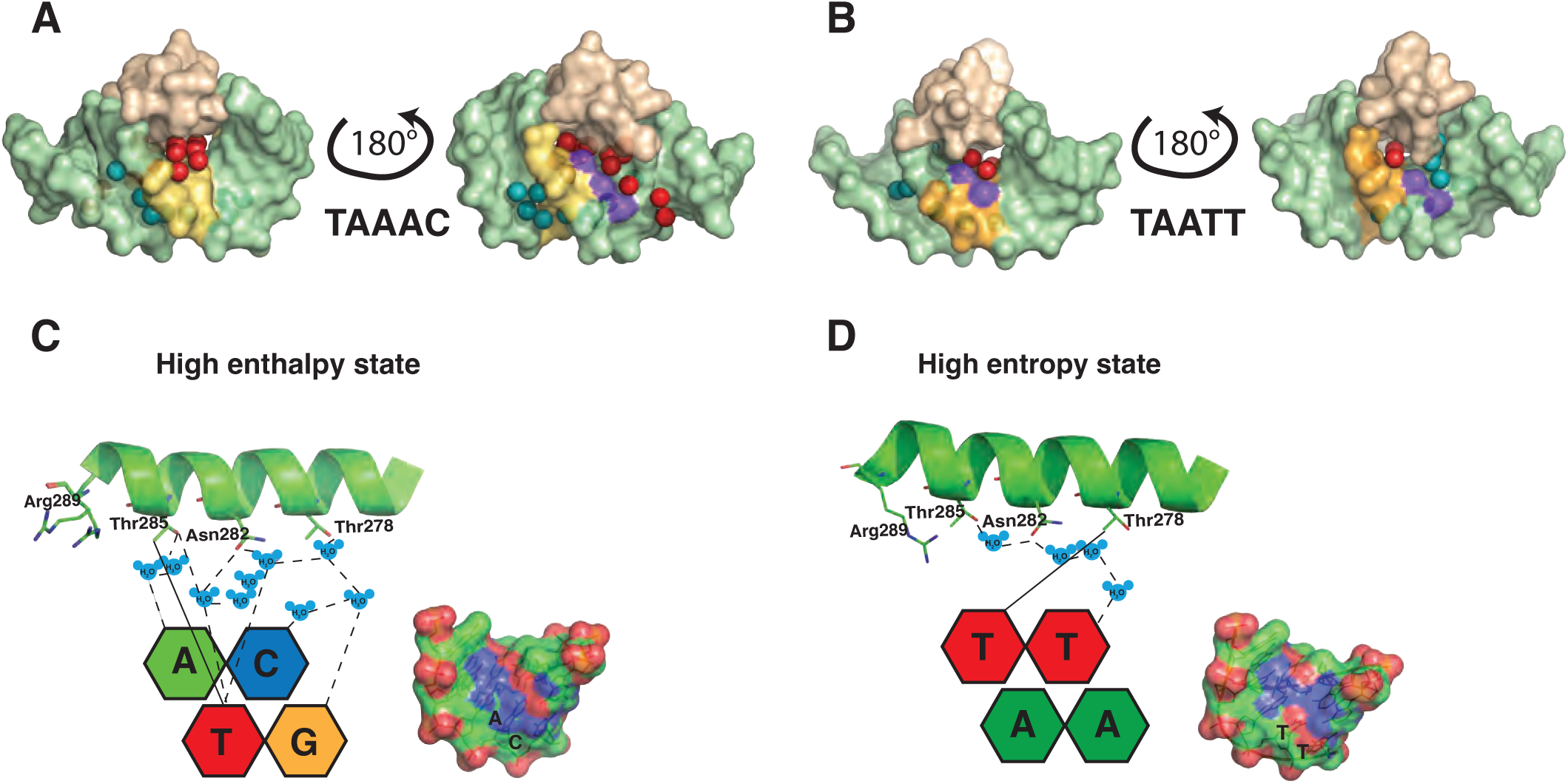
Surface and schematic representations of two binding optima. **a, b**, Surface representation of the protein-DNA interface near the variable bases AC (**a**) and TT (**b**). Only the recognition helix from BARHL2 (wheat) is represented for clarity. The variable bases are in yellow, and the methyl group (CH3) of T is in violet. The red and teal spheres represent the water molecules observed in the interface and minor groove, respectively. **c**, **d**, Schematic representation of the two optimal sites representing enthalpy (**c**) and entropy (**d**) optima. Only the recognition helices and the divergent dinucleotides are shown for clarity. The amino acid residues involved in the recognition are presented as sticks. The bases are color coded as follows: adenine, green; thymine, red; cytosine, blue; guanine, yellow. Note that the high entropy state (TT) has very few fixed water molecules whereas the enthalpic state (AC) contains several properly fixed water molecules which are used for formation of the hydrogen bonds linking BARHL2 to DNA. The small panels of c and d show the charge distribution on the surface of the optimal sequences. Positively charged atoms (nitrogen) are colored blue, negatively charged atoms (oxygen) are colored red, carbon and phosphorus are in green and orange, respectively. Note that TAA**AC** surface shows the distribution of the positive and negative charges as parallel lines that facilitate the ordering of water molecules to high enthalpy interactions, whereas the distribution in TAA**TT** has a “cross” shape favoring entropy and high mobility of waters.

In summary, our work highlights the role of water in molecular recognition, and in local interactions that enable recognition of features that are larger than those that can be bound by an individual chemical bond. Recent advances in prediction of protein structures from sequence ^35,36^ raise the possibility that also a more difficult problem – determining the affinity of macromolecular interactions from sequence – could also be solved using machine learning. However, the non-additive nature of the solvent interactions increases the complexity of the problem, and highlight the need for explicitly considering both enthalpy and entropy in building computational and predictive models for macromolecular recognition.

## Acknowledgements

The authors thank Drs. Inderpreet Sur, Connor Rogerson and Ben Luisi for the critical review of the manuscript, Dr. Lucy Malinina for the fruitful discussion of the DNA structure at high resolution, Dr. Pratyush Das and Karolinska Institutet Protein Science Facility for protein production and Margareta Kling Pilström for technical assistance. This work was supported by the Swedish Research Council, BBSRC/UKRI and Cancer Research UK.

## Author Contributions

E.M. - data curation, formal analysis, investigation, visualization, writing—original draft, writing—review and editing; Y. Y., F. Z., T. X., I. S. - data curation, formal analysis; A. P., C. L. - data curation; J. T., conceptualization, data curation, supervision, project administration, writing—review and editing.

## Accession codes

The atomic coordinates and diffraction data have been deposited to Protein Data Bank with the accession codes 7Z5I and 7Z5K, for MYF5 bound to symmetrical DNA and MYF5 bound to non-symmetrical DNA, respectively and 8PMF for BARHL:DNA^AC^ at the resolution 0.95Å and 8PMN for the BARHL2:DNA^AC^ at the resolution 1.3Å; 8PMC, 8PM5, 8PM7, 8PMV, 8PN4, 8PNA and 8PNC for BARHL2:DNA^TT^, BARHL2:DNA^AT^, BARHL2:DNA^TC^, BARHL2:DNA^TAAGC^, BARHL2:DNA^CC^, BARHL2:DNA^TG^, BARHL2:DNA^GT^, respectively. All sequence reads are deposited to the European Nucleotide Archive under the accession number PRJEB65950.

## Materials and methods

### Protein expression, purification, and crystallization

Expression and purification of the DNA-binding domain fragment of human MYF5 (residues 82-136) as well as BARHL2 (residues 232-292) were performed as described in Refs.^37^ and^22^. The DNA fragments used in crystallization were obtained as single-strand oligonucleotides (Eurofins) and annealed in 20 mM HEPES (pH 7.5) containing 150 mM NaCl and 0.5 mM Tris (2-carboxyethyl) phosphine (TCEP) and 5% glycerol. For each complex, the purified and concentrated protein was first mixed with a solution of annealed DNA duplex at a molar ratio 1:1.2 and after one hour on ice subjected to the crystallization trials. The crystallization conditions for both MYF5 complexes were optimized using an in house developed crystal screening kit combining different PEGs with different additives. Complexes of MYF5 with symmetrical DNA were crystalized in sitting drops by vapor diffusion technique from solution containing 50 mM sodium acetate buffer at pH 4,5, 10% PEG (3350) and 2% 2-Methyl-2,4-pentanediol (MPD). Complexes of MYF5 with non-symmetrical DNA were also crystalized from 0.05M sodium acetate buffer at pH 4.5 but containing 11% of PEG (1000) 2% MPD and 5% PEG (400). All crystals of BARHL2 complexes with different DNAs were obtained in the same conditions from the reservoir solution containing 100 mM Sodium Acetate buffer (pH 4.8), 34% PEG (1000) and 0.06M Sodium Malonate (pH 7.0) All data sets were collected at ESRF from a single crystal on beamline ID23-1, at 100 K using the reservoir solution as cryo-protectant. Data were integrated with the program XDS ^38^ and scaled with SCALA^39^. Statistics of data collection are presented in **Extended Data Table 1**.

### Structure determination and refinement

All structures were solved by molecular replacement using program Phaser as implemented in Phenix^40^ and CCP4^41^ the structure of MYOD (pdb entry 1MDY) as a search model for MYF5 and structure of Drosophila Clawless homeodomain protein (pdb entry 3A01, chain A) as a search model for BARHL2. After the positioning of protein, the density of DNA was clear, and the molecule was built manually using COOT^42^. The rigid body refinement with REFMAC5 was followed by restrain refinement with REFMAC5, as implemented in CCP4^41^ and Phenix.refine^43^. The manual rebuilding of the model was done using COOT. The refinement statistics are presented in **Extended Data Table 1**. In both MYF5 complexes all residues as well as all DNA bases were well defined in the electron density maps. In the structure of MYF5^GT/AC^ only three water molecules were well defined while in the structure of MYF5^AA/AC^ 71 water molecules were traced. All residues and all bases on all BARHL2 complex structures were well visible in the maps. Figures showing structural representations were prepared using PyMOL^44^.

### HT-SELEX and motif analysis<colcnt=4>

MYF6 and BARHL2 HT-SELEX experiments were performed essentially as described in ^22^. The PWM models were generated from cycle of new MYF6 and BARHL2 HT-SELEX reads, using the multinomial (setting = 1) method^45^ with the following seeds: MYF6 single PWM: NRWCAGCTGWYN, AA…TT flank: NAACAGCTGTTN, GT…AC flank: NGTCAGCTGACN; BARHL2 single NSYTAAACGNYN. PWM: NSYTAAWYGNYN, TT: NSYTAATTGNYN, AC: kmer counts were generated using spacek40 (https://github.com/jttoivon/moder2/blob/master/myspacek40.c) for BARHL2 methylated and non-methylated SELEX data.

### Molecular dynamic simulations

All simulations were performed using AMBER 21^46^. For crystal lattice simulations, models of the complete unit cell were created using UCSF Chimera^47^, neutralizing sodium ions added using the AMBER *AddToBox* tool, and then the same tool used to add sufficient additional waters to achieve a final unit cell density of c 1.0. Control simulations of the DNA alone in water were prepared from the crystal structure coordinates using the Amber *tleap* tool, adding neutralising sodium ions and enough waters to fill a truncated octahedral periodic box extending a minimum of 10 Å beyond any solute atom. Systems were parameterised using the FF14SB force field^48^ for the protein component, with additional BSC1 parameters for the DNA^49^. Waters and ions were modelled using the TIP3P and Joung-Cheatham^50^ parameters respectively.

Simulations were performed using *pmemd.cuda*. Unless specified otherwise, all simulations used default parameters. Systems were first energy minimised with restraints (0.1 kcal mol^-1^ Å^-2^) on all atoms except modelled waters (i.e., waters observed in the crystal structures were in the restrained group), before a second energy minimisation step without restraints. Molecular dynamics simulations used Langevin dynamics with a collision frequency of 5 ps^-1^ and a 2 fs timestep, with SHAKE on all bonds to hydrogen atoms. Non-bonded interactions were handled using the Particle Mesh Ewald method, with a direct space cutoff of 9 Å. Snapshots were saved every 100 ps. For the crystal lattice simulations, MD began with a 10ns NVT simulation, with a target temperature of 300K, and restraints of 0.01 kcal mol^-1^ Å^-2^ on all solute atoms, followed by a 50 ns NVT simulation in which only DNA atoms were restrained (same force constant). For the 50 ns production NVT simulation only DNA C1’ atoms remained restrained. For reference simulations of the DNA alone in solution, MD began with a 50ns NPT simulation at 300K (pressure coupling parameter 2 ps^-1^) with restraints (0.01 kcal mol^-1^ Å^-2^), followed by a 200 ns NPT production simulation in which the only restraints were on the Watson-Crick hydrogen bonds in the terminal base pairs (a flat-bottomed potential that was zero between 1.9 and 2.1 Å, and had a force constant of 10 kcal mol^-1^ Å^-2^ outside this range).

DNA helical parameters were analysed using the AMBER21 *cpptraj* tool^51^. Other analysis and visualisation were performed in Jupyter notebooks, using the *MDTraj*^52^ and *Matplotlib* packages. The method to analyse water molecule mobility involves first permuting the indices of the water oxygen atoms in each frame in the trajectory so that each remains within a small region of space. The mean position and fluctuation of each water can then be measured. The re-indexing is an iterative process using the linear sum assignment approach, the Python code to implement this is available in the supporting information.

### Mutational analysis

The pETG20A_SPB vectors with BARHL2 and its mutant sequences (GenScript) were expressed in Rosetta(DE3)pLysS *E.coli* strain (Millipore). Briefly, the bacteria were grown overnight at 37 ^0^C in LB Broth medium (Gibco) with carbenicillin (0.1 mg/ml) and chloramphenicol (34 ug/ml), and then transferred to the induction medium consisting of the previous one with additional reagents with the following final concentrations: 1 mM MgSO_4_, metal mixture (50 µM FeCl_3_, 20 µM CaCl_3_, 10 µM each of MgCl_2_, and ZnSO_4_, 2 µM each of CoCl_2_, CuCl_2_, NiCl_2_, Na_2_MoO_4_, Na_2_SeO_3_, H_3_BO_3_), 60 µM HCl, NPS (50 mM KH_2_PO_4_, 50 mM Na_2_HPO_4_, 25 mM (NH_4_)_2_SO_4_) and 5052 (0.5% glycerol, 0.05% glucose, 0.2% alpha-lactose)^53^. The cells were then incubated for 8h at 37 °C, followed by 40h at 17 °C. Then, cells were collected by centrifugation and lysed by shaking for 20 minutes in 100 µl of lysis buffer (0.5 mg/ml Lysozyme, 1 mM PMSF, 400 mM NaCl, 100 mM KCl, 10% glycerol, 0.5% Triton X-100, 10 mM imidazole in 50 mM potassium phosphate buffer, pH 7.8), and stored overnight at −80 °C. After thawing, the lysate was incubated for 45 minutes with 20 µl of Ni Sepharose 6 Fast Flow beads (GE Healthcare), washed and resuspended in 100 µl of buffer A (30 mM NaCl, 10 mM imidazole in 50 mM Tris-HCl, pH 7.5). For digestion of bacterial DNA, DNAse I and MgSO_4_ were added to 10 ug/ml and 15 mM final concentration, respectively. After incubation for 45 minutes at room temperature, the beads were washed two times with 600 µl of buffer A and 2 times with 600 µl of buffer A containing 50 mM imidazole. The proteins were eluted in 100 µl of buffer A with 500 mM imidazole. Protein concentration was then measured using a Bradford assay (#B6916, Sigma), after which the proteins were diluted in Promega buffer (50 mM NaCl, 10 mM MgCl_2_, 4% glycerol in 10 mM Tris-HCl, pH 7.5) to 30-100 ug/ml concentration.

SELEX experiments were performed as described in^54^. Briefly, 100-250 ng of proteins were mixed with 200-500 ng of DNA ligands and incubated at room temperature for 20 minutes, the incubation buffer contained 51.4 mM NaCl, 1.4 mM MgCl_2_, 4% glycerol, 100 µM EGTA, 0.7 mM 1,4-dithiothreitol, 3.7 µg/ml poly-dI-dC, 1.4 uM ZnSO_4_ in 10.4 mM Tris-HCl, pH 7.5 (SELEX buffer). Magnetic Ni-Sepharose beads (GE Healthcare) were washed in Promega buffer containing 0.2% BSA, and suspended in 25 µl of SELEX buffer. For each protein-DNA mixture, 1.75 µl of the bead suspension was added, followed by incubation for 40 minutes at room temperature. The beads were then washed in wash buffer (5 mM EDTA, 1 mM 1,4-dithiothreitol in 5 mM Tris-HCl, pH 7.5) using Tecan HydroSpeed plate washer, followed by suspension of the beads to 35 µl of Elution buffer (1 mM MgCl_2_, 0.1% Tween 20 in 10 mM Tris-HCL, pH 7.8) followed by incubation for 10 minutes at 80 °C. To ensure that the ligands were double stranded, the eluted DNA ligands were amplified twice. First, the products were amplified to completion (33 PCR cycles for first SELEX cycle and 26 for all subsequent cycles). Then, the PCR products were diluted 10-fold and amplified for two PCR cycles to ensure that single-stranded DNA and annealed products containing mismatched random sequences were fully converted to double-stranded form. The PCR reaction was performed with Phusion Hot Start II DNA Polymerase (Thermo Scientific). The DNA concentration was controlled by qPCR with 1x SYBR™ Green I Nucleic Acid Gel Stain (Invitrogen) using LightCycler-480 Instrument II (Roche). Amplified DNA ligands were used for incubation with proteins in the next SELEX cycle. A total of four SELEX cycles were performed. The DNA ligands from 0, 3rd and 4th SELEX cycles were sequenced (Illumina HiSeq 2000) and analyzed as described in Ref.^21^.

### Temperature HT-SELEX

To examine how DNA-binding specificity of BARHL2 changes with temperature, HT-SELEX of BARHL2 was performed under five different temperatures of 0°C, 10°C, 20°C, 30°C, 40°C. For Temperature HT-SELEX, the DNA ligands was designed according to Illumina’s Truseq library (**Extended Data Table 2**, 101N SELEX Ligand) and synthesized from IDT as Ultramer DNA oligos. The oligos contain a 101-bp region with randomized nucleotides, flanked by adapters of fixed sequences for amplification. First, double-stranded ligands (the input of SELEX) were synthesized from the oligos by PCR amplification with primers that match to the adapters (**Extended Data Table 2**, PCR primers). The PCR primers were also used to amplify the library between SELEX cycles. Before sequencing, the ligands were further amplified with primers (**Extended Data Table 2**, PE primers) containing multiplexing indices and sequences of the Illumina flow cell (P5 or P7).

HT-SELEX was performed in microplates according to the previous protocol^7,55^. First, 100–200 ng double-stranded DNA ligand was mixed with 20-200 ng purified his-tagged TFs in 20 µl volume of the incubation buffer (140 mM KCl, 5 mM NaCl, 2 mM MgSO_4_, 3 μM ZnSO_4_, 100 μM EGTA, 1 mM K_2_HPO_4_, 20 mM HEPES, pH 7.0). The mixture was incubated for 20 min on a PCR incubator (Bio-Rad S1000 Thermal Cycler) for temperature control. Then, 1.8 µl of nickel magnetic Sepharose beads (28-9799-17, GE Healthcare; pre-blocked with 25 mM Tris-HCL, 0.5% BSA, 0.1% Tween 20, 0.02% NaN_3_) was added into the mixture to pull down the TFs and their associated DNA ligands. After mixing at 1900 rpm with a microplate shaker (13500-890, VWR), the plates were washed 15 times on a microplate washer (Tecan Hydrospeed) with the washing buffer (10 mM EDTA in 5 mM Tris-HCl pH 7.5, kept under 4°C before use). Suspension of the washed beads was then PCR amplified to produce DNA ligands for the next SELEX cycle. After repeating SELEX for four cycles, the ligands from each cycle and the input were sequenced and analyzed.

