## Supplementary Figures and tables for "Interfacial water confers transcription factors with dinucleotide specificity"

### Extended Data Fig. 1

**a**

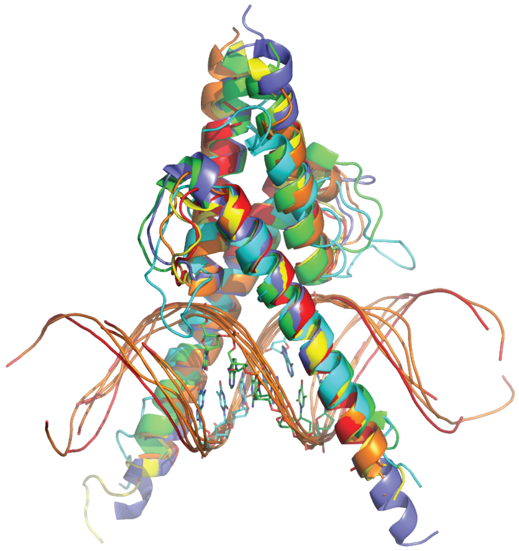

MYF5/USF/MYOD/SCL/E47/CLOCK/BMAL1/TCF4

**b**

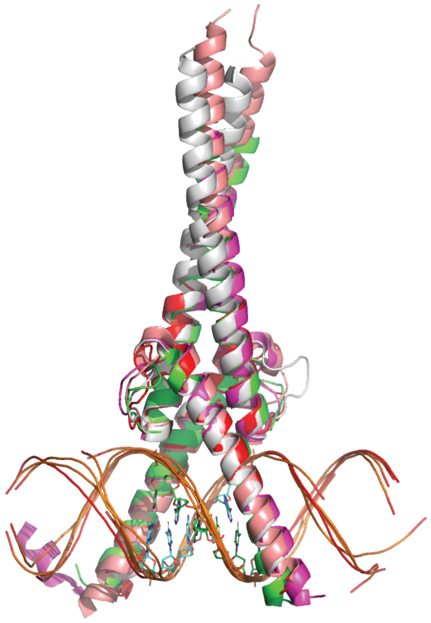

MYF5/SREBP/MAX/MYC/MAD

**c**

```
                    80        90        100       110               120       130
MYF5   ENLYFQS------MDRRKAATMRERRRLKKVNQAFETLKRCT-TTN-P--NQRLPKVEILRNAIRYIESLQE---------------------------
MYF6   ENLYFQSKRKSAPTDRRKAATLRERRRLKKINEAFEALKRRT-VAN-P--NQRLPKVEILRSAISYIERLQD---------------------------
USF    -----------MDEKRRAQHNEVERRRRDKINNWIVQLSKIIPDSSMESTKSGQSKGGILSKASDYIQELRQSNHR-----------------------
MYOD   ----MELKRKTTNADRRKAATMRERRRLSKVNEAFETLKRST-SSN-P--NQRLPKVEILRNAIRYIEGLQALLRD-----------------------
SCL    SQDPEISDGPHTKVVRRIFTNSRERWRQQNVNGAFAELRKLIPTHP-P--DKKLSKNEILRLAMKYINFLAKLLND---QEEEGTQR------------
E47    MADLSLEEKDLRDRERRMANNARERVRVRDINEAFRELGRMCQMHLKS--DKAQTKLLILQQAVQVILGLEQQVRE---RNLNPKAA------------
CLOCK  --------MDDKDKAKRVSRNKSEKKRRDQFNVLIKELGSMLPGN-----ARKMDKSTVLQKSIDFLRKHKEITAW-----------------------
BMAL1  -------MHQGRIKNAREAHSQIEKRRRDKMNSFIDELASLVPTCNAM--SRKLDKLTVLRMAVQHMKTLRG--AW-----------------------
TCF4   -------------HMRRMANNARERLRVRDINEAFKELGRMVQLHLKS--DKPQTKLLILHQAVAVILSLEQQVRE---R-------------------
SREBP  ----------QSRGEKRTAHNAIEKRYRSSINDKIIELKDLVVGT-----EAKLNKSAVLRKAIDYIRFLQHSNQKLKQENLSLRTAV-HKSKSLKDL-
MAX    ---------------KRAHHNALERKRRDHIKDSFHSLRDSVPSLQGE----KASRAQILDKATEYIQYMRRKNHTHQQDIDDLKRQNALLEQQV----
MYC    ----------GHMNVKRRTHNVLERQRRNELKRSFFALRDQIPELENN---EKAPKVVIL TKAKLHIKKLEDSDRKAVHQIDQLQREQRHLKRQLEKL
MAD    ---------------SRSTHNEMEKNRRAHLRLSLEKLKGLVPLGPDS---SRHTTLSLL TKAKLHIKKLEDSDRKAVHQIDQLQREQRHLKRQLEKL
```

**d**

| | |
|---|---|
| MYF5(this study) | GCGCAACAGCTGACGCGG |
| MYF5(this study) | GCGCGTCAGCTGACGCGG |
| USF(1AN4) | CACCCGGTCACGTGGCCTACA |
| MYOD(1MDY) | CACCCGGTCACGTGGCCTACA |
| SCL/E47(2YPB) | ACCATCTGTTC |
| CLOCK/BMAL(4H10) | AGGAACACGTGACCC |
| TCF4(6OD3) | CATACACGTGTAT |
| SREBP(1AM9) | CATGAGATCACCCCACTGCAA |
| MAX/MAD(1NLW) | GAGTAGCACGTGCTACTC |
| MAX/MYC(1NKP) | CGAGTAGCACGTGCTACTC |

**e**

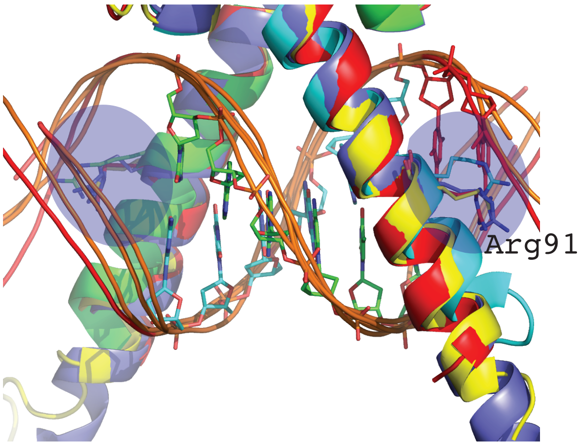


| | 91 |
|---|---|
| MYF5 | MRERRRL |
| MYOD | MRERRRR |
| SCL | SRERRRL |
| E47 | ARERVRV |
| TCF4 | ARERLRV |

**Extended Data Fig. 1 Comparison of the structure of MYF5 solved in this study with different representatives of bHLH (basic helix-loop-helix) and bZIP (basic helix-loop-helix-zipper) families.** (**a**) Structural alignment of DBD of MYF5 homodimer (red) with USF homodimer (cyan, 1AN4.pdb), MYOD homodimer (yellow, 1MDY.pdb), SCL/E47 complex (light blue, 2YPB.pdb), CLOCK/BMAL1 complex (orange, 4H10.pdb) and TCF4 homodimer (green, 6OD3.pdb). (**b**) Structural alignment of MYF5 homodimer with SREBP1 homodimer (light green, 1AM9.pdb), MAX homodimer (magenta, 1HLO.pdb), MYC/MAX complex (salmon, 1NKP.pdb) and MAD/MAX complex (gray, 1NLW.pdb). The proteins used for structural alignments in **a** and **b** are listed under the figure. The DNA bases involved in the binding are presented as sticks. (**c**) Sequence alignment of the DBD of proteins, used for the structural alignment in **a** and **b** with MYF6 in addition**.** The numbering on the top corresponds to MYF5 numbering. The helixes observed in the MYF5 structure are highlighted in light yellow. The light green box underlines the binding site. The color code for the amino acids is kept with the respect of Clustal Omega server used for the sequence alignment (https://www.ebi.ac.uk/Tools/msa/clustalo/). Hydrophobic amino acids are colored red, polar – green, negatively charged - blue and positively charged – magenta. (**d**) The DNA sequences used in the structural studies. The binding sites are highlighted with the light green box. (**e**) Close view of the alignment of MYF5/MYOD/SCL/E47/TCF4 structures which are known to recognize the flanking dinucleotides. The sequence alignment shows that those bHLH contain arginine residue (Arg91 in MYF5) in the specific position which is responsible for that recognition, conversely the MAX/MYC/MAD/CLOCK/BMAL1/USF/SREBP structures and sequences that arginine is replaced by different residues which are not suitable for the flank recognition.

**Estended Data Fig. 2**

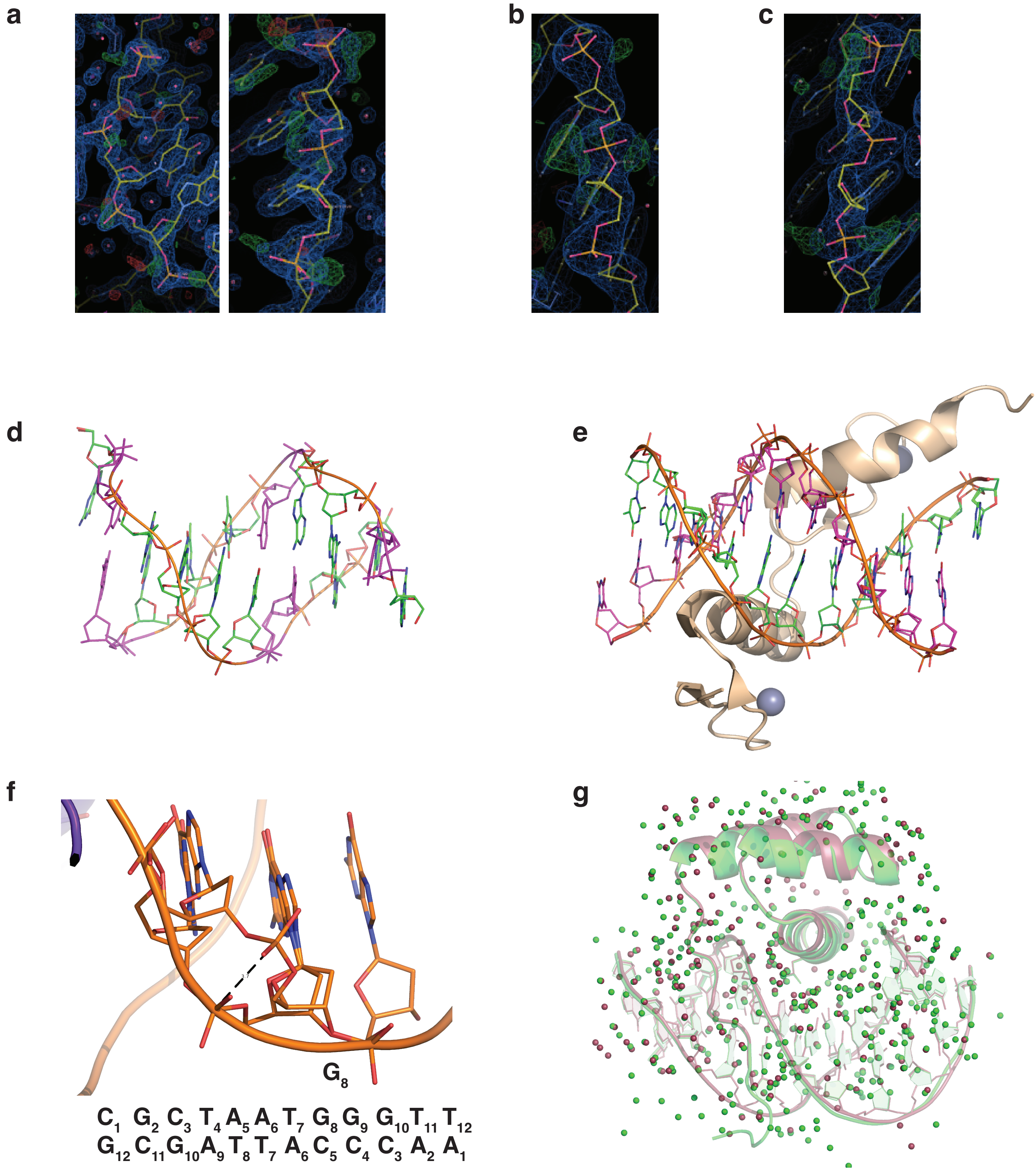

**Extended Data Fig. 2 High resolution structures of BARHL2 showed double conformations of the DNA.**

**(a)** Electron density map of the backbone phosphates in the structure with DNA TAAAC at 1.3Å resolution. The green “balls” are the possible alternative positions of the oxygen atoms of phosphates. (**b, c)** Electron density map observed for the BARHL2- $DNA^{GC}$ ($G_{12}C_{11}G_{10}$) and BARHL2- $DNA^{CC}$ ($G_5C_4C_3$), respectively. (**d**) The structure of the nonamer (GCGAATTCG) solved at 0.89 Å resolution showed the similar double conformation feature of the backbone phosphates (Soler-Lopez et al., 2000). The figure is created from the coordinates of 1ENN.pdb file. The backbone phosphates observed in the double conformations are colored in magenta. (**e**) An example of the alternative conformation observed in the structure of protein-DNA complex. The figure is created from the coordinates of mouse Znf57-DNA complex obtained from 4GZN.pdb file. The protein moiety is colored in beige, two Zn atoms are presented in gray balls, the different DNA strands are colored in magenta and orange. Note that the most phosphates on the top strand show double conformation (Liu et.al., 2012). (**f**) Double conformation of phosphates for $\mathbf{G_8G_9}$ observed in complex BARHL2- $DNA^{\mathbf{TG}}$. The distance between two phosphorous atoms is 3.7Å. (**g**) Structural alignment of two structures of BARHL2:$DNA^{AC}$ at the resolutions 1.3Å (raspberry) and 0.95Å (green). The water molecules represented as small spheres and coloured respectively to the complex. Note that the water molecules in the interface are very well conserved.

### Extended Data Fig. 3

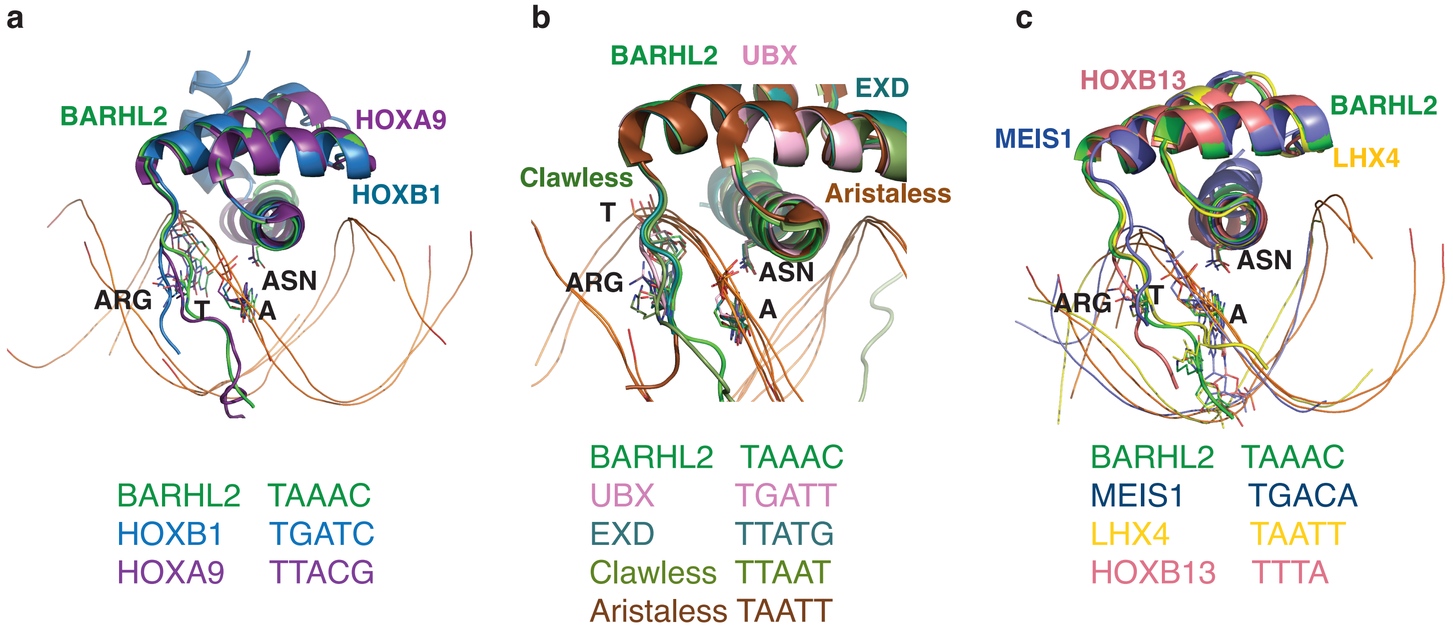


**d**

```
                                                                278  282 285  289
BARHL2       RAKKPRKARTAFSDHQLNQLERSFERQKYLSVQDRMDLAAALNLTDTQVKTWYQNRRTKWKRQTAVGLELLAE
HOXB1        GLGSPSGLRTNFTTRQLTELEKEFHFNKYLSRARRVEIAATLELNETQVKIWFQNRRMKQKKREREGG-----
HOXA9        HARSTRKKRCPYTKHQTLELEKEFLFNMYLTRDRRYEVARLLNLTERQVKIWFQNRRMKMKKINKD-------

BARHL2       RAKKPRKARTAFSDHQLNQL---ERSFERQKYLSVQDRMDLAAALNLTDTQVKTWYQNRRTKWKRQTAVGLELLAE
UBX          TNGLRRRGRQTYTRYQTLEL---EKEFHTNHYLTRRRRIEMAHALSLTERQIKIWFQNRRMKLKKEIQA-------
EDX          ----ARRKRRNFSKQASEILNEYFYSHLSNPYPSEEAKEELARKCGITVSQVSNWFGNKRIRYKKNI---------
Clawless     -MRTPPKRKKPRTSFTRIQVAELEKRFHKQKYLASAERAALARGLKMTDAQVKTWFQNRRTKWRRQTAEEREAERQ
Aristaless   --APKRKQRRYRTTFTSFQLEELEKAFSRTHYPDVFTREELAMKIGLTEARIQVWFQNRRAKWRKQEKV-------

BARHL2       RAKKPRKARTAFSDHQLNQLERSFERQKYLSVQDRMDLAAALNLTDTQVKTWYQNRRTKWKRQTAVGLELLAE
MEIS1        --AFPKVATNI-----MRAWLFQHLTHPYPSEEQKKQLAQDTGLTILQVNNWFINARRRIVQPM---------
LHX4         GA---KRPRTTITAKQLETLKNAYKNSPKPARHVREQLSSETGLDMRVVQVWFQNRRAKEKRLK---------
HOXB13       -----RKKRIPYSKGQLRELEREYAANKFITKDKRRKISAATSLSERQITIWFQNRRVKEKKVLAKVK-----
```

**Extended data Fig. 3 Comparison of BARHL2-DNA$^{AC}$ structure with the representatives of different homeobox TF families.**

Structural alignment of BARHL2-DNA$^{AC}$ complex (green) with the complexes of anterior HOXB1-DNA$^{TGAT}$ (blue) and posterior HOXA9-DNA$^{TTAC}$ (violet) (**a**). Structural alignment of BARHL2-DNA$^{AC}$ complex (green) with UBX-DNA$^{AT}$ (pink), EXD-DNA$^{AT}$ (dark green), Clawless-DNA$^{TAAT}$ (light green) and Aristaless-DNA$^{TAAT}$ (brown) (**b**). Structural alignment of BARHL2-DNA$^{AC}$ complex (green) with LHX4-DNA$^{AT}$ (yellow), MEIS1-DNA$^{TGAC}$ (dark blue) and HOXB13-DNA$^{TTATT}$ (salmon) (**c**) (All structures are solved in Taipale lab). The key residues ASN and ARG as well as the key bases T and A involved into binding are labeled. (**d**) Sequence alignment of the structures presented in (**a-c**). The recognition helix is highlighted in yellow, the recognizing residues colored red. The numbering on the top of the sequence is corresponding to BARHL2 numbering. (**e**) The DNA-binding domain sequences of fourteen mutants designed to test the role of the threonine (Thr278 and Thr285) in the DNA sequence recognition. In the BARHL2 DNA-binding domain sequence we mutated the Thr278 and Thr285 to the corresponding residues of 14 different homeobox proteins. Thr278 and Thr285 are colored green in the original BARHL2 sequence (on the top) and in the mutated sequences if they belong to the original sequence of other homeobox protein. The mutations are colored red.

### Extended Data Table 1. X-ray data collection and refinement statistics.

| Protein/core | MYF5/CAGCTG | | BARHL2/TAA | | | | | | | | |
|---|---|---|---|---|---|---|---|---|---|---|---|
| Flanking DNA | AC/GT | AA/GT | HRAC | AC | TT | AT | TC | GC | CC | TG | GT |
| Resolution range (Å) * | 41.67-3.0 (3.10-3.0) | 46.00 - 2.28 (2.36 - 2.28) | 28.78 - 0.95 (0.98 -0.95) | 29.89 - 1.30 (1.35 -1.30) | 39.49 - 1.85 (1.91 -1.85) | 46.91 - 2.4 (2.48 - 2.4) | 39.55 - 1.7 (1.76 - 1.7) | 39.04 - 2.1 (2.17 - 2.1) | 39.37 - 2.6 (2.69 - 2.6) | 35.88 - 1.45 (1.50 - 1.45) | 39.43 - 2.05 (2.12 - 2.05) |
| Space group | C 2 | C 2 | P $2_12_12_1$ | P $2_12_12_1$ | P $2_1$ | P $2_1$ | P $2_1$ | P $2_1$ | P $2_1$ | P $2_12_12_1$ | P $2_1$ |
| Unit cell (Å, º) | 166.7 33.9 53.7 90 91.04 90 | 170.4 33.8 53.6 90 91.9 90 | 38.8 47.3 72.5 90 90 90 | 38.6 47.1 72.3 90 90 90 | 39.6 123.1 62.2 90 94.2 90 | 39.0 134.3 65.6 90 92.6 90 | 39.6 123.8 62.7 90 93.5 90 | 39.1 133.7 64.9 90 92.7 90 | 39.4 134.5 64.5 90 93.2 90 | 38.8 46.8 71.8 90 90 90 | 39.6 123.7 62.4 90 93.9 90 |
| Unique reflections | 5660 (589) | 14126 (1362) | 83984 (8167) | 32862 (3157) | 50254 (4922) | 26180 (2627) | 65754 (6406) | 38783 (3913) | 20528 (1981) | 23749 (2210 | 37338 (3673) |
| Multiplicity | 2.8 (2.7) | 5.0 (5.2) | 6.5 (6.0) | 7.1 (6.6) | 3.8 (3.7) | 7.2 (6.6) | 3.8 (2.9) | 5.7 (5.7) | 3.8 (4.0) | 7.9 (3.3) | 5.2 (4.8) |
| Completeness (%) | 90.1 (93.2) | 94.1 (97.2) | 99.5 (97.9) | 99.5 (97.2) | 99.3 (97.7) | 99.03 (100) | 99.3 (97.2) | 99.6 (99.8) | 99.3 (96.9) | 99.2 (94.4) | 99.6 (98.7) |
| Mean I/sigma(I) | 8.46 (1.08) | 17.5 (1.8) | 15.3 (0.5) | 10.2 (0.7) | 8.5 (0.4) | 8.8 (0.7) | 9.5 (0.5) | 7.0 (0.3) | 5.9 (1.0) | 13.6 (0.6) | 7.7 (0.8) |
| Wilson B-factor | 86.17 | 58.00 | 10.84 | 18.84 | 35.24 | 47.10 | 32.02 | 51.27 | 51.48 | 18.23 | 35.58 |
| R-merge | 0.063 (1.062) | 0.024 (0.72) | 0.045 (2.89) | 0.073 (1.96) | 0.071 (2.70) | 0.145 (2.71) | 0.050 (1.77) | 0.11 (4.61) | 0.19 (1.54) | 0.077 (1.41) | 0.153 (2.00) |
| R-meas | 0.07799 (1.293) | 0.028 (0.83) | 0.053 (3.54) | 0.084 (2.33) | 0.093 (3.51) | 0.167 (3.22) | 0.066 (2.57) | 0.132 (5.6) | 0.263 (2.11) | 0.086 (1.91) | 0.19 (2.6) |
| R-pim | 0.04471 (0.7246) | 0.014 (0.41) | 0.021 (1.42) | 0.032 (0.88) | 0.047 (1.77) | 0.063 (1.23) | 0.034 (1.41) | 0.054 (2.3) | 0.135 (1.05) | 0.029 (0.96) | 0.083 (1.17) |
| CC1/2 | 0.999 (0.616) | 1.000 (0.89) | 0.998 (0.17) | 0.998 (0.26) | 0.998 (0.15) | 0.996 (0.18) | 0.999 (0.29) | 0.999 (0.08) | 0.985 (0.28) | 0.999 (0.28) | 0.993 (0.18) |
| Reflections used in refinement | 5673 (589) | 14060 (1359) | 83944 (8167) | 32821 (3156) | 50206 (4922) | 26124 (2627) | 65719 (6405) | 38705 (3914) | 20488 (1981) | 23721 (2209) | 37295 (3673) |
| R-work | 0.23 (0.46) | 0.24 (0.37) | 0.14 (0.38) | 0.15 (0.39) | 0.23 (0.43) | 0.23 (0.42) | 0.22 (0.44) | 0.24 (0.44) | 0.24 (0.38) | 0.18 (0.37) | 0.23 (0.38) |
| R-free | 0.28 (0.53) | 0.22 (0.37) | 0.17 (0.38) | 0.2 (0.4) | 0.26 (0.4) | 0.28 (0.45) | 0.25 (0.4) | 0.27 (0.46) | 0.27 (0.37) | 0.22 (0.33) | 0.26 (0.41) |
| Number of non-hydrogen atoms | 1683 | 1779 | 1937 | 1307 | 4384 | 4884 | 4472 | 4148 | 4572 | 1362 | 4427 |
| macromolecules | 1680 | 1701 | 1507 | 1041 | 4053 | 4557 | 4035 | 4027 | 4398 | 1128 | 4076 |
| solvent | 3 | 78 | 430 | 266 | 331 | 327 | 437 | 121 | 174 | 234 | 351 |
| Protein residues | 112 | 112 | 66 | 63 | 241 | 301 | 239 | 243 | 285 | 67 | 243 |
| RMS(bonds) | 0.016 | 0.005 | 0.018 | 0.015 | 0.013 | 0.014 | 0.012 | 0.015 | 0.015 | 0.016 | 0.016 |
| RMS(angles) | 1.66 | 1.11 | 2.43 | 1.92 | 1.69 | 1.78 | 1.71 | 2.28 | 2.17 | 2.20 | 2.35 |
| Ramachandran favoured (%) | 96.30 | 96.30 | 100.0 | 98.36 | 99.57 | 97.92 | 100.0 | 99.15 | 96.36 | 100.0 | 100.0 |

| | | | | | | | | | | |
|---|---|---|---|---|---|---|---|---|---|---|
| Ramachandran allowed (%) | 2.78 | 2.78 | 0.00 | 1.64 | 0.43 | 1.38 | 0.00 | 0.85 | 2.18 | 0.00 | 0.00 |
| Ramachandran outliers (%) | 0.93 | 0.93 | 0.00 | 0.00 | 0.00 | 0.69 | 0.00 | 0.00 | 1.45 | 0.00 | 0.00 |
| Average B-factor | 106.02 | 84.13 | 17.59 | 24.03 | 45.18 | 48.82 | 42.33 | 65.87 | 56.94 | 23.28 | 43.38 |
| macromolecules | 106.05 | 84.40 | 15.60 | 21.27 | 45.15 | 49.19 | 42.08 | 67.67 | 58.88 | 23.28 | 41.14 |
| solvent | 89.90 | 78.12 | 33.43 | 34.05 | 45.55 | 43.73 | 44.67 | 59.32 | 40.65 | 31.60 | 43.43 |

* Statistics for the highest-resolution shell are shown in parentheses.

**Extended Data Table 2 - Sequence information for DNA ligands**

This table contains the infomation of all oligos used in NCAP-SELEX and free-DNA SELEX

The PCR_primers are used to amplify lig147 and lig200 between cycles, and to synthesize dou

Before sequencing, all libraries are amplified using the PE_PCR_primers to add barcode and n

| | |
|---|---|
| **Lig147** | CCCTACACGACGCTCTTCCGATCTNNNNNNNNNNNNNNNNNNNNNN |
| **Lig200** | CCCTACACGACGCTCTTCCGATCTNNNNNNNNNNNNNNNNNNNNNN |
| **Lig70Nlinker (the control ligand that positions nucleosome at the center with 127 bp of wid** | |
| | CCCTACACGACGCTCTTCCGATCTNNNNNNNNNNNNNNNNNNNNNN |
| **PCR_primer (forward)** | CCCTACACGACGCTCTTCC |
| **PCR_primer (reverse)** | CAGACGTGTGCTCTTCCG |
| **PE_PCR_primer (forwar** | AATGATACGGCGACCACCGAGATCTACACTCTTTCCCTACACGACG |
| **PE_PCR_primer (reverse, with bases corresponding to the barcodes in lower case)** | |

| **Barcode** | **Sequence** |
|---|---|
| CCCGCGCT | CAAGCAGAAGACGGCATACGAGATagcgcgggGTGACTGGAGTTCA |
| TCAATACC | CAAGCAGAAGACGGCATACGAGATggtattgaGTGACTGGAGTTCAG |
| TTAGATAA | CAAGCAGAAGACGGCATACGAGATttatctaaGTGACTGGAGTTCAG |
| ATGTATCA | CAAGCAGAAGACGGCATACGAGATtgatacatGTGACTGGAGTTCAG |
| ACTTCTGT | CAAGCAGAAGACGGCATACGAGATacagaagtGTGACTGGAGTTCA |
| CCAGTAAG | CAAGCAGAAGACGGCATACGAGATcttactggGTGACTGGAGTTCAG |
| AGTGGGTA | CAAGCAGAAGACGGCATACGAGATtacccactGTGACTGGAGTTCAG |
| GACCCGGT | CAAGCAGAAGACGGCATACGAGATaccgggtcGTGACTGGAGTTCA |
| TCGAAGAT | CAAGCAGAAGACGGCATACGAGATatcttcgaGTGACTGGAGTTCAG |
| GCTCCGAT | CAAGCAGAAGACGGCATACGAGATatcggagcGTGACTGGAGTTCA |
| AAAAAACG | CAAGCAGAAGACGGCATACGAGATcgttttttGTGACTGGAGTTCAGA |
| GTGGATTC | CAAGCAGAAGACGGCATACGAGATgaatccacGTGACTGGAGTTCA |
| TGGAGGTG | CAAGCAGAAGACGGCATACGAGATcacctccaGTGACTGGAGTTCAG |
| AGCTCAGA | CAAGCAGAAGACGGCATACGAGATtctgagctGTGACTGGAGTTCAG |
| TTCAACGC | CAAGCAGAAGACGGCATACGAGATgcgttgaaGTGACTGGAGTTCA |
| GGGCTGCA | CAAGCAGAAGACGGCATACGAGATtgcagcccGTGACTGGAGTTCA |
| GCCGAATT | CAAGCAGAAGACGGCATACGAGATaattcggcGTGACTGGAGTTCAG |
| TAGTTGAT | CAAGCAGAAGACGGCATACGAGATatcaactaGTGACTGGAGTTCAG |
| AATGCCTA | CAAGCAGAAGACGGCATACGAGATtaggcattGTGACTGGAGTTCAG |
| CGCGAAGT | CAAGCAGAAGACGGCATACGAGATacttcgcgGTGACTGGAGTTCAG |
| AACATAAC | CAAGCAGAAGACGGCATACGAGATgttatgttGTGACTGGAGTTCAGA |
| CCCCAGAT | CAAGCAGAAGACGGCATACGAGATatctggggGTGACTGGAGTTCA |
| ATCGAATG | CAAGCAGAAGACGGCATACGAGATcattcgatGTGACTGGAGTTCAG |
| GAAACGTG | CAAGCAGAAGACGGCATACGAGATcacgtttcGTGACTGGAGTTCAG |
| CCAATCTT | CAAGCAGAAGACGGCATACGAGATaagattggGTGACTGGAGTTCA |
| CACCGACC | CAAGCAGAAGACGGCATACGAGATggtcggtgGTGACTGGAGTTCA |
| ATTGCCCT | CAAGCAGAAGACGGCATACGAGATagggcaatGTGACTGGAGTTCA |
| TTGGTGGT | CAAGCAGAAGACGGCATACGAGATaccaccaaGTGACTGGAGTTCA |
| TGATCCTA | CAAGCAGAAGACGGCATACGAGATtaggatcaGTGACTGGAGTTCAG |
| AGTTAAAA | CAAGCAGAAGACGGCATACGAGATttttaactGTGACTGGAGTTCAGA |
| CCAAATAA | CAAGCAGAAGACGGCATACGAGATttatttggGTGACTGGAGTTCAGA |
| TACCTCAC | CAAGCAGAAGACGGCATACGAGATgtgaggtaGTGACTGGAGTTCA |
| CATGATAT | CAAGCAGAAGACGGCATACGAGATatatcatgGTGACTGGAGTTCAG |
| CCTCACCT | CAAGCAGAAGACGGCATACGAGATaggtgaggGTGACTGGAGTTCA |
| GCACCAGT | CAAGCAGAAGACGGCATACGAGATactggtgcGTGACTGGAGTTCAG |
| TGCTCCGT | CAAGCAGAAGACGGCATACGAGATacggagcaGTGACTGGAGTTCA |
| CCTCGTTG | CAAGCAGAAGACGGCATACGAGATcaacgaggGTGACTGGAGTTCA |
| CTCGCTAT | CAAGCAGAAGACGGCATACGAGATatagcgagGTGACTGGAGTTCA |
| GACTCTCT | CAAGCAGAAGACGGCATACGAGATagagagtcGTGACTGGAGTTCA |

| | |
|---|---|
| TAAGTTAT | CAAGCAGAAGACGGCATACGAGATataacttaGTGACTGGAGTTCAG |
| GCGGGTGT | CAAGCAGAAGACGGCATACGAGATacacccgcGTGACTGGAGTTCA |
| TCTGCCGA | CAAGCAGAAGACGGCATACGAGATtcggcagaGTGACTGGAGTTCA |
| GGCGCGGA | CAAGCAGAAGACGGCATACGAGATtccgcgccGTGACTGGAGTTCAG |
| CGTTCTTT | CAAGCAGAAGACGGCATACGAGATaaagaacgGTGACTGGAGTTCA |
| ATCGAGAC | CAAGCAGAAGACGGCATACGAGATgtctcgatGTGACTGGAGTTCAG |
| ATGCTCAC | CAAGCAGAAGACGGCATACGAGATgtgagcatGTGACTGGAGTTCAG |
| TCGATTGA | CAAGCAGAAGACGGCATACGAGATtcaatcgaGTGACTGGAGTTCAG |
| CAGCAGTA | CAAGCAGAAGACGGCATACGAGATtactgctgGTGACTGGAGTTCAG |
| TTTCCCTA | CAAGCAGAAGACGGCATACGAGATtagggaaaGTGACTGGAGTTCA |
| CCTCCATC | CAAGCAGAAGACGGCATACGAGATgatggaggGTGACTGGAGTTCA |
| GATCCACG | CAAGCAGAAGACGGCATACGAGATcgtggatcGTGACTGGAGTTCAG |
| TTCTTCCC | CAAGCAGAAGACGGCATACGAGATgggaagaaGTGACTGGAGTTCA |
| AAGTGCTA | CAAGCAGAAGACGGCATACGAGATtagcacttGTGACTGGAGTTCAG |
| CGAGAAAA | CAAGCAGAAGACGGCATACGAGATttttctcgGTGACTGGAGTTCAGA |
| ATATTCCG | CAAGCAGAAGACGGCATACGAGATcggaatatGTGACTGGAGTTCAG |
| CGTCAGCA | CAAGCAGAAGACGGCATACGAGATtgctgacgGTGACTGGAGTTCAG |
| TCGCAACT | CAAGCAGAAGACGGCATACGAGATagttgcgaGTGACTGGAGTTCAG |
| AGATCATC | CAAGCAGAAGACGGCATACGAGATgatgatctGTGACTGGAGTTCAG |
| GGCATCCA | CAAGCAGAAGACGGCATACGAGATtggatgccGTGACTGGAGTTCAG |
| GTATTTAT | CAAGCAGAAGACGGCATACGAGATataaatacGTGACTGGAGTTCAG |
| GTCCATGC | CAAGCAGAAGACGGCATACGAGATgcatggacGTGACTGGAGTTCA |
| GCATCTTC | CAAGCAGAAGACGGCATACGAGATgaagatgcGTGACTGGAGTTCA |
| GACGAAAA | CAAGCAGAAGACGGCATACGAGATttttcgtcGTGACTGGAGTTCAGA |
| TCCCTAGT | CAAGCAGAAGACGGCATACGAGATactagggaGTGACTGGAGTTCA |
| CCCACAGC | CAAGCAGAAGACGGCATACGAGATgctgtgggGTGACTGGAGTTCAG |
| TATATCTG | CAAGCAGAAGACGGCATACGAGATcagatataGTGACTGGAGTTCAG |
| CAAATATC | CAAGCAGAAGACGGCATACGAGATgatatttgGTGACTGGAGTTCAG |
| TAGGCTGT | CAAGCAGAAGACGGCATACGAGATacagcctaGTGACTGGAGTTCAG |
| AGAGATCG | CAAGCAGAAGACGGCATACGAGATcgatctctGTGACTGGAGTTCAG |
| TACGCGGG | CAAGCAGAAGACGGCATACGAGATcccgcgtaGTGACTGGAGTTCAG |
| CTGTTCCA | CAAGCAGAAGACGGCATACGAGATtggaacagGTGACTGGAGTTCA |
| CTTAGATC | CAAGCAGAAGACGGCATACGAGATgatctaagGTGACTGGAGTTCAG |
| ATGGGTAC | CAAGCAGAAGACGGCATACGAGATgtacccatGTGACTGGAGTTCAG |
| TTAGAGCT | CAAGCAGAAGACGGCATACGAGATagctctaaGTGACTGGAGTTCAG |
| GTTGCATT | CAAGCAGAAGACGGCATACGAGATaatgcaacGTGACTGGAGTTCA |
| ATCACTGA | CAAGCAGAAGACGGCATACGAGATtcagtgatGTGACTGGAGTTCAG |
| GCATGCCA | CAAGCAGAAGACGGCATACGAGATtggcatgcGTGACTGGAGTTCAG |
| AAACTAGT | CAAGCAGAAGACGGCATACGAGATactagtttGTGACTGGAGTTCAG |
| CATGGCCA | CAAGCAGAAGACGGCATACGAGATtggccatgGTGACTGGAGTTCAG |
| AGGTGTAA | CAAGCAGAAGACGGCATACGAGATttacacctGTGACTGGAGTTCAG |
| GACTTGAC | CAAGCAGAAGACGGCATACGAGATgtcaagtcGTGACTGGAGTTCAG |
| TCACTATG | CAAGCAGAAGACGGCATACGAGATcatagtgaGTGACTGGAGTTCAG |
| TTCACTCT | CAAGCAGAAGACGGCATACGAGATagagtgaaGTGACTGGAGTTCA |
| TAGAAGTC | CAAGCAGAAGACGGCATACGAGATgacttctaGTGACTGGAGTTCAG |
| GAGGCAGC | CAAGCAGAAGACGGCATACGAGATgctgcctcGTGACTGGAGTTCAG |
| CTAAAAAT | CAAGCAGAAGACGGCATACGAGATatttttagGTGACTGGAGTTCAGA |
| ACATCACA | CAAGCAGAAGACGGCATACGAGATtgtgatgtGTGACTGGAGTTCAG |
| GGAAGTCC | CAAGCAGAAGACGGCATACGAGATggacttccGTGACTGGAGTTCAG |
| TGGGGACG | CAAGCAGAAGACGGCATACGAGATcgtccccaGTGACTGGAGTTCAG |
| ACGTACGC | CAAGCAGAAGACGGCATACGAGATgcgtacgtGTGACTGGAGTTCAG |
| ACAATAAA | CAAGCAGAAGACGGCATACGAGATtttattgtGTGACTGGAGTTCAGA |
| GAGTACTT | CAAGCAGAAGACGGCATACGAGATaagtactcGTGACTGGAGTTCAG |

| | |
|---|---|
| GTGCCGTA | CAAGCAGAAGACGGCATACGAGATtacggcacGTGACTGGAGTTCAG |
| CGAGGATG | CAAGCAGAAGACGGCATACGAGATcatcctcgGTGACTGGAGTTCAG |
| CGGAGCCG | CAAGCAGAAGACGGCATACGAGATcggctccgGTGACTGGAGTTCAG |
| GTGGTTGA | CAAGCAGAAGACGGCATACGAGATtcaaccacGTGACTGGAGTTCAG |
| CTATCGAT | CAAGCAGAAGACGGCATACGAGATatcgatagGTGACTGGAGTTCAG |
| GTCCCTTT | CAAGCAGAAGACGGCATACGAGATaaagggacGTGACTGGAGTTCA |
| TAAGGTCG | CAAGCAGAAGACGGCATACGAGATcgaccttaGTGACTGGAGTTCAG |
| CCGGCTTT | CAAGCAGAAGACGGCATACGAGATaaagccggGTGACTGGAGTTCA |
| ACACGTGC | CAAGCAGAAGACGGCATACGAGATgcacgtgtGTGACTGGAGTTCAG |
| CAGGCGAG | CAAGCAGAAGACGGCATACGAGATctcgcctgGTGACTGGAGTTCAG |
| TGAATTCG | CAAGCAGAAGACGGCATACGAGATcgaattcaGTGACTGGAGTTCAG |
| AACGTCCC | CAAGCAGAAGACGGCATACGAGATgggacgttGTGACTGGAGTTCAG |
| CTGCCATG | CAAGCAGAAGACGGCATACGAGATcatggcagGTGACTGGAGTTCA |
| TGTTATTA | CAAGCAGAAGACGGCATACGAGATtaataacaGTGACTGGAGTTCAG |
| GTTTGGAT | CAAGCAGAAGACGGCATACGAGATatccaaacGTGACTGGAGTTCA |
| CTACCGGG | CAAGCAGAAGACGGCATACGAGATcccggtagGTGACTGGAGTTCAG |
| TCAGACTT | CAAGCAGAAGACGGCATACGAGATaagtctgaGTGACTGGAGTTCAG |
| TATGCTTG | CAAGCAGAAGACGGCATACGAGATcaagcataGTGACTGGAGTTCA |
| GGCATTTC | CAAGCAGAAGACGGCATACGAGATgaaatgccGTGACTGGAGTTCA |
| AATGGAGA | CAAGCAGAAGACGGCATACGAGATtctccattGTGACTGGAGTTCAGA |
| AACACCGC | CAAGCAGAAGACGGCATACGAGATgcggtgttGTGACTGGAGTTCAG |
| ATCCGCTA | CAAGCAGAAGACGGCATACGAGATtagcggatGTGACTGGAGTTCAG |
| CCTCGGGT | CAAGCAGAAGACGGCATACGAGATacccgaggGTGACTGGAGTTCA |
| TGGACAGT | CAAGCAGAAGACGGCATACGAGATactgtccaGTGACTGGAGTTCAG |
| TGGGTCCT | CAAGCAGAAGACGGCATACGAGATaggacccaGTGACTGGAGTTCA |
| GCTACATG | CAAGCAGAAGACGGCATACGAGATcatgtagcGTGACTGGAGTTCAG |
| TTAGCCAG | CAAGCAGAAGACGGCATACGAGATctggctaaGTGACTGGAGTTCAG |
| GTTGTAAA | CAAGCAGAAGACGGCATACGAGATtttacaacGTGACTGGAGTTCAG |
| CGGGCAAC | CAAGCAGAAGACGGCATACGAGATgttgcccgGTGACTGGAGTTCAG |
| AAGCAAAC | CAAGCAGAAGACGGCATACGAGATgtttgcttGTGACTGGAGTTCAGA |
| TTAAGGGA | CAAGCAGAAGACGGCATACGAGATtcccttaaGTGACTGGAGTTCAG |
| CTCCTTCG | CAAGCAGAAGACGGCATACGAGATcgaaggagGTGACTGGAGTTCA |
| ATACCGCC | CAAGCAGAAGACGGCATACGAGATggcggtatGTGACTGGAGTTCAG |
| GTAAGAGC | CAAGCAGAAGACGGCATACGAGATgctcttacGTGACTGGAGTTCAG |
| ACCTGGTT | CAAGCAGAAGACGGCATACGAGATaaccaggtGTGACTGGAGTTCA |
| TACTATGG | CAAGCAGAAGACGGCATACGAGATccatagtaGTGACTGGAGTTCAG |
| GGCTACCC | CAAGCAGAAGACGGCATACGAGATgggtagccGTGACTGGAGTTCA |
| TAGATACT | CAAGCAGAAGACGGCATACGAGATagtatctaGTGACTGGAGTTCAG |
| GGACCCGG | CAAGCAGAAGACGGCATACGAGATccgggtccGTGACTGGAGTTCAG |
| CTCTTGTC | CAAGCAGAAGACGGCATACGAGATgacaagagGTGACTGGAGTTCA |
| CCATGCTC | CAAGCAGAAGACGGCATACGAGATgagcatggGTGACTGGAGTTCA |
| TAGCGTCC | CAAGCAGAAGACGGCATACGAGATggacgctaGTGACTGGAGTTCA |
| GCGAACTC | CAAGCAGAAGACGGCATACGAGATgagttcgcGTGACTGGAGTTCAG |
| TGAAGACA | CAAGCAGAAGACGGCATACGAGATtgtcttcaGTGACTGGAGTTCAGA |
| TACTGACA | CAAGCAGAAGACGGCATACGAGATtgtcagtaGTGACTGGAGTTCAG |
| CGACTCCA | CAAGCAGAAGACGGCATACGAGATtggagtcgGTGACTGGAGTTCAG |
| CAACAGGT | CAAGCAGAAGACGGCATACGAGATacctgttgGTGACTGGAGTTCAG |
| GCTCGATA | CAAGCAGAAGACGGCATACGAGATtatcgagcGTGACTGGAGTTCAG |
| TACCTTTA | CAAGCAGAAGACGGCATACGAGATtaaaggtaGTGACTGGAGTTCA |
| CGGATATT | CAAGCAGAAGACGGCATACGAGATaatatccgGTGACTGGAGTTCAG |
| CCGCAAAG | CAAGCAGAAGACGGCATACGAGATctttgcggGTGACTGGAGTTCAG |
| AAGTGAGT | CAAGCAGAAGACGGCATACGAGATactcacttGTGACTGGAGTTCAG |
| CCATGGCT | CAAGCAGAAGACGGCATACGAGATagccatggGTGACTGGAGTTCA |

| | |
|---|---|
| CAGGTCTG | CAAGCAGAAGACGGCATACGAGATcagacctgGTGACTGGAGTTCAG |
| GCCTTAAT | CAAGCAGAAGACGGCATACGAGATattaaggcGTGACTGGAGTTCAG |
| GCGGTGAA | CAAGCAGAAGACGGCATACGAGATttcaccgcGTGACTGGAGTTCAG |
| TAGCCGCG | CAAGCAGAAGACGGCATACGAGATcgcggctaGTGACTGGAGTTCAG |
| CTCACAAG | CAAGCAGAAGACGGCATACGAGATcttgtgagGTGACTGGAGTTCAG |
| GCCTCGTA | CAAGCAGAAGACGGCATACGAGATtacgaggcGTGACTGGAGTTCAG |
| AATACATT | CAAGCAGAAGACGGCATACGAGATaatgtattGTGACTGGAGTTCAGA |
| ATTCGGCT | CAAGCAGAAGACGGCATACGAGATagccgaatGTGACTGGAGTTCAG |
| CAGACTAC | CAAGCAGAAGACGGCATACGAGATgtagtctgGTGACTGGAGTTCAG |
| CCAACTGT | CAAGCAGAAGACGGCATACGAGATacagttggGTGACTGGAGTTCAG |
| TTGATTTG | CAAGCAGAAGACGGCATACGAGATcaaatcaaGTGACTGGAGTTCAG |
| GTTCAAAC | CAAGCAGAAGACGGCATACGAGATgtttgaacGTGACTGGAGTTCAG |
| CTCCTGGT | CAAGCAGAAGACGGCATACGAGATaccaggagGTGACTGGAGTTCA |
| GGATTCTG | CAAGCAGAAGACGGCATACGAGATcagaatccGTGACTGGAGTTCAG |
| GCGACTGC | CAAGCAGAAGACGGCATACGAGATgcagtcgcGTGACTGGAGTTCAG |
| GGAGATGA | CAAGCAGAAGACGGCATACGAGATtcatctccGTGACTGGAGTTCAGA |
| CCGGAGCC | CAAGCAGAAGACGGCATACGAGATggctccggGTGACTGGAGTTCAG |
| TGGTCTCT | CAAGCAGAAGACGGCATACGAGATagagaccaGTGACTGGAGTTCA |
| CCTTGAGA | CAAGCAGAAGACGGCATACGAGATtctcaaggGTGACTGGAGTTCAG |
| AAAGCGAC | CAAGCAGAAGACGGCATACGAGATgtcgctttGTGACTGGAGTTCAGA |
| CCGCTCTA | CAAGCAGAAGACGGCATACGAGATtagagcggGTGACTGGAGTTCAG |
| CGCTTACA | CAAGCAGAAGACGGCATACGAGATtgtaagcgGTGACTGGAGTTCAG |
| GTACTCGT | CAAGCAGAAGACGGCATACGAGATacgagtacGTGACTGGAGTTCAG |
| CACACGAA | CAAGCAGAAGACGGCATACGAGATttcgtgtgGTGACTGGAGTTCAGA |
| TTAATTGT | CAAGCAGAAGACGGCATACGAGATacaattaaGTGACTGGAGTTCAG |
| GTGGCACA | CAAGCAGAAGACGGCATACGAGATtgtgccacGTGACTGGAGTTCAG |
| ACGGTGTC | CAAGCAGAAGACGGCATACGAGATgacaccgtGTGACTGGAGTTCAG |
| AGCCGATG | CAAGCAGAAGACGGCATACGAGATcatcggctGTGACTGGAGTTCAG |
| ACTGTCAC | CAAGCAGAAGACGGCATACGAGATgtgacagtGTGACTGGAGTTCAG |
| AATTCCAG | CAAGCAGAAGACGGCATACGAGATctggaattGTGACTGGAGTTCAG |
| CGCAAGCG | CAAGCAGAAGACGGCATACGAGATcgcttgcgGTGACTGGAGTTCAG |
| GCAATGAG | CAAGCAGAAGACGGCATACGAGATctcattgcGTGACTGGAGTTCAGA |
| TTTCTTGC | CAAGCAGAAGACGGCATACGAGATgcaagaaaGTGACTGGAGTTCA |
| GCTTTGGA | CAAGCAGAAGACGGCATACGAGATtccaaagcGTGACTGGAGTTCAG |
| CGGCCCGT | CAAGCAGAAGACGGCATACGAGATacgggccgGTGACTGGAGTTCAG |
| GTGACCCG | CAAGCAGAAGACGGCATACGAGATcgggtcacGTGACTGGAGTTCAG |
| TGGGAGAG | CAAGCAGAAGACGGCATACGAGATctctcccaGTGACTGGAGTTCAGA |
| ATAAGACT | CAAGCAGAAGACGGCATACGAGATagtcttatGTGACTGGAGTTCAGA |
| CAATTACG | CAAGCAGAAGACGGCATACGAGATcgtaattgGTGACTGGAGTTCAG |
| TGAGTGTG | CAAGCAGAAGACGGCATACGAGATcacactcaGTGACTGGAGTTCAG |
| ACGCAGGA | CAAGCAGAAGACGGCATACGAGATtcctgcgtGTGACTGGAGTTCAGA |
| TAGCTCTT | CAAGCAGAAGACGGCATACGAGATaagagctaGTGACTGGAGTTCAG |
| TGGAATGC | CAAGCAGAAGACGGCATACGAGATgcattccaGTGACTGGAGTTCAG |
| AGCAGGCT | CAAGCAGAAGACGGCATACGAGATagcctgctGTGACTGGAGTTCAG |
| CACCGCTG | CAAGCAGAAGACGGCATACGAGATcagcggtgGTGACTGGAGTTCAG |
| AACCGGGA | CAAGCAGAAGACGGCATACGAGATtcccggttGTGACTGGAGTTCAGA |
| GAAGAATC | CAAGCAGAAGACGGCATACGAGATgattcttcGTGACTGGAGTTCAGA |
| GCTTGCAC | CAAGCAGAAGACGGCATACGAGATgtgcaagcGTGACTGGAGTTCAG |
| AGAAGTGT | CAAGCAGAAGACGGCATACGAGATacacttctGTGACTGGAGTTCAGA |
| TTCAGCCA | CAAGCAGAAGACGGCATACGAGATtggctgaaGTGACTGGAGTTCAG |
| ACAAACCA | CAAGCAGAAGACGGCATACGAGATtggtttgtGTGACTGGAGTTCAGA |
| CATTTTAA | CAAGCAGAAGACGGCATACGAGATttaaaatgGTGACTGGAGTTCAG |
| CCTGGTCT | CAAGCAGAAGACGGCATACGAGATagaccaggGTGACTGGAGTTCA |

| | |
|---|---|
| CACAGGGT | CAAGCAGAAGACGGCATACGAGATaccctgtgGTGACTGGAGTTCAG |
| TAACTGAG | CAAGCAGAAGACGGCATACGAGATctcagttaGTGACTGGAGTTCAG |
| ACGGGCCA | CAAGCAGAAGACGGCATACGAGATtggcccgtGTGACTGGAGTTCAG |
| CTAGGGAG | CAAGCAGAAGACGGCATACGAGATctccctagGTGACTGGAGTTCAG |
| TCCCCTCC | CAAGCAGAAGACGGCATACGAGATggaggggaGTGACTGGAGTTCA |
| CGACCATT | CAAGCAGAAGACGGCATACGAGATaatggtcgGTGACTGGAGTTCAG |
| CATATACA | CAAGCAGAAGACGGCATACGAGATtgtatatgGTGACTGGAGTTCAG |
| TAGTCGGA | CAAGCAGAAGACGGCATACGAGATtccgactaGTGACTGGAGTTCAG |
| GTAGAAGT | CAAGCAGAAGACGGCATACGAGATacttctacGTGACTGGAGTTCAG |
| CGCCACTA | CAAGCAGAAGACGGCATACGAGATtagtggcgGTGACTGGAGTTCAG |
| TTGAACAG | CAAGCAGAAGACGGCATACGAGATctgttcaaGTGACTGGAGTTCAG |
| AAAGTGGA | CAAGCAGAAGACGGCATACGAGATtccactttGTGACTGGAGTTCAGA |
| TGTCGTAG | CAAGCAGAAGACGGCATACGAGATctacgacaGTGACTGGAGTTCAG |
| CATCTTTT | CAAGCAGAAGACGGCATACGAGATaaaagatgGTGACTGGAGTTCA |
| GGCGGCTT | CAAGCAGAAGACGGCATACGAGATaagccgccGTGACTGGAGTTCA |
| TCCTAACG | CAAGCAGAAGACGGCATACGAGATcgttaggaGTGACTGGAGTTCAG |
| ACGCCCAT | CAAGCAGAAGACGGCATACGAGATatgggcgtGTGACTGGAGTTCAG |
| ATAAACTT | CAAGCAGAAGACGGCATACGAGATaagtttatGTGACTGGAGTTCAG |
| TCCGGGTA | CAAGCAGAAGACGGCATACGAGATtacccggaGTGACTGGAGTTCAG |
| TGACGGCG | CAAGCAGAAGACGGCATACGAGATcgccgtcaGTGACTGGAGTTCAG |
| ACACGGCA | CAAGCAGAAGACGGCATACGAGATtgccgtgtGTGACTGGAGTTCAG |
| GCATGATT | CAAGCAGAAGACGGCATACGAGATaatcatgcGTGACTGGAGTTCAG |
| ATGTACTG | CAAGCAGAAGACGGCATACGAGATcagtacatGTGACTGGAGTTCAG |
| ATACATAT | CAAGCAGAAGACGGCATACGAGATatatgtatGTGACTGGAGTTCAG |
| AGTGTTGG | CAAGCAGAAGACGGCATACGAGATccaacactGTGACTGGAGTTCAG |
| AGTCACTT | CAAGCAGAAGACGGCATACGAGATaagtgactGTGACTGGAGTTCAG |
| TCTTCATA | CAAGCAGAAGACGGCATACGAGATtatgaagaGTGACTGGAGTTCAG |
| ATGGTTTT | CAAGCAGAAGACGGCATACGAGATaaaaccatGTGACTGGAGTTCA |
| GGTCTTGT | CAAGCAGAAGACGGCATACGAGATacaagaccGTGACTGGAGTTCA |
| TAAGCCCT | CAAGCAGAAGACGGCATACGAGATagggcttaGTGACTGGAGTTCAG |
| ACCTGCGG | CAAGCAGAAGACGGCATACGAGATccgcaggtGTGACTGGAGTTCAG |
| AGGAGAGC | CAAGCAGAAGACGGCATACGAGATgctctcctGTGACTGGAGTTCAG |
| GATTAAGC | CAAGCAGAAGACGGCATACGAGATgcttaatcGTGACTGGAGTTCAG |
| CTGCTCGG | CAAGCAGAAGACGGCATACGAGATccgagcagGTGACTGGAGTTCA |
| CGGCACAC | CAAGCAGAAGACGGCATACGAGATgtgtgccgGTGACTGGAGTTCAG |
| CGTACCCT | CAAGCAGAAGACGGCATACGAGATagggtacgGTGACTGGAGTTCA |
| GAAGGCAT | CAAGCAGAAGACGGCATACGAGATatgccttcGTGACTGGAGTTCAG |
| CCGTGCAG | CAAGCAGAAGACGGCATACGAGATctgcacggGTGACTGGAGTTCAG |
| AGAATGAC | CAAGCAGAAGACGGCATACGAGATgtcattctGTGACTGGAGTTCAGA |
| CAATCGCA | CAAGCAGAAGACGGCATACGAGATtgcgattgGTGACTGGAGTTCAG |
| ATGAAATA | CAAGCAGAAGACGGCATACGAGATtatttcatGTGACTGGAGTTCAGA |
| CCCAGCGA | CAAGCAGAAGACGGCATACGAGATtcgctgggGTGACTGGAGTTCAG |
| AACCCCCA | CAAGCAGAAGACGGCATACGAGATtgggggttGTGACTGGAGTTCAG |
| TACAGCGG | CAAGCAGAAGACGGCATACGAGATccgctgtaGTGACTGGAGTTCAG |
| CTGCATTT | CAAGCAGAAGACGGCATACGAGATaaatgcagGTGACTGGAGTTCA |
| ATCCCAGG | CAAGCAGAAGACGGCATACGAGATcctgggatGTGACTGGAGTTCAG |
| GTCTCGCG | CAAGCAGAAGACGGCATACGAGATcgcgagacGTGACTGGAGTTCA |
| GGCAGAGG | CAAGCAGAAGACGGCATACGAGATcctctgccGTGACTGGAGTTCAG |
| TTTATATT | CAAGCAGAAGACGGCATACGAGATaatataaaGTGACTGGAGTTCAG |
| AGATGGTG | CAAGCAGAAGACGGCATACGAGATcaccatctGTGACTGGAGTTCAG |
| TTGCCTGG | CAAGCAGAAGACGGCATACGAGATccaggcaaGTGACTGGAGTTCA |
| CAGGCCCC | CAAGCAGAAGACGGCATACGAGATggggcctgGTGACTGGAGTTCA |
| TTGCAGCC | CAAGCAGAAGACGGCATACGAGATggctgcaaGTGACTGGAGTTCA |

| | |
|---|---|
| TGCTTGAA | CAAGCAGAAGACGGCATACGAGATttcaagcaGTGACTGGAGTTCAG |
| AATACGGG | CAAGCAGAAGACGGCATACGAGATcccgtattGTGACTGGAGTTCAG |
| GGTTACAT | CAAGCAGAAGACGGCATACGAGATatgtaaccGTGACTGGAGTTCAG |
| TTTCACGT | CAAGCAGAAGACGGCATACGAGATacgtgaaaGTGACTGGAGTTCA |
| GAGTTTCG | CAAGCAGAAGACGGCATACGAGATcgaaactcGTGACTGGAGTTCAG |
| CTGGTGCG | CAAGCAGAAGACGGCATACGAGATcgcaccagGTGACTGGAGTTCA |
| CGTAATGT | CAAGCAGAAGACGGCATACGAGATacattacgGTGACTGGAGTTCAG |
| TATTGCCT | CAAGCAGAAGACGGCATACGAGATaggcaataGTGACTGGAGTTCA |
| ACTTGTAG | CAAGCAGAAGACGGCATACGAGATctacaagtGTGACTGGAGTTCAG |
| TCCTGTAC | CAAGCAGAAGACGGCATACGAGATgtacaggaGTGACTGGAGTTCA |
| GACCAGTC | CAAGCAGAAGACGGCATACGAGATgactggtcGTGACTGGAGTTCAG |
| CCGTAAGT | CAAGCAGAAGACGGCATACGAGATacttacggGTGACTGGAGTTCAG |
| GTGATGCC | CAAGCAGAAGACGGCATACGAGATggcatcacGTGACTGGAGTTCAG |
| CAACCCCG | CAAGCAGAAGACGGCATACGAGATcggggttgGTGACTGGAGTTCAG |
| ATCGGGCG | CAAGCAGAAGACGGCATACGAGATcgcccgatGTGACTGGAGTTCAG |
| ATTCTGGG | CAAGCAGAAGACGGCATACGAGATcccagaatGTGACTGGAGTTCAG |
| AAGGTCGT | CAAGCAGAAGACGGCATACGAGATacgaccttGTGACTGGAGTTCAG |
| CCATTTAC | CAAGCAGAAGACGGCATACGAGATgtaaatggGTGACTGGAGTTCAG |
| TCCACCTG | CAAGCAGAAGACGGCATACGAGATcaggtggaGTGACTGGAGTTCA |
| ACGTGACG | CAAGCAGAAGACGGCATACGAGATcgtcacgtGTGACTGGAGTTCAG |
| TTATGAAG | CAAGCAGAAGACGGCATACGAGATcttcataaGTGACTGGAGTTCAG |
| GAATAGAG | CAAGCAGAAGACGGCATACGAGATctctattcGTGACTGGAGTTCAGA |
| AGGATCTC | CAAGCAGAAGACGGCATACGAGATgagatcctGTGACTGGAGTTCAG |
| AAATACTC | CAAGCAGAAGACGGCATACGAGATgagtatttGTGACTGGAGTTCAG |
| GAAGTGCC | CAAGCAGAAGACGGCATACGAGATggcacttcGTGACTGGAGTTCAG |
| GGCCAGCT | CAAGCAGAAGACGGCATACGAGATagctggccGTGACTGGAGTTCAG |
| CAACGATA | CAAGCAGAAGACGGCATACGAGATtatcgttgGTGACTGGAGTTCAG |
| TGTAGCTC | CAAGCAGAAGACGGCATACGAGATgagctacaGTGACTGGAGTTCA |
| CACGCAGA | CAAGCAGAAGACGGCATACGAGATtctgcgtgGTGACTGGAGTTCAG |
| ATCTTTGT | CAAGCAGAAGACGGCATACGAGATacaaagatGTGACTGGAGTTCA |
| GTTAAGTT | CAAGCAGAAGACGGCATACGAGATaacttaacGTGACTGGAGTTCAG |
| TATGATCC | CAAGCAGAAGACGGCATACGAGATggatcataGTGACTGGAGTTCAG |
| CCGCTAGC | CAAGCAGAAGACGGCATACGAGATgctagcggGTGACTGGAGTTCA |
| TGCCCGTT | CAAGCAGAAGACGGCATACGAGATaacgggcaGTGACTGGAGTTCA |
| GCCGTCGT | CAAGCAGAAGACGGCATACGAGATacgacggcGTGACTGGAGTTCA |
| ACCAATAC | CAAGCAGAAGACGGCATACGAGATgtattggtGTGACTGGAGTTCAG |
| ACTTAGTG | CAAGCAGAAGACGGCATACGAGATcactaagtGTGACTGGAGTTCAG |
| GTTAGCTG | CAAGCAGAAGACGGCATACGAGATcagctaacGTGACTGGAGTTCA |
| TCGTTTCC | CAAGCAGAAGACGGCATACGAGATggaaacgaGTGACTGGAGTTCA |
| GGAACACT | CAAGCAGAAGACGGCATACGAGATagtgttccGTGACTGGAGTTCAG |
| ATTTGCCC | CAAGCAGAAGACGGCATACGAGATgggcaaatGTGACTGGAGTTCA |
| GTACACTA | CAAGCAGAAGACGGCATACGAGATtagtgtacGTGACTGGAGTTCAG |
| ACCCCGGC | CAAGCAGAAGACGGCATACGAGATgccggggtGTGACTGGAGTTCA |
| CTTCTCAA | CAAGCAGAAGACGGCATACGAGATttgagaagGTGACTGGAGTTCAG |
| AAAAGGTC | CAAGCAGAAGACGGCATACGAGATgaccttttGTGACTGGAGTTCAGA |
| CTTAGTAA | CAAGCAGAAGACGGCATACGAGATttactaagGTGACTGGAGTTCAG |
| ATTATACC | CAAGCAGAAGACGGCATACGAGATggtataatGTGACTGGAGTTCAG |
| TCTGAGTC | CAAGCAGAAGACGGCATACGAGATgactcagaGTGACTGGAGTTCA |
| AGGTAGAT | CAAGCAGAAGACGGCATACGAGATatctacctGTGACTGGAGTTCAG |
| CTCACTTC | CAAGCAGAAGACGGCATACGAGATgaagtgagGTGACTGGAGTTCA |
| AGTACGCA | CAAGCAGAAGACGGCATACGAGATtgcgtactGTGACTGGAGTTCAG |
| CATTGTGT | CAAGCAGAAGACGGCATACGAGATacacaatgGTGACTGGAGTTCA |
| GAGCGCTC | CAAGCAGAAGACGGCATACGAGATgagcgctcGTGACTGGAGTTCAG |

| | |
|---|---|
| TCTACGAG | CAAGCAGAAGACGGCATACGAGATctcgtagaGTGACTGGAGTTCAG |
| ACCATCAT | CAAGCAGAAGACGGCATACGAGATatgatggtGTGACTGGAGTTCAG |
| TGTCAATC | CAAGCAGAAGACGGCATACGAGATgattgacaGTGACTGGAGTTCAG |
| AGACTTTC | CAAGCAGAAGACGGCATACGAGATgaaagtctGTGACTGGAGTTCAG |
| TATGTAAC | CAAGCAGAAGACGGCATACGAGATgttacataGTGACTGGAGTTCAG |
| GAAGCTTT | CAAGCAGAAGACGGCATACGAGATaaagcttcGTGACTGGAGTTCAG |
| CGACAACC | CAAGCAGAAGACGGCATACGAGATggttgtcgGTGACTGGAGTTCAG |
| AGCGCCTC | CAAGCAGAAGACGGCATACGAGATgaggcgctGTGACTGGAGTTCAG |
| CTTATGAC | CAAGCAGAAGACGGCATACGAGATgtcataagGTGACTGGAGTTCAG |
| CCAGATGC | CAAGCAGAAGACGGCATACGAGATgcatctggGTGACTGGAGTTCAG |
| CATCCTAG | CAAGCAGAAGACGGCATACGAGATctaggatgGTGACTGGAGTTCAG |
| AAATTTTG | CAAGCAGAAGACGGCATACGAGATcaaaatttGTGACTGGAGTTCAG |
| GCTGTACT | CAAGCAGAAGACGGCATACGAGATagtacagcGTGACTGGAGTTCAG |
| ATCCTATT | CAAGCAGAAGACGGCATACGAGATaataggatGTGACTGGAGTTCAG |
| ACCGTTAG | CAAGCAGAAGACGGCATACGAGATctaacggtGTGACTGGAGTTCAG |
| TTGTAGTT | CAAGCAGAAGACGGCATACGAGATaactacaaGTGACTGGAGTTCAG |
| TCGGATCG | CAAGCAGAAGACGGCATACGAGATcgatccgaGTGACTGGAGTTCAG |
| AGGAAGCC | CAAGCAGAAGACGGCATACGAGATggcttcctGTGACTGGAGTTCAG |
| CGGGGCGA | CAAGCAGAAGACGGCATACGAGATtcgccccgGTGACTGGAGTTCAG |
| GGGCCACC | CAAGCAGAAGACGGCATACGAGATggtggcccGTGACTGGAGTTCAG |
| GAGAGGGC | CAAGCAGAAGACGGCATACGAGATgccctctcGTGACTGGAGTTCAG |
| TCGCGTTA | CAAGCAGAAGACGGCATACGAGATtaacgcgaGTGACTGGAGTTCAG |
| TATGGGTT | CAAGCAGAAGACGGCATACGAGATaacccataGTGACTGGAGTTCAG |
| CTTGTTCC | CAAGCAGAAGACGGCATACGAGATggaacaagGTGACTGGAGTTCA |
| GTAAGGCG | CAAGCAGAAGACGGCATACGAGATcgccttacGTGACTGGAGTTCAG |
| TGTCAGAT | CAAGCAGAAGACGGCATACGAGATatctgacaGTGACTGGAGTTCAG |
| GTCCGACG | CAAGCAGAAGACGGCATACGAGATcgtcggacGTGACTGGAGTTCAG |
| GTCAGCGT | CAAGCAGAAGACGGCATACGAGATacgctgacGTGACTGGAGTTCAG |
| CGATGTGC | CAAGCAGAAGACGGCATACGAGATgcacatcgGTGACTGGAGTTCAG |
| TTTGTGTA | CAAGCAGAAGACGGCATACGAGATtacacaaaGTGACTGGAGTTCAG |
| TAGACCAT | CAAGCAGAAGACGGCATACGAGATatggtctaGTGACTGGAGTTCAG |
| GACGTATG | CAAGCAGAAGACGGCATACGAGATcatacgtcGTGACTGGAGTTCAG |
| CGCGGTCG | CAAGCAGAAGACGGCATACGAGATcgaccgcgGTGACTGGAGTTCAG |
| ATACAAGA | CAAGCAGAAGACGGCATACGAGATtcttgtatGTGACTGGAGTTCAGA |
| TTTTCACG | CAAGCAGAAGACGGCATACGAGATcgtgaaaaGTGACTGGAGTTCAG |
| AATAACAA | CAAGCAGAAGACGGCATACGAGATttgttattGTGACTGGAGTTCAGA |
| TGTATGCC | CAAGCAGAAGACGGCATACGAGATggcatacaGTGACTGGAGTTCAG |
| TTTAGTCG | CAAGCAGAAGACGGCATACGAGATcgactaaaGTGACTGGAGTTCAG |
| TGATATAT | CAAGCAGAAGACGGCATACGAGATatatatcaGTGACTGGAGTTCAG |
| AGCACTTG | CAAGCAGAAGACGGCATACGAGATcaagtgctGTGACTGGAGTTCAG |
| TGACTGGA | CAAGCAGAAGACGGCATACGAGATtccagtcaGTGACTGGAGTTCAG |
| TTGTGCAC | CAAGCAGAAGACGGCATACGAGATgtgcacaaGTGACTGGAGTTCAG |
| GAATATGT | CAAGCAGAAGACGGCATACGAGATacatattcGTGACTGGAGTTCAG |
| TCGTCTTG | CAAGCAGAAGACGGCATACGAGATcaagacgaGTGACTGGAGTTCA |
| AACCAACT | CAAGCAGAAGACGGCATACGAGATagttggttGTGACTGGAGTTCAG |
| GGGTTAGA | CAAGCAGAAGACGGCATACGAGATtctaacccGTGACTGGAGTTCAG |
| ACTCTTCG | CAAGCAGAAGACGGCATACGAGATcgaagagtGTGACTGGAGTTCAG |
| ACGGTAAT | CAAGCAGAAGACGGCATACGAGATattaccgtGTGACTGGAGTTCAG |
| TCTAATGG | CAAGCAGAAGACGGCATACGAGATccattagaGTGACTGGAGTTCAG |
| CGGTGGGT | CAAGCAGAAGACGGCATACGAGATacccaccgGTGACTGGAGTTCAG |
| GTGAGTAG | CAAGCAGAAGACGGCATACGAGATctactcacGTGACTGGAGTTCAG |
| AAGCATCG | CAAGCAGAAGACGGCATACGAGATcgatgcttGTGACTGGAGTTCAG |
| TTATCATT | CAAGCAGAAGACGGCATACGAGATaatgataaGTGACTGGAGTTCAG |

| | |
|---|---|
| TGAATCGC | CAAGCAGAAGACGGCATACGAGATgcgattcaGTGACTGGAGTTCAG |
| GCGTCCCC | CAAGCAGAAGACGGCATACGAGATggggacgcGTGACTGGAGTTCA |
| GCCATTCG | CAAGCAGAAGACGGCATACGAGATcgaatggcGTGACTGGAGTTCA |
| TCCCACGA | CAAGCAGAAGACGGCATACGAGATtcgtgggaGTGACTGGAGTTCAG |
| TAGTGATC | CAAGCAGAAGACGGCATACGAGATgatcactaGTGACTGGAGTTCAG |
| TACCCAAA | CAAGCAGAAGACGGCATACGAGATtttgggtaGTGACTGGAGTTCAG |
| TTCCTTAT | CAAGCAGAAGACGGCATACGAGATataaggaaGTGACTGGAGTTCA |
| AAAATTGC | CAAGCAGAAGACGGCATACGAGATgcaattttGTGACTGGAGTTCAG |
| TATTGGAA | CAAGCAGAAGACGGCATACGAGATttccaataGTGACTGGAGTTCAG |
| CGTTTCGC | CAAGCAGAAGACGGCATACGAGATgcgaaacgGTGACTGGAGTTCA |
| TCTTAGGT | CAAGCAGAAGACGGCATACGAGATacctaagaGTGACTGGAGTTCA |
| TAACGGGC | CAAGCAGAAGACGGCATACGAGATgcccgttaGTGACTGGAGTTCAG |
| CATCACGC | CAAGCAGAAGACGGCATACGAGATgcgtgatgGTGACTGGAGTTCAG |
| TAAGGCTC | CAAGCAGAAGACGGCATACGAGATgagccttaGTGACTGGAGTTCAG |
| GTAGCGTC | CAAGCAGAAGACGGCATACGAGATgacgctacGTGACTGGAGTTCA |
| CGGTAATA | CAAGCAGAAGACGGCATACGAGATtattaccgGTGACTGGAGTTCAG |
| GGGTCGTG | CAAGCAGAAGACGGCATACGAGATcacgacccGTGACTGGAGTTCA |
| CCCAAGTC | CAAGCAGAAGACGGCATACGAGATgacttgggGTGACTGGAGTTCAG |
| AACAAGCA | CAAGCAGAAGACGGCATACGAGATtgcttgttGTGACTGGAGTTCAGA |
| GGCGCCCG | CAAGCAGAAGACGGCATACGAGATcgggcgccGTGACTGGAGTTCA |
| AGTGGAAT | CAAGCAGAAGACGGCATACGAGATattccactGTGACTGGAGTTCAG |
| ACACGCTG | CAAGCAGAAGACGGCATACGAGATcagcgtgtGTGACTGGAGTTCAG |
| CCACGACG | CAAGCAGAAGACGGCATACGAGATcgtcgtggGTGACTGGAGTTCAG |
| TAGTAAAG | CAAGCAGAAGACGGCATACGAGATctttactaGTGACTGGAGTTCAG |
| TCGGTACA | CAAGCAGAAGACGGCATACGAGATtgtaccgaGTGACTGGAGTTCAG |
| AATTCTTC | CAAGCAGAAGACGGCATACGAGATgaagaattGTGACTGGAGTTCA |
| GGTATGAA | CAAGCAGAAGACGGCATACGAGATttcataccGTGACTGGAGTTCAG |
| CGCAACTG | CAAGCAGAAGACGGCATACGAGATcagttgcgGTGACTGGAGTTCAG |
| AGGTCAAT | CAAGCAGAAGACGGCATACGAGATattgacctGTGACTGGAGTTCAG |
| AGACCTGC | CAAGCAGAAGACGGCATACGAGATgcaggtctGTGACTGGAGTTCAG |
| GCCAGCCC | CAAGCAGAAGACGGCATACGAGATgggctggcGTGACTGGAGTTCA |
| GGTCCTTT | CAAGCAGAAGACGGCATACGAGATaaaggaccGTGACTGGAGTTCA |
| AATTTGGT | CAAGCAGAAGACGGCATACGAGATaccaaattGTGACTGGAGTTCAG |
| GCCGACTC | CAAGCAGAAGACGGCATACGAGATgagtcggcGTGACTGGAGTTCA |
| TCCAATCG | CAAGCAGAAGACGGCATACGAGATcgattggaGTGACTGGAGTTCAG |
| GGGAGTTC | CAAGCAGAAGACGGCATACGAGATgaactcccGTGACTGGAGTTCAG |
| AGGCTGAC | CAAGCAGAAGACGGCATACGAGATgtcagcctGTGACTGGAGTTCAG |
| GCGACGCG | CAAGCAGAAGACGGCATACGAGATcgcgtcgcGTGACTGGAGTTCAG |
| ACGCGCTT | CAAGCAGAAGACGGCATACGAGATaagcgcgtGTGACTGGAGTTCA |
| AAATTGGC | CAAGCAGAAGACGGCATACGAGATgccaatttGTGACTGGAGTTCAG |
| TCTGTTAG | CAAGCAGAAGACGGCATACGAGATctaacagaGTGACTGGAGTTCA |
| GCGCGCCA | CAAGCAGAAGACGGCATACGAGATtggcgcgcGTGACTGGAGTTCA |
| AGCTCCGG | CAAGCAGAAGACGGCATACGAGATccggagctGTGACTGGAGTTCA |
| GAACGCAG | CAAGCAGAAGACGGCATACGAGATctgcgttcGTGACTGGAGTTCAG |
| GGCTCGAC | CAAGCAGAAGACGGCATACGAGATgtcgagccGTGACTGGAGTTCA |
| CGCCTGAT | CAAGCAGAAGACGGCATACGAGATatcaggcgGTGACTGGAGTTCA |
| CTTGCCGT | CAAGCAGAAGACGGCATACGAGATacggcaagGTGACTGGAGTTCA |
| CCAGCCTT | CAAGCAGAAGACGGCATACGAGATaaggctggGTGACTGGAGTTCA |
| CGAATGGA | CAAGCAGAAGACGGCATACGAGATtccattcgGTGACTGGAGTTCAG |
| CAGCTGTG | CAAGCAGAAGACGGCATACGAGATcacagctgGTGACTGGAGTTCA |
| TTGCTGGA | CAAGCAGAAGACGGCATACGAGATtccagcaaGTGACTGGAGTTCA |
| CTGAGTGT | CAAGCAGAAGACGGCATACGAGATacactcagGTGACTGGAGTTCA |
| AAGTCACT | CAAGCAGAAGACGGCATACGAGATagtgacttGTGACTGGAGTTCAG |

| | |
|---|---|
| CATCAGTG | CAAGCAGAAGACGGCATACGAGATcactgatgGTGACTGGAGTTCAG |
| CGATAGTA | CAAGCAGAAGACGGCATACGAGATtactatcgGTGACTGGAGTTCAG |
| GTGCTTCT | CAAGCAGAAGACGGCATACGAGATagaagcacGTGACTGGAGTTCA |
| GGAAAAAA | CAAGCAGAAGACGGCATACGAGATttttttccGTGACTGGAGTTCAGA |
| TCATCACT | CAAGCAGAAGACGGCATACGAGATagtgatgaGTGACTGGAGTTCAG |
| GACCGCTT | CAAGCAGAAGACGGCATACGAGATaagcggtcGTGACTGGAGTTCA |
| AGCCGCGA | CAAGCAGAAGACGGCATACGAGATtcgcggctGTGACTGGAGTTCAG |
| AAGAGGCC | CAAGCAGAAGACGGCATACGAGATggcctcttGTGACTGGAGTTCAG |
| GGGTGTAT | CAAGCAGAAGACGGCATACGAGATatacacccGTGACTGGAGTTCAG |
| GCTGAGCA | CAAGCAGAAGACGGCATACGAGATtgctcagcGTGACTGGAGTTCAG |
| AGGAAAAG | CAAGCAGAAGACGGCATACGAGATcttttcctGTGACTGGAGTTCAGA |
| CAGGAGCA | CAAGCAGAAGACGGCATACGAGATtgctcctgGTGACTGGAGTTCAG |
| GACCCTGC | CAAGCAGAAGACGGCATACGAGATgcagggtcGTGACTGGAGTTCA |
| AGAGTCAA | CAAGCAGAAGACGGCATACGAGATttgactctGTGACTGGAGTTCAGA |
| CACATCAG | CAAGCAGAAGACGGCATACGAGATctgatgtgGTGACTGGAGTTCAG |
| AAGAGCGT | CAAGCAGAAGACGGCATACGAGATacgctcttGTGACTGGAGTTCAG |
| TGTGGTGC | CAAGCAGAAGACGGCATACGAGATgcaccacaGTGACTGGAGTTCA |
| GTTTTTCC | CAAGCAGAAGACGGCATACGAGATggaaaaacGTGACTGGAGTTCA |
| GTAGTTTT | CAAGCAGAAGACGGCATACGAGATaaaactacGTGACTGGAGTTCA |
| TTGTAACC | CAAGCAGAAGACGGCATACGAGATggttacaaGTGACTGGAGTTCAG |
| TATCTCCT | CAAGCAGAAGACGGCATACGAGATaggagataGTGACTGGAGTTCA |
| GATCTATA | CAAGCAGAAGACGGCATACGAGATtatagatcGTGACTGGAGTTCAG |
| TCACACAG | CAAGCAGAAGACGGCATACGAGATctgtgtgaGTGACTGGAGTTCAG |
| CAATTAGT | CAAGCAGAAGACGGCATACGAGATactaattgGTGACTGGAGTTCAG |
| ATGTCGCA | CAAGCAGAAGACGGCATACGAGATtgcgacatGTGACTGGAGTTCAG |
| CCTTACCA | CAAGCAGAAGACGGCATACGAGATtggtaaggGTGACTGGAGTTCA |
| AAGCCGTC | CAAGCAGAAGACGGCATACGAGATgacggcttGTGACTGGAGTTCAG |
| GGACCTAA | CAAGCAGAAGACGGCATACGAGATttaggtccGTGACTGGAGTTCAG |
| GATATCAT | CAAGCAGAAGACGGCATACGAGATatgatatcGTGACTGGAGTTCAG |
| GTAATATT | CAAGCAGAAGACGGCATACGAGATaatattacGTGACTGGAGTTCAG |
| TCAAATTC | CAAGCAGAAGACGGCATACGAGATgaatttgaGTGACTGGAGTTCAG |
| GGCCTACG | CAAGCAGAAGACGGCATACGAGATcgtaggccGTGACTGGAGTTCA |
| CCTGAGTA | CAAGCAGAAGACGGCATACGAGATtactcaggGTGACTGGAGTTCAG |
| ACATATCC | CAAGCAGAAGACGGCATACGAGATggatatgtGTGACTGGAGTTCAG |
| GTCCGAGC | CAAGCAGAAGACGGCATACGAGATgctcggacGTGACTGGAGTTCA |
| CCAGGTTC | CAAGCAGAAGACGGCATACGAGATgaacctggGTGACTGGAGTTCA |
| ACGATCTG | CAAGCAGAAGACGGCATACGAGATcagatcgtGTGACTGGAGTTCAG |
| TGCTGGGG | CAAGCAGAAGACGGCATACGAGATccccagcaGTGACTGGAGTTCA |
| GAGTCGCA | CAAGCAGAAGACGGCATACGAGATtgcgactcGTGACTGGAGTTCAG |
| CGCTAACG | CAAGCAGAAGACGGCATACGAGATcgttagcgGTGACTGGAGTTCAG |
| GTATCTTG | CAAGCAGAAGACGGCATACGAGATcaagatacGTGACTGGAGTTCA |
| ATCCAAAC | CAAGCAGAAGACGGCATACGAGATgtttggatGTGACTGGAGTTCAG |
| GTAATGCT | CAAGCAGAAGACGGCATACGAGATagcattacGTGACTGGAGTTCAG |
| TGACACCA | CAAGCAGAAGACGGCATACGAGATtggtgtcaGTGACTGGAGTTCAG |
| GCTCGGCG | CAAGCAGAAGACGGCATACGAGATcgccgagcGTGACTGGAGTTCA |
| GGCCTCTA | CAAGCAGAAGACGGCATACGAGATtagaggccGTGACTGGAGTTCA |
| GGTGAATG | CAAGCAGAAGACGGCATACGAGATcattcaccGTGACTGGAGTTCAG |
| CATGCTCT | CAAGCAGAAGACGGCATACGAGATagagcatgGTGACTGGAGTTCA |
| TGCGTAAC | CAAGCAGAAGACGGCATACGAGATgttacgcaGTGACTGGAGTTCAG |
| TTTCGGAC | CAAGCAGAAGACGGCATACGAGATgtccgaaaGTGACTGGAGTTCA |
| TACATCTA | CAAGCAGAAGACGGCATACGAGATtagatgtaGTGACTGGAGTTCAG |
| AGGCAATT | CAAGCAGAAGACGGCATACGAGATaattgcctGTGACTGGAGTTCAG |
| GGAGCAGT | CAAGCAGAAGACGGCATACGAGATactgctccGTGACTGGAGTTCAG |

| | |
|---|---|
| TAAAACAA | CAAGCAGAAGACGGCATACGAGATttgttttaGTGACTGGAGTTCAGA |
| CTAGTTGT | CAAGCAGAAGACGGCATACGAGATacaactagGTGACTGGAGTTCA |
| ACTATGCA | CAAGCAGAAGACGGCATACGAGATtgcatagtGTGACTGGAGTTCAG |
| GGGCCCGA | CAAGCAGAAGACGGCATACGAGATtcgggcccGTGACTGGAGTTCAG |
| TCGGGTTT | CAAGCAGAAGACGGCATACGAGATaaacccgaGTGACTGGAGTTCA |
| CGATTAGG | CAAGCAGAAGACGGCATACGAGATcctaatcgGTGACTGGAGTTCAG |
| ATAATGTA | CAAGCAGAAGACGGCATACGAGATtacattatGTGACTGGAGTTCAG |
| ATGTGGCC | CAAGCAGAAGACGGCATACGAGATggccacatGTGACTGGAGTTCA |
| GGCAAACC | CAAGCAGAAGACGGCATACGAGATggtttgccGTGACTGGAGTTCAG |
| CCCCTCGG | CAAGCAGAAGACGGCATACGAGATccgaggggGTGACTGGAGTTCA |
| GGTACCCA | CAAGCAGAAGACGGCATACGAGATtgggtaccGTGACTGGAGTTCAG |
| TCTTACGC | CAAGCAGAAGACGGCATACGAGATgcgtaagaGTGACTGGAGTTCA |
| GGTGCGCG | CAAGCAGAAGACGGCATACGAGATcgcgcaccGTGACTGGAGTTCA |
| CGCGTTAA | CAAGCAGAAGACGGCATACGAGATttaacgcgGTGACTGGAGTTCAG |
| GCCAGAGA | CAAGCAGAAGACGGCATACGAGATtctctggcGTGACTGGAGTTCAG |
| GAACTTAA | CAAGCAGAAGACGGCATACGAGATttaagttcGTGACTGGAGTTCAG |
| TCGCCCTA | CAAGCAGAAGACGGCATACGAGATtagggcgaGTGACTGGAGTTCA |
| CCAAGCAG | CAAGCAGAAGACGGCATACGAGATctgcttggGTGACTGGAGTTCAG |
| AGGGGTTT | CAAGCAGAAGACGGCATACGAGATaaacccctGTGACTGGAGTTCAG |
| GGTCGTCA | CAAGCAGAAGACGGCATACGAGATtgacgaccGTGACTGGAGTTCAG |
| GTTCGGCC | CAAGCAGAAGACGGCATACGAGATggccgaacGTGACTGGAGTTCA |
| AGTGATAC | CAAGCAGAAGACGGCATACGAGATgtatcactGTGACTGGAGTTCAG |
| AGGAGATG | CAAGCAGAAGACGGCATACGAGATcatctcctGTGACTGGAGTTCAG |
| ACACAGAA | CAAGCAGAAGACGGCATACGAGATttctgtgtGTGACTGGAGTTCAGA |
| ACCGAACG | CAAGCAGAAGACGGCATACGAGATcgttcggtGTGACTGGAGTTCAG |
| CCCGCATT | CAAGCAGAAGACGGCATACGAGATaatgcgggGTGACTGGAGTTCA |
| ATTCTTAA | CAAGCAGAAGACGGCATACGAGATttaagaatGTGACTGGAGTTCAG |
| GTTTTCGA | CAAGCAGAAGACGGCATACGAGATtcgaaaacGTGACTGGAGTTCA |
| CAACGGAT | CAAGCAGAAGACGGCATACGAGATatccgttgGTGACTGGAGTTCAG |
| AACTTCCA | CAAGCAGAAGACGGCATACGAGATtggaagttGTGACTGGAGTTCAG |
| AGCCTTAG | CAAGCAGAAGACGGCATACGAGATctaaggctGTGACTGGAGTTCAG |
| CAGTAGGG | CAAGCAGAAGACGGCATACGAGATccctactgGTGACTGGAGTTCAG |
| TTTAATCC | CAAGCAGAAGACGGCATACGAGATggattaaaGTGACTGGAGTTCA |
| CTAGGCGT | CAAGCAGAAGACGGCATACGAGATacgcctagGTGACTGGAGTTCA |
| CCCTGAGT | CAAGCAGAAGACGGCATACGAGATactcagggGTGACTGGAGTTCA |
| TTCCCTTC | CAAGCAGAAGACGGCATACGAGATgaagggaaGTGACTGGAGTTCA |
| AGTGACCG | CAAGCAGAAGACGGCATACGAGATcggtcactGTGACTGGAGTTCAG |
| GCGCTCCG | CAAGCAGAAGACGGCATACGAGATcggagcgcGTGACTGGAGTTCA |
| GAGCGCAT | CAAGCAGAAGACGGCATACGAGATatgcgctcGTGACTGGAGTTCAG |
| AGTAAACC | CAAGCAGAAGACGGCATACGAGATggtttactGTGACTGGAGTTCAG |
| CTCCACAG | CAAGCAGAAGACGGCATACGAGATctgtggagGTGACTGGAGTTCAG |
| GTTAAAAG | CAAGCAGAAGACGGCATACGAGATcttttaacGTGACTGGAGTTCAG |
| GAATATCG | CAAGCAGAAGACGGCATACGAGATcgatattcGTGACTGGAGTTCAG |
| GTGTCAGC | CAAGCAGAAGACGGCATACGAGATgctgacacGTGACTGGAGTTCA |
| ACACGAAA | CAAGCAGAAGACGGCATACGAGATtttcgtgtGTGACTGGAGTTCAGA |
| CTATGGAA | CAAGCAGAAGACGGCATACGAGATttccatagGTGACTGGAGTTCAG |
| GAAGGACA | CAAGCAGAAGACGGCATACGAGATtgtccttcGTGACTGGAGTTCAGA |
| GTTAGGGC | CAAGCAGAAGACGGCATACGAGATgccctaacGTGACTGGAGTTCAG |
| TATGGCAG | CAAGCAGAAGACGGCATACGAGATctgccataGTGACTGGAGTTCAG |
| AGCCGAAT | CAAGCAGAAGACGGCATACGAGATattcggctGTGACTGGAGTTCAG |
| TCGCGGCG | CAAGCAGAAGACGGCATACGAGATcgccgcgaGTGACTGGAGTTCA |
| TGTGCTTA | CAAGCAGAAGACGGCATACGAGATtaagcacaGTGACTGGAGTTCA |
| ATTATCGA | CAAGCAGAAGACGGCATACGAGATtcgataatGTGACTGGAGTTCAG |

| | |
|---|---|
| CGCAGCAG | CAAGCAGAAGACGGCATACGAGATctgctgcgGTGACTGGAGTTCAG |
| TCGGAGGG | CAAGCAGAAGACGGCATACGAGATccctccgaGTGACTGGAGTTCAG |
| CTACGTTA | CAAGCAGAAGACGGCATACGAGATtaacgtagGTGACTGGAGTTCAG |
| ATTAAGTA | CAAGCAGAAGACGGCATACGAGATtacttaatGTGACTGGAGTTCAGA |
| AGGTCCGC | CAAGCAGAAGACGGCATACGAGATgcggacctGTGACTGGAGTTCAG |
| CGAGTCAT | CAAGCAGAAGACGGCATACGAGATatgactcgGTGACTGGAGTTCAG |
| CTACAATA | CAAGCAGAAGACGGCATACGAGATtattgtagGTGACTGGAGTTCAGA |
| GATGGAAG | CAAGCAGAAGACGGCATACGAGATcttccatcGTGACTGGAGTTCAGA |
| AGCCTTGT | CAAGCAGAAGACGGCATACGAGATacaaggctGTGACTGGAGTTCAG |
| GATCTGAC | CAAGCAGAAGACGGCATACGAGATgtcagatcGTGACTGGAGTTCAG |
| CTAATTTA | CAAGCAGAAGACGGCATACGAGATtaaattagGTGACTGGAGTTCAG |
| TTTGGAGA | CAAGCAGAAGACGGCATACGAGATtctccaaaGTGACTGGAGTTCAG |
| ATGGCGGG | CAAGCAGAAGACGGCATACGAGATcccgccatGTGACTGGAGTTCAG |
| CGGGATTT | CAAGCAGAAGACGGCATACGAGATaaatcccgGTGACTGGAGTTCAG |
| GCCTAATA | CAAGCAGAAGACGGCATACGAGATtattaggcGTGACTGGAGTTCAG |
| AGAATATC | CAAGCAGAAGACGGCATACGAGATgatattctGTGACTGGAGTTCAGA |
| GCGAGAAG | CAAGCAGAAGACGGCATACGAGATcttctcgcGTGACTGGAGTTCAGA |
| AGAACACA | CAAGCAGAAGACGGCATACGAGATtgtgttctGTGACTGGAGTTCAGA |
| TCGTATAT | CAAGCAGAAGACGGCATACGAGATatatacgaGTGACTGGAGTTCAG |
| AGCCTATA | CAAGCAGAAGACGGCATACGAGATtataggctGTGACTGGAGTTCAG |
| GAGTTAGT | CAAGCAGAAGACGGCATACGAGATactaactcGTGACTGGAGTTCAG |
| TCATTAAC | CAAGCAGAAGACGGCATACGAGATgttaatgaGTGACTGGAGTTCAG |
| GGTGTGCA | CAAGCAGAAGACGGCATACGAGATtgcacaccGTGACTGGAGTTCAG |
| GAGGTAAG | CAAGCAGAAGACGGCATACGAGATcttacctcGTGACTGGAGTTCAGA |
| TACAGGAT | CAAGCAGAAGACGGCATACGAGATatcctgtaGTGACTGGAGTTCAGA |
| TATTTATC | CAAGCAGAAGACGGCATACGAGATgataaataGTGACTGGAGTTCAG |
| CGTAACTT | CAAGCAGAAGACGGCATACGAGATaagttacgGTGACTGGAGTTCAG |
| AATCTACT | CAAGCAGAAGACGGCATACGAGATagtagattGTGACTGGAGTTCAG |
| AGAAAGCG | CAAGCAGAAGACGGCATACGAGATcgctttctGTGACTGGAGTTCAGA |
| GAGCCCAA | CAAGCAGAAGACGGCATACGAGATttgggctcGTGACTGGAGTTCAGA |
| GCTTACGA | CAAGCAGAAGACGGCATACGAGATtcgtaagcGTGACTGGAGTTCAG |
| CCAATCCG | CAAGCAGAAGACGGCATACGAGATcggattggGTGACTGGAGTTCAG |
| TCCACTAA | CAAGCAGAAGACGGCATACGAGATttagtggaGTGACTGGAGTTCAG |
| AGTACCTT | CAAGCAGAAGACGGCATACGAGATaaggtactGTGACTGGAGTTCAG |
| TATTCACT | CAAGCAGAAGACGGCATACGAGATagtgaataGTGACTGGAGTTCAG |
| GGCCTGTT | CAAGCAGAAGACGGCATACGAGATaacaggccGTGACTGGAGTTCAG |
| GCGCAAGG | CAAGCAGAAGACGGCATACGAGATccttgcgcGTGACTGGAGTTCAG |
| CTGGGCTT | CAAGCAGAAGACGGCATACGAGATaagcccagGTGACTGGAGTTCAG |
| ATACTCTG | CAAGCAGAAGACGGCATACGAGATcagagtatGTGACTGGAGTTCAG |
| GGTCCCGC | CAAGCAGAAGACGGCATACGAGATgcgggaccGTGACTGGAGTTCAG |
| GCGGCTCA | CAAGCAGAAGACGGCATACGAGATtgagccgcGTGACTGGAGTTCAG |
| ATTAAGGG | CAAGCAGAAGACGGCATACGAGATcccttaatGTGACTGGAGTTCAGA |
| GTTTGTAC | CAAGCAGAAGACGGCATACGAGATgtacaaacGTGACTGGAGTTCAG |
| CTCCTACC | CAAGCAGAAGACGGCATACGAGATggtaggagGTGACTGGAGTTCAG |
| GCATTTCC | CAAGCAGAAGACGGCATACGAGATggaaatgcGTGACTGGAGTTCAG |
| TAAAGGTT | CAAGCAGAAGACGGCATACGAGATaacctttaGTGACTGGAGTTCAGA |
| TCACCGTG | CAAGCAGAAGACGGCATACGAGATcacggtgaGTGACTGGAGTTCAG |
| TCCTTGTT | CAAGCAGAAGACGGCATACGAGATaacaaggaGTGACTGGAGTTCA |
| AGGCTTCA | CAAGCAGAAGACGGCATACGAGATtgaagcctGTGACTGGAGTTCAG |
| AGTACAAA | CAAGCAGAAGACGGCATACGAGATtttgtactGTGACTGGAGTTCAGA |
| TCATCCAC | CAAGCAGAAGACGGCATACGAGATgtggatgaGTGACTGGAGTTCAG |
| CACGTTAC | CAAGCAGAAGACGGCATACGAGATgtaacgtgGTGACTGGAGTTCAG |
| GCGGAGCT | CAAGCAGAAGACGGCATACGAGATagctccgcGTGACTGGAGTTCAG |

| | |
|---|---|
| CTCGGGCA | CAAGCAGAAGACGGCATACGAGATtgcccgagGTGACTGGAGTTCAG |
| ATTACTTT | CAAGCAGAAGACGGCATACGAGATaaagtaatGTGACTGGAGTTCAG |
| CCAATATG | CAAGCAGAAGACGGCATACGAGATcatattggGTGACTGGAGTTCAG |
| GGCGCTGC | CAAGCAGAAGACGGCATACGAGATgcagcgccGTGACTGGAGTTCAG |
| TACTCTCA | CAAGCAGAAGACGGCATACGAGATtgagagtaGTGACTGGAGTTCAG |
| TCATCTCA | CAAGCAGAAGACGGCATACGAGATtgagatgaGTGACTGGAGTTCAG |
| AGTCCCCT | CAAGCAGAAGACGGCATACGAGATaggggactGTGACTGGAGTTCAG |
| AAGTTGCC | CAAGCAGAAGACGGCATACGAGATggcaacttGTGACTGGAGTTCAG |
| CCTATGTA | CAAGCAGAAGACGGCATACGAGATtacataggGTGACTGGAGTTCAG |
| TTATATTC | CAAGCAGAAGACGGCATACGAGATgaatataaGTGACTGGAGTTCAG |
| AAGGTTTC | CAAGCAGAAGACGGCATACGAGATgaaaccttGTGACTGGAGTTCAG |
| GCTGGGGG | CAAGCAGAAGACGGCATACGAGATcccccagcGTGACTGGAGTTCAG |
| CCTAGGAA | CAAGCAGAAGACGGCATACGAGATttcctaggGTGACTGGAGTTCAGA |
| CAGGAGAC | CAAGCAGAAGACGGCATACGAGATgtctcctgGTGACTGGAGTTCAGA |
| TGATTTGC | CAAGCAGAAGACGGCATACGAGATgcaaatcaGTGACTGGAGTTCAG |
| GATCTCTC | CAAGCAGAAGACGGCATACGAGATgagagatcGTGACTGGAGTTCAG |
| AAAAATTG | CAAGCAGAAGACGGCATACGAGATcaatttttGTGACTGGAGTTCAGA |
| AACAGGAC | CAAGCAGAAGACGGCATACGAGATgtcctgttGTGACTGGAGTTCAGA |
| GTTACACT | CAAGCAGAAGACGGCATACGAGATagtgtaacGTGACTGGAGTTCAG |
| CAAATCGT | CAAGCAGAAGACGGCATACGAGATacgatttgGTGACTGGAGTTCAG |
| TCGTGCCG | CAAGCAGAAGACGGCATACGAGATcggcacgaGTGACTGGAGTTCAG |
| TTTTAGCC | CAAGCAGAAGACGGCATACGAGATggctaaaaGTGACTGGAGTTCAG |
| CTGTTTCG | CAAGCAGAAGACGGCATACGAGATcgaaacagGTGACTGGAGTTCA |
| TTAGCAAC | CAAGCAGAAGACGGCATACGAGATgttgctaaGTGACTGGAGTTCAG |
| AGGACTGC | CAAGCAGAAGACGGCATACGAGATgcagtcctGTGACTGGAGTTCAG |
| CTTAAAGG | CAAGCAGAAGACGGCATACGAGATcctttaagGTGACTGGAGTTCAGA |
| ACACAACC | CAAGCAGAAGACGGCATACGAGATggttgtgtGTGACTGGAGTTCAGA |
| AACTAACG | CAAGCAGAAGACGGCATACGAGATcgttagttGTGACTGGAGTTCAGA |
| TATGCGAG | CAAGCAGAAGACGGCATACGAGATctcgcataGTGACTGGAGTTCAG |
| TGTTGACC | CAAGCAGAAGACGGCATACGAGATggtcaacaGTGACTGGAGTTCAG |
| AACTACTA | CAAGCAGAAGACGGCATACGAGATtagtagttGTGACTGGAGTTCAGA |
| TTAGATGC | CAAGCAGAAGACGGCATACGAGATgcatctaaGTGACTGGAGTTCAG |
| AATCCTTA | CAAGCAGAAGACGGCATACGAGATtaaggattGTGACTGGAGTTCAG |
| AGAGCCAC | CAAGCAGAAGACGGCATACGAGATgtggctctGTGACTGGAGTTCAGA |
| GACTCTAC | CAAGCAGAAGACGGCATACGAGATgtagagtcGTGACTGGAGTTCAG |
| TAAGCATA | CAAGCAGAAGACGGCATACGAGATtatgcttaGTGACTGGAGTTCAGA |
| GCAGCAAT | CAAGCAGAAGACGGCATACGAGATattgctgcGTGACTGGAGTTCAGA |
| GACCCGCA | CAAGCAGAAGACGGCATACGAGATtgcgggtcGTGACTGGAGTTCAG |
| CCCAATGG | CAAGCAGAAGACGGCATACGAGATccattgggGTGACTGGAGTTCAG |
| TCGAGAAA | CAAGCAGAAGACGGCATACGAGATtttctcgaGTGACTGGAGTTCAGA |
| GCTCAATC | CAAGCAGAAGACGGCATACGAGATgattgagcGTGACTGGAGTTCAG |
| ACCGGGAG | CAAGCAGAAGACGGCATACGAGATctcccggtGTGACTGGAGTTCAG |
| AGTTTCTT | CAAGCAGAAGACGGCATACGAGATaagaaactGTGACTGGAGTTCAG |
| ACCGTGCA | CAAGCAGAAGACGGCATACGAGATtgcacggtGTGACTGGAGTTCAG |
| TGAGTAAA | CAAGCAGAAGACGGCATACGAGATtttactcaGTGACTGGAGTTCAGA |
| TTAAATCT | CAAGCAGAAGACGGCATACGAGATagatttaaGTGACTGGAGTTCAG |
| ACATGTTC | CAAGCAGAAGACGGCATACGAGATgaacatgtGTGACTGGAGTTCAG |
| GCGTGAAA | CAAGCAGAAGACGGCATACGAGATtttcacgcGTGACTGGAGTTCAGA |
| GAAATGCG | CAAGCAGAAGACGGCATACGAGATcgcatttcGTGACTGGAGTTCAGA |
| AAGTTCCG | CAAGCAGAAGACGGCATACGAGATcggaacttGTGACTGGAGTTCAG |
| TGTCGCCC | CAAGCAGAAGACGGCATACGAGATgggcgacaGTGACTGGAGTTCA |
| TTAAATAC | CAAGCAGAAGACGGCATACGAGATgtatttaaGTGACTGGAGTTCAGA |
| GATAGAGA | CAAGCAGAAGACGGCATACGAGATtctctatcGTGACTGGAGTTCAGA |

| | |
|---|---|
| GTTTCTTC | CAAGCAGAAGACGGCATACGAGATgaagaaacGTGACTGGAGTTCA |
| TCCCGAGC | CAAGCAGAAGACGGCATACGAGATgctcgggaGTGACTGGAGTTCA |
| CGATCTCA | CAAGCAGAAGACGGCATACGAGATtgagatcgGTGACTGGAGTTCAG |
| GGTTGATC | CAAGCAGAAGACGGCATACGAGATgatcaaccGTGACTGGAGTTCA |
| AATAATCA | CAAGCAGAAGACGGCATACGAGATtgattattGTGACTGGAGTTCAGA |
| GGTGTTTT | CAAGCAGAAGACGGCATACGAGATaaaacaccGTGACTGGAGTTCA |
| GCACGCGA | CAAGCAGAAGACGGCATACGAGATtcgcgtgcGTGACTGGAGTTCAG |
| TCCCTGCA | CAAGCAGAAGACGGCATACGAGATtgcagggaGTGACTGGAGTTCA |
| TAGGCGTC | CAAGCAGAAGACGGCATACGAGATgacgcctaGTGACTGGAGTTCA |
| ACCACCGT | CAAGCAGAAGACGGCATACGAGATacggtggtGTGACTGGAGTTCA |
| TTCCACAT | CAAGCAGAAGACGGCATACGAGATatgtggaaGTGACTGGAGTTCA |
| TCGTCAAA | CAAGCAGAAGACGGCATACGAGATtttgacgaGTGACTGGAGTTCAG |
| CCGCTTGA | CAAGCAGAAGACGGCATACGAGATtcaagcggGTGACTGGAGTTCA |
| CTAAGAAG | CAAGCAGAAGACGGCATACGAGATcttcttagGTGACTGGAGTTCAG |
| ATGACATT | CAAGCAGAAGACGGCATACGAGATaatgtcatGTGACTGGAGTTCAG |
| CTGTAGAA | CAAGCAGAAGACGGCATACGAGATttctacagGTGACTGGAGTTCAG |
| GCTTGCTG | CAAGCAGAAGACGGCATACGAGATcagcaagcGTGACTGGAGTTCA |
| ACGTCAGA | CAAGCAGAAGACGGCATACGAGATtctgacgtGTGACTGGAGTTCAG |
| ACCCACTC | CAAGCAGAAGACGGCATACGAGATgagtgggtGTGACTGGAGTTCA |
| AACACTAG | CAAGCAGAAGACGGCATACGAGATctagtgttGTGACTGGAGTTCAG |
| AATGAACA | CAAGCAGAAGACGGCATACGAGATtgttcattGTGACTGGAGTTCAGA |
| GGCAGTAT | CAAGCAGAAGACGGCATACGAGATatactgccGTGACTGGAGTTCAG |
| ACCACACG | CAAGCAGAAGACGGCATACGAGATcgtgtggtGTGACTGGAGTTCAG |
| GCACAGCC | CAAGCAGAAGACGGCATACGAGATggctgtgcGTGACTGGAGTTCAG |
| GCCTTTGA | CAAGCAGAAGACGGCATACGAGATtcaaaggcGTGACTGGAGTTCA |
| GACTGTGG | CAAGCAGAAGACGGCATACGAGATccacagtcGTGACTGGAGTTCA |
| AAACAAGC | CAAGCAGAAGACGGCATACGAGATgcttgtttGTGACTGGAGTTCAGA |
| CCTCGTAA | CAAGCAGAAGACGGCATACGAGATttacgaggGTGACTGGAGTTCA |
| AAATCGGA | CAAGCAGAAGACGGCATACGAGATtccgatttGTGACTGGAGTTCAG |
| ACGGAGAC | CAAGCAGAAGACGGCATACGAGATgtctccgtGTGACTGGAGTTCAG |
| GTGCGCCG | CAAGCAGAAGACGGCATACGAGATcggcgcacGTGACTGGAGTTCA |
| TGGGCATT | CAAGCAGAAGACGGCATACGAGATaatgcccaGTGACTGGAGTTCA |
| AGGGAACG | CAAGCAGAAGACGGCATACGAGATcgttccctGTGACTGGAGTTCAG |
| CTAGTGGC | CAAGCAGAAGACGGCATACGAGATgccactagGTGACTGGAGTTCA |
| TTATAGGC | CAAGCAGAAGACGGCATACGAGATgcctataaGTGACTGGAGTTCAG |
| ATCATTAC | CAAGCAGAAGACGGCATACGAGATgtaatgatGTGACTGGAGTTCAG |
| CTGCTTTG | CAAGCAGAAGACGGCATACGAGATcaaagcagGTGACTGGAGTTCA |
| ACTTTTCC | CAAGCAGAAGACGGCATACGAGATggaaaagtGTGACTGGAGTTCA |
| CGGCTCTC | CAAGCAGAAGACGGCATACGAGATgagagccgGTGACTGGAGTTCA |
| CTGTCAAG | CAAGCAGAAGACGGCATACGAGATcttgacagGTGACTGGAGTTCAG |
| TCGCTAGA | CAAGCAGAAGACGGCATACGAGATtctagcgaGTGACTGGAGTTCAG |
| AAGGCGAT | CAAGCAGAAGACGGCATACGAGATatcgccttGTGACTGGAGTTCAG |
| TTGCAGTG | CAAGCAGAAGACGGCATACGAGATcactgcaaGTGACTGGAGTTCA |
| ACAGACGC | CAAGCAGAAGACGGCATACGAGATgcgtctgtGTGACTGGAGTTCAG |
| GGAATAAG | CAAGCAGAAGACGGCATACGAGATcttattccGTGACTGGAGTTCAGA |
| GCCTAGTT | CAAGCAGAAGACGGCATACGAGATaactaggcGTGACTGGAGTTCA |
| GACGCCAT | CAAGCAGAAGACGGCATACGAGATatggcgtcGTGACTGGAGTTCAG |
| ATGCTGCA | CAAGCAGAAGACGGCATACGAGATtgcagcatGTGACTGGAGTTCAG |
| CCAACCGG | CAAGCAGAAGACGGCATACGAGATccggttggGTGACTGGAGTTCAG |
| ACTAAGGC | CAAGCAGAAGACGGCATACGAGATgccttagtGTGACTGGAGTTCAG |
| CGCGCACT | CAAGCAGAAGACGGCATACGAGATagtgcgcgGTGACTGGAGTTCA |
| CCGCCAAA | CAAGCAGAAGACGGCATACGAGATtttggcggGTGACTGGAGTTCAG |
| ACCTCTCG | CAAGCAGAAGACGGCATACGAGATcgagaggtGTGACTGGAGTTCA |

| | |
|---|---|
| TGCAAGGC | CAAGCAGAAGACGGCATACGAGATgccttgcaGTGACTGGAGTTCAG |
| CCTACGGA | CAAGCAGAAGACGGCATACGAGATtccgtaggGTGACTGGAGTTCAG |
| GCGCAAAC | CAAGCAGAAGACGGCATACGAGATgtttgcgcGTGACTGGAGTTCAG |
| TGGCCTAC | CAAGCAGAAGACGGCATACGAGATgtaggccaGTGACTGGAGTTCA |
| ATTGCACC | CAAGCAGAAGACGGCATACGAGATggtgcaatGTGACTGGAGTTCAG |
| ACCTCGGA | CAAGCAGAAGACGGCATACGAGATtccgaggtGTGACTGGAGTTCAG |
| ACAAAGTG | CAAGCAGAAGACGGCATACGAGATcactttgtGTGACTGGAGTTCAG |
| TGAATCTA | CAAGCAGAAGACGGCATACGAGATtagattcaGTGACTGGAGTTCAG |
| TCGACGAA | CAAGCAGAAGACGGCATACGAGATttcgtcgaGTGACTGGAGTTCAG |
| AAGTAACA | CAAGCAGAAGACGGCATACGAGATtgttacttGTGACTGGAGTTCAGA |
| ATCAGCCT | CAAGCAGAAGACGGCATACGAGATaggctgatGTGACTGGAGTTCAG |
| ATCAGTCA | CAAGCAGAAGACGGCATACGAGATtgactgatGTGACTGGAGTTCAG |
| AAACGACA | CAAGCAGAAGACGGCATACGAGATtgtcgtttGTGACTGGAGTTCAGA |
| TATTAGCG | CAAGCAGAAGACGGCATACGAGATcgctaataGTGACTGGAGTTCAG |
| ATGTGTTT | CAAGCAGAAGACGGCATACGAGATaaacacatGTGACTGGAGTTCA |
| TGAGATTA | CAAGCAGAAGACGGCATACGAGATtaatctcaGTGACTGGAGTTCAG |
| ATTGATTG | CAAGCAGAAGACGGCATACGAGATcaatcaatGTGACTGGAGTTCAG |
| CTACCAAT | CAAGCAGAAGACGGCATACGAGATattggtagGTGACTGGAGTTCAG |
| GTGCCCGT | CAAGCAGAAGACGGCATACGAGATacgggcacGTGACTGGAGTTCA |
| CTCAGCGG | CAAGCAGAAGACGGCATACGAGATccgctgagGTGACTGGAGTTCAG |
| TTCCATTT | CAAGCAGAAGACGGCATACGAGATaaatggaaGTGACTGGAGTTCA |
| TTAGGGTG | CAAGCAGAAGACGGCATACGAGATcaccctaaGTGACTGGAGTTCAG |
| TGCGGCCA | CAAGCAGAAGACGGCATACGAGATtggccgcaGTGACTGGAGTTCA |
| GGGACGAA | CAAGCAGAAGACGGCATACGAGATttcgtcccGTGACTGGAGTTCAG |
| AAGACGAC | CAAGCAGAAGACGGCATACGAGATgtcgtcttGTGACTGGAGTTCAGA |
| CTGTCTAA | CAAGCAGAAGACGGCATACGAGATttagacagGTGACTGGAGTTCAG |
| ATGCCTTT | CAAGCAGAAGACGGCATACGAGATaaaggcatGTGACTGGAGTTCA |
| TAGACCTC | CAAGCAGAAGACGGCATACGAGATgaggtctaGTGACTGGAGTTCAG |
| CTCGACGG | CAAGCAGAAGACGGCATACGAGATccgtcgagGTGACTGGAGTTCA |
| TGTATACT | CAAGCAGAAGACGGCATACGAGATagtatacaGTGACTGGAGTTCAG |
| ACCTTCAA | CAAGCAGAAGACGGCATACGAGATttgaaggtGTGACTGGAGTTCAG |
| CCGTGGAT | CAAGCAGAAGACGGCATACGAGATatccacggGTGACTGGAGTTCA |
| GGTAGTGA | CAAGCAGAAGACGGCATACGAGATtcactaccGTGACTGGAGTTCAG |
| TTTAACCT | CAAGCAGAAGACGGCATACGAGATaggttaaaGTGACTGGAGTTCA |
| GTTCGATG | CAAGCAGAAGACGGCATACGAGATcatcgaacGTGACTGGAGTTCA |
| TGTGTTAC | CAAGCAGAAGACGGCATACGAGATgtaacacaGTGACTGGAGTTCA |
| GCAGATTA | CAAGCAGAAGACGGCATACGAGATtaatctgcGTGACTGGAGTTCAG |
| AGCTCGCG | CAAGCAGAAGACGGCATACGAGATcgcgagctGTGACTGGAGTTCA |
| TTTCAGCG | CAAGCAGAAGACGGCATACGAGATcgctgaaaGTGACTGGAGTTCA |
| GCGGGTAC | CAAGCAGAAGACGGCATACGAGATgtacccgcGTGACTGGAGTTCAG |
| ACCGTCTG | CAAGCAGAAGACGGCATACGAGATcagacggtGTGACTGGAGTTCA |
| GTCCAATG | CAAGCAGAAGACGGCATACGAGATcattggacGTGACTGGAGTTCAG |
| GGTTTCTA | CAAGCAGAAGACGGCATACGAGATtagaaaccGTGACTGGAGTTCA |
| TGGTTCCC | CAAGCAGAAGACGGCATACGAGATgggaaccaGTGACTGGAGTTCA |
| AAGTGGAC | CAAGCAGAAGACGGCATACGAGATgtccacttGTGACTGGAGTTCAG |
| CCACTGTC | CAAGCAGAAGACGGCATACGAGATgacagtggGTGACTGGAGTTCA |
| CATTCGGC | CAAGCAGAAGACGGCATACGAGATgccgaatgGTGACTGGAGTTCA |
| ATCTCGGT | CAAGCAGAAGACGGCATACGAGATaccgagatGTGACTGGAGTTCA |
| TTTTCGAA | CAAGCAGAAGACGGCATACGAGATttcgaaaaGTGACTGGAGTTCAG |
| ATGCGCAG | CAAGCAGAAGACGGCATACGAGATctgcgcatGTGACTGGAGTTCAG |
| AAGGAATC | CAAGCAGAAGACGGCATACGAGATgattccttGTGACTGGAGTTCAGA |
| ATAGCTCC | CAAGCAGAAGACGGCATACGAGATggagctatGTGACTGGAGTTCAG |
| GTTGCTAC | CAAGCAGAAGACGGCATACGAGATgtagcaacGTGACTGGAGTTCA |

| | |
|---|---|
| CCTATTAC | CAAGCAGAAGACGGCATACGAGATgtaataggGTGACTGGAGTTCAG |
| GCGCATCG | CAAGCAGAAGACGGCATACGAGATcgatgcgcGTGACTGGAGTTCAG |
| CTTATCTG | CAAGCAGAAGACGGCATACGAGATcagataagGTGACTGGAGTTCA |
| AAGGGATT | CAAGCAGAAGACGGCATACGAGATaatcccttGTGACTGGAGTTCAG |
| TGCTCCCC | CAAGCAGAAGACGGCATACGAGATggggagcaGTGACTGGAGTTCA |
| GATATTTT | CAAGCAGAAGACGGCATACGAGATaaaatatcGTGACTGGAGTTCAG |
| GAGTCCCG | CAAGCAGAAGACGGCATACGAGATcgggactcGTGACTGGAGTTCAG |
| TCCCTCAG | CAAGCAGAAGACGGCATACGAGATctgagggaGTGACTGGAGTTCA |
| AGGCGTGC | CAAGCAGAAGACGGCATACGAGATgcacgcctGTGACTGGAGTTCAG |
| TGCAGGTC | CAAGCAGAAGACGGCATACGAGATgacctgcaGTGACTGGAGTTCAG |
| TCGGTCTA | CAAGCAGAAGACGGCATACGAGATtagaccgaGTGACTGGAGTTCA |
| GTCTACAA | CAAGCAGAAGACGGCATACGAGATttgtagacGTGACTGGAGTTCAG |
| GACATCCG | CAAGCAGAAGACGGCATACGAGATcggatgtcGTGACTGGAGTTCAG |
| GCGCACAG | CAAGCAGAAGACGGCATACGAGATctgtgcgcGTGACTGGAGTTCAG |
| CCTAATTT | CAAGCAGAAGACGGCATACGAGATaaattaggGTGACTGGAGTTCAG |
| GGGCCTTC | CAAGCAGAAGACGGCATACGAGATgaaggcccGTGACTGGAGTTCA |
| AGGAGTGG | CAAGCAGAAGACGGCATACGAGATccactcctGTGACTGGAGTTCAG |
| CAATGATG | CAAGCAGAAGACGGCATACGAGATcatcattgGTGACTGGAGTTCAG |
| AGATCTGG | CAAGCAGAAGACGGCATACGAGATccagatctGTGACTGGAGTTCAG |
| CTAAATTT | CAAGCAGAAGACGGCATACGAGATaaatttagGTGACTGGAGTTCAG |
| CCTACTCG | CAAGCAGAAGACGGCATACGAGATcgagtaggGTGACTGGAGTTCA |
| AAAAGCAC | CAAGCAGAAGACGGCATACGAGATgtgcttttGTGACTGGAGTTCAGA |
| TGGCACAA | CAAGCAGAAGACGGCATACGAGATttgtgccaGTGACTGGAGTTCAG |
| TAATGATT | CAAGCAGAAGACGGCATACGAGATaatcattaGTGACTGGAGTTCAG |
| GGAACGGG | CAAGCAGAAGACGGCATACGAGATcccgttccGTGACTGGAGTTCAG |
| TGGTTACT | CAAGCAGAAGACGGCATACGAGATagtaaccaGTGACTGGAGTTCA |
| GGCAATTT | CAAGCAGAAGACGGCATACGAGATaaattgccGTGACTGGAGTTCAG |
| AGTCTTTG | CAAGCAGAAGACGGCATACGAGATcaaagactGTGACTGGAGTTCA |
| CAAGGAGA | CAAGCAGAAGACGGCATACGAGATtctccttgGTGACTGGAGTTCAGA |
| AGGCCCGG | CAAGCAGAAGACGGCATACGAGATccgggcctGTGACTGGAGTTCAG |
| ACGGCAGC | CAAGCAGAAGACGGCATACGAGATgctgccgtGTGACTGGAGTTCAG |
| GCCTCGGC | CAAGCAGAAGACGGCATACGAGATgccgaggcGTGACTGGAGTTCA |
| CAGACATG | CAAGCAGAAGACGGCATACGAGATcatgtctgGTGACTGGAGTTCAG |
| GCCGTGAG | CAAGCAGAAGACGGCATACGAGATctcacggcGTGACTGGAGTTCAG |
| TATGTTGT | CAAGCAGAAGACGGCATACGAGATacaacataGTGACTGGAGTTCA |
| TACGCACG | CAAGCAGAAGACGGCATACGAGATcgtgcgtaGTGACTGGAGTTCAG |
| GGCCGCTC | CAAGCAGAAGACGGCATACGAGATgagcggccGTGACTGGAGTTCA |
| CAACTGCG | CAAGCAGAAGACGGCATACGAGATcgcagttgGTGACTGGAGTTCAG |
| CAGAATTA | CAAGCAGAAGACGGCATACGAGATtaattctgGTGACTGGAGTTCAG |
| GGAAAGAT | CAAGCAGAAGACGGCATACGAGATatctttccGTGACTGGAGTTCAGA |

ble-stranded lig147 and lig200 from the ordered single-stranded oligos
ecessary sequences for the Illumina flowcell

INNNNNNNNNNNNNNNNNNNNNNNNNNNNNNNNNNNNNNNNNNNNNNNNNNNNNNNNNNNNNNNNNNNNNNNN
INNNNNNNNNNNNNNNNNNNNNNNNNNNNNNNNNNNNNNNNNNNNNNNNNNNNNNNNNNNNNNNNNNNNNNNN
**dom601 sequence)**
INNNNNNNNNNNNNNNNNNNNNNNNNNNNNNNNNNNNNNNNNNNNNNNNNNNNNNGGCCGCTCAATTGGTC

GCTCTTCC

GACGTGTGCTCTTCCG
ACGTGTGCTCTTCCG
ACGTGTGCTCTTCCG
ACGTGTGCTCTTCCG
GACGTGTGCTCTTCCG
ACGTGTGCTCTTCCG
ACGTGTGCTCTTCCG
GACGTGTGCTCTTCCG
ACGTGTGCTCTTCCG
GACGTGTGCTCTTCCG
CGTGTGCTCTTCCG
GACGTGTGCTCTTCCG
GACGTGTGCTCTTCCG
ACGTGTGCTCTTCCG
GACGTGTGCTCTTCCG
GACGTGTGCTCTTCCG
GACGTGTGCTCTTCCG
GACGTGTGCTCTTCCG
ACGTGTGCTCTTCCG
ACGTGTGCTCTTCCG
ACGTGTGCTCTTCCG
GACGTGTGCTCTTCCG
ACGTGTGCTCTTCCG
ACGTGTGCTCTTCCG
GACGTGTGCTCTTCCG
GACGTGTGCTCTTCCG
GACGTGTGCTCTTCCG
GACGTGTGCTCTTCCG
GACGTGTGCTCTTCCG
ACGTGTGCTCTTCCG
ACGTGTGCTCTTCCG
GACGTGTGCTCTTCCG
ACGTGTGCTCTTCCG
GACGTGTGCTCTTCCG
GACGTGTGCTCTTCCG
GACGTGTGCTCTTCCG
GACGTGTGCTCTTCCG
GACGTGTGCTCTTCCG
GACGTGTGCTCTTCCG

ACGTGTGCTCTTCCG
GACGTGTGCTCTTCCG
GACGTGTGCTCTTCCG
ACGTGTGCTCTTCCG
GACGTGTGCTCTTCCG
ACGTGTGCTCTTCCG
GACGTGTGCTCTTCCG
ACGTGTGCTCTTCCG
ACGTGTGCTCTTCCG
GACGTGTGCTCTTCCG
GACGTGTGCTCTTCCG
ACGTGTGCTCTTCCG
GACGTGTGCTCTTCCG
ACGTGTGCTCTTCCG
CGTGTGCTCTTCCG
GACGTGTGCTCTTCCG
ACGTGTGCTCTTCCG
GACGTGTGCTCTTCCG
ACGTGTGCTCTTCCG
ACGTGTGCTCTTCCG
GACGTGTGCTCTTCCG
GACGTGTGCTCTTCCG
GACGTGTGCTCTTCCG
CGTGTGCTCTTCCG
GACGTGTGCTCTTCCG
GACGTGTGCTCTTCCG
GACGTGTGCTCTTCCG
ACGTGTGCTCTTCCG
GACGTGTGCTCTTCCG
ACGTGTGCTCTTCCG
GACGTGTGCTCTTCCG
GACGTGTGCTCTTCCG
GACGTGTGCTCTTCCG
ACGTGTGCTCTTCCG
ACGTGTGCTCTTCCG
GACGTGTGCTCTTCCG
ACGTGTGCTCTTCCG
ACGTGTGCTCTTCCG
ACGTGTGCTCTTCCG
ACGTGTGCTCTTCCG
ACGTGTGCTCTTCCG
ACGTGTGCTCTTCCG
GACGTGTGCTCTTCCG
GACGTGTGCTCTTCCG
ACGTGTGCTCTTCCG
ACGTGTGCTCTTCCG
CGTGTGCTCTTCCG
ACGTGTGCTCTTCCG
ACGTGTGCTCTTCCG
ACGTGTGCTCTTCCG
ACGTGTGCTCTTCCG
CGTGTGCTCTTCCG
ACGTGTGCTCTTCCG

GACGTGTGCTCTTCCG
ACGTGTGCTCTTCCG
GACGTGTGCTCTTCCG
GACGTGTGCTCTTCCG
GACGTGTGCTCTTCCG
GACGTGTGCTCTTCCG
GACGTGTGCTCTTCCG
GACGTGTGCTCTTCCG
GACGTGTGCTCTTCCG
ACGTGTGCTCTTCCG
GACGTGTGCTCTTCCG
GACGTGTGCTCTTCCG
GACGTGTGCTCTTCCG
GACGTGTGCTCTTCCG
GACGTGTGCTCTTCCG
GACGTGTGCTCTTCCG
GACGTGTGCTCTTCCG
GACGTGTGCTCTTCCG
GACGTGTGCTCTTCCG
ACGTGTGCTCTTCCG
ACGTGTGCTCTTCCG
GACGTGTGCTCTTCCG
GACGTGTGCTCTTCCG
GACGTGTGCTCTTCCG
GACGTGTGCTCTTCCG
GACGTGTGCTCTTCCG
GACGTGTGCTCTTCCG
ACGTGTGCTCTTCCG
GACGTGTGCTCTTCCG
ACGTGTGCTCTTCCG
ACGTGTGCTCTTCCG
GACGTGTGCTCTTCCG
GACGTGTGCTCTTCCG
ACGTGTGCTCTTCCG
GACGTGTGCTCTTCCG
GACGTGTGCTCTTCCG
GACGTGTGCTCTTCCG
ACGTGTGCTCTTCCG
GACGTGTGCTCTTCCG
GACGTGTGCTCTTCCG
GACGTGTGCTCTTCCG
GACGTGTGCTCTTCCG
GACGTGTGCTCTTCCG
ACGTGTGCTCTTCCG
ACGTGTGCTCTTCCG
GACGTGTGCTCTTCCG
ACGTGTGCTCTTCCG
GACGTGTGCTCTTCCG
GACGTGTGCTCTTCCG
GACGTGTGCTCTTCCG
ACGTGTGCTCTTCCG
ACGTGTGCTCTTCCG
GACGTGTGCTCTTCCG

GACGTGTGCTCTTCCG
GACGTGTGCTCTTCCG
ACGTGTGCTCTTCCG
GACGTGTGCTCTTCCG
ACGTGTGCTCTTCCG
GACGTGTGCTCTTCCG
ACGTGTGCTCTTCCG
GACGTGTGCTCTTCCG
ACGTGTGCTCTTCCG
GACGTGTGCTCTTCCG
GACGTGTGCTCTTCCG
ACGTGTGCTCTTCCG
GACGTGTGCTCTTCCG
GACGTGTGCTCTTCCG
GACGTGTGCTCTTCCG
ACGTGTGCTCTTCCG
GACGTGTGCTCTTCCG
GACGTGTGCTCTTCCG
GACGTGTGCTCTTCCG
ACGTGTGCTCTTCCG
GACGTGTGCTCTTCCG
GACGTGTGCTCTTCCG
GACGTGTGCTCTTCCG
ACGTGTGCTCTTCCG
GACGTGTGCTCTTCCG
ACGTGTGCTCTTCCG
GACGTGTGCTCTTCCG
ACGTGTGCTCTTCCG
GACGTGTGCTCTTCCG
ACGTGTGCTCTTCCG
ACGTGTGCTCTTCCG
ACGTGTGCTCTTCCG
GACGTGTGCTCTTCCG
GACGTGTGCTCTTCCG
GACGTGTGCTCTTCCG
GACGTGTGCTCTTCCG
ACGTGTGCTCTTCCG
ACGTGTGCTCTTCCG
ACGTGTGCTCTTCCG
GACGTGTGCTCTTCCG
ACGTGTGCTCTTCCG
GACGTGTGCTCTTCCG
ACGTGTGCTCTTCCG
ACGTGTGCTCTTCCG
GACGTGTGCTCTTCCG
ACGTGTGCTCTTCCG
ACGTGTGCTCTTCCG
GACGTGTGCTCTTCCG
ACGTGTGCTCTTCCG
GACGTGTGCTCTTCCG
ACGTGTGCTCTTCCG
ACGTGTGCTCTTCCG
GACGTGTGCTCTTCCG

ACGTGTGCTCTTCCG
ACGTGTGCTCTTCCG
ACGTGTGCTCTTCCG
ACGTGTGCTCTTCCG
GACGTGTGCTCTTCCG
GACGTGTGCTCTTCCG
ACGTGTGCTCTTCCG
ACGTGTGCTCTTCCG
ACGTGTGCTCTTCCG
GACGTGTGCTCTTCCG
ACGTGTGCTCTTCCG
CGTGTGCTCTTCCG
GACGTGTGCTCTTCCG
GACGTGTGCTCTTCCG
GACGTGTGCTCTTCCG
GACGTGTGCTCTTCCG
GACGTGTGCTCTTCCG
ACGTGTGCTCTTCCG
GACGTGTGCTCTTCCG
GACGTGTGCTCTTCCG
ACGTGTGCTCTTCCG
ACGTGTGCTCTTCCG
ACGTGTGCTCTTCCG
ACGTGTGCTCTTCCG
GACGTGTGCTCTTCCG
GACGTGTGCTCTTCCG
GACGTGTGCTCTTCCG
GACGTGTGCTCTTCCG
GACGTGTGCTCTTCCG
GACGTGTGCTCTTCCG
GACGTGTGCTCTTCCG
CGTGTGCTCTTCCG
ACGTGTGCTCTTCCG
GACGTGTGCTCTTCCG
ACGTGTGCTCTTCCG
GACGTGTGCTCTTCCG
ACGTGTGCTCTTCCG
GACGTGTGCTCTTCCG
CGTGTGCTCTTCCG
ACGTGTGCTCTTCCG
CGTGTGCTCTTCCG
ACGTGTGCTCTTCCG
ACGTGTGCTCTTCCG
ACGTGTGCTCTTCCG
GACGTGTGCTCTTCCG
ACGTGTGCTCTTCCG
GACGTGTGCTCTTCCG
ACGTGTGCTCTTCCG
GACGTGTGCTCTTCCG
ACGTGTGCTCTTCCG
GACGTGTGCTCTTCCG
GACGTGTGCTCTTCCG
GACGTGTGCTCTTCCG

ACGTGTGCTCTTCCG
ACGTGTGCTCTTCCG
ACGTGTGCTCTTCCG
GACGTGTGCTCTTCCG
GACGTGTGCTCTTCCG
GACGTGTGCTCTTCCG
ACGTGTGCTCTTCCG
GACGTGTGCTCTTCCG
ACGTGTGCTCTTCCG
GACGTGTGCTCTTCCG
ACGTGTGCTCTTCCG
ACGTGTGCTCTTCCG
GACGTGTGCTCTTCCG
GACGTGTGCTCTTCCG
GACGTGTGCTCTTCCG
GACGTGTGCTCTTCCG
ACGTGTGCTCTTCCG
GACGTGTGCTCTTCCG
GACGTGTGCTCTTCCG
ACGTGTGCTCTTCCG
ACGTGTGCTCTTCCG
CGTGTGCTCTTCCG
ACGTGTGCTCTTCCG
ACGTGTGCTCTTCCG
ACGTGTGCTCTTCCG
GACGTGTGCTCTTCCG
ACGTGTGCTCTTCCG
GACGTGTGCTCTTCCG
ACGTGTGCTCTTCCG
GACGTGTGCTCTTCCG
ACGTGTGCTCTTCCG
GACGTGTGCTCTTCCG
GACGTGTGCTCTTCCG
GACGTGTGCTCTTCCG
GACGTGTGCTCTTCCG
ACGTGTGCTCTTCCG
ACGTGTGCTCTTCCG
GACGTGTGCTCTTCCG
GACGTGTGCTCTTCCG
ACGTGTGCTCTTCCG
GACGTGTGCTCTTCCG
ACGTGTGCTCTTCCG
GACGTGTGCTCTTCCG
GACGTGTGCTCTTCCG
ACGTGTGCTCTTCCG
ACGTGTGCTCTTCCG
ACGTGTGCTCTTCCG
GACGTGTGCTCTTCCG
ACGTGTGCTCTTCCG
GACGTGTGCTCTTCCG
ACGTGTGCTCTTCCG
GACGTGTGCTCTTCCG
GACGTGTGCTCTTCCG

ACGTGTGCTCTTCCG
ACGTGTGCTCTTCCG
GACGTGTGCTCTTCCG
GACGTGTGCTCTTCCG
ACGTGTGCTCTTCCG
ACGTGTGCTCTTCCG
ACGTGTGCTCTTCCG
GACGTGTGCTCTTCCG
GACGTGTGCTCTTCCG
ACGTGTGCTCTTCCG
GACGTGTGCTCTTCCG
ACGTGTGCTCTTCCG
GACGTGTGCTCTTCCG
GACGTGTGCTCTTCCG
ACGTGTGCTCTTCCG
GACGTGTGCTCTTCCG
GACGTGTGCTCTTCCG
ACGTGTGCTCTTCCG
ACGTGTGCTCTTCCG
GACGTGTGCTCTTCCG
ACGTGTGCTCTTCCG
GACGTGTGCTCTTCCG
GACGTGTGCTCTTCCG
GACGTGTGCTCTTCCG
ACGTGTGCTCTTCCG
ACGTGTGCTCTTCCG
GACGTGTGCTCTTCCG
GACGTGTGCTCTTCCG
GACGTGTGCTCTTCCG
GACGTGTGCTCTTCCG
ACGTGTGCTCTTCCG
ACGTGTGCTCTTCCG
GACGTGTGCTCTTCCG
CGTGTGCTCTTCCG
GACGTGTGCTCTTCCG
CGTGTGCTCTTCCG
GACGTGTGCTCTTCCG
GACGTGTGCTCTTCCG
ACGTGTGCTCTTCCG
ACGTGTGCTCTTCCG
ACGTGTGCTCTTCCG
GACGTGTGCTCTTCCG
ACGTGTGCTCTTCCG
GACGTGTGCTCTTCCG
ACGTGTGCTCTTCCG
ACGTGTGCTCTTCCG
GACGTGTGCTCTTCCG
ACGTGTGCTCTTCCG
ACGTGTGCTCTTCCG
GACGTGTGCTCTTCCG
ACGTGTGCTCTTCCG
ACGTGTGCTCTTCCG
GACGTGTGCTCTTCCG

ACGTGTGCTCTTCCG
GACGTGTGCTCTTCCG
GACGTGTGCTCTTCCG
GACGTGTGCTCTTCCG
ACGTGTGCTCTTCCG
ACGTGTGCTCTTCCG
GACGTGTGCTCTTCCG
ACGTGTGCTCTTCCG
ACGTGTGCTCTTCCG
GACGTGTGCTCTTCCG
GACGTGTGCTCTTCCG
ACGTGTGCTCTTCCG
GACGTGTGCTCTTCCG
ACGTGTGCTCTTCCG
GACGTGTGCTCTTCCG
ACGTGTGCTCTTCCG
GACGTGTGCTCTTCCG
GACGTGTGCTCTTCCG
CGTGTGCTCTTCCG
GACGTGTGCTCTTCCG
ACGTGTGCTCTTCCG
ACGTGTGCTCTTCCG
ACGTGTGCTCTTCCG
ACGTGTGCTCTTCCG
ACGTGTGCTCTTCCG
GACGTGTGCTCTTCCG
ACGTGTGCTCTTCCG
ACGTGTGCTCTTCCG
ACGTGTGCTCTTCCG
ACGTGTGCTCTTCCG
GACGTGTGCTCTTCCG
GACGTGTGCTCTTCCG
ACGTGTGCTCTTCCG
GACGTGTGCTCTTCCG
GACGTGTGCTCTTCCG
GACGTGTGCTCTTCCG
ACGTGTGCTCTTCCG
GACGTGTGCTCTTCCG
GACGTGTGCTCTTCCG
ACGTGTGCTCTTCCG
GACGTGTGCTCTTCCG
GACGTGTGCTCTTCCG
GACGTGTGCTCTTCCG
ACGTGTGCTCTTCCG
GACGTGTGCTCTTCCG
GACGTGTGCTCTTCCG
GACGTGTGCTCTTCCG
GACGTGTGCTCTTCCG
ACGTGTGCTCTTCCG
GACGTGTGCTCTTCCG
GACGTGTGCTCTTCCG
GACGTGTGCTCTTCCG
ACGTGTGCTCTTCCG

ACGTGTGCTCTTCCG
ACGTGTGCTCTTCCG
GACGTGTGCTCTTCCG
CGTGTGCTCTTCCG
GACGTGTGCTCTTCCG
GACGTGTGCTCTTCCG
ACGTGTGCTCTTCCG
ACGTGTGCTCTTCCG
GACGTGTGCTCTTCCG
ACGTGTGCTCTTCCG
CGTGTGCTCTTCCG
ACGTGTGCTCTTCCG
GACGTGTGCTCTTCCG
ACGTGTGCTCTTCCG
ACGTGTGCTCTTCCG
ACGTGTGCTCTTCCG
GACGTGTGCTCTTCCG
GACGTGTGCTCTTCCG
GACGTGTGCTCTTCCG
GACGTGTGCTCTTCCG
GACGTGTGCTCTTCCG
ACGTGTGCTCTTCCG
ACGTGTGCTCTTCCG
ACGTGTGCTCTTCCG
ACGTGTGCTCTTCCG
GACGTGTGCTCTTCCG
ACGTGTGCTCTTCCG
ACGTGTGCTCTTCCG
ACGTGTGCTCTTCCG
ACGTGTGCTCTTCCG
ACGTGTGCTCTTCCG
GACGTGTGCTCTTCCG
ACGTGTGCTCTTCCG
ACGTGTGCTCTTCCG
GACGTGTGCTCTTCCG
GACGTGTGCTCTTCCG
ACGTGTGCTCTTCCG
GACGTGTGCTCTTCCG
ACGTGTGCTCTTCCG
ACGTGTGCTCTTCCG
GACGTGTGCTCTTCCG
ACGTGTGCTCTTCCG
ACGTGTGCTCTTCCG
ACGTGTGCTCTTCCG
GACGTGTGCTCTTCCG
GACGTGTGCTCTTCCG
ACGTGTGCTCTTCCG
GACGTGTGCTCTTCCG
ACGTGTGCTCTTCCG
GACGTGTGCTCTTCCG
ACGTGTGCTCTTCCG
ACGTGTGCTCTTCCG
ACGTGTGCTCTTCCG

CGTGTGCTCTTCCG
GACGTGTGCTCTTCCG
ACGTGTGCTCTTCCG
GACGTGTGCTCTTCCG
GACGTGTGCTCTTCCG
ACGTGTGCTCTTCCG
ACGTGTGCTCTTCCG
GACGTGTGCTCTTCCG
ACGTGTGCTCTTCCG
GACGTGTGCTCTTCCG
ACGTGTGCTCTTCCG
GACGTGTGCTCTTCCG
GACGTGTGCTCTTCCG
ACGTGTGCTCTTCCG
ACGTGTGCTCTTCCG
ACGTGTGCTCTTCCG
GACGTGTGCTCTTCCG
ACGTGTGCTCTTCCG
GACGTGTGCTCTTCCG
GACGTGTGCTCTTCCG
GACGTGTGCTCTTCCG
ACGTGTGCTCTTCCG
ACGTGTGCTCTTCCG
CGTGTGCTCTTCCG
ACGTGTGCTCTTCCG
GACGTGTGCTCTTCCG
ACGTGTGCTCTTCCG
GACGTGTGCTCTTCCG
ACGTGTGCTCTTCCG
ACGTGTGCTCTTCCG
ACGTGTGCTCTTCCG
ACGTGTGCTCTTCCG
GACGTGTGCTCTTCCG
GACGTGTGCTCTTCCG
GACGTGTGCTCTTCCG
GACGTGTGCTCTTCCG
ACGTGTGCTCTTCCG
GACGTGTGCTCTTCCG
ACGTGTGCTCTTCCG
ACGTGTGCTCTTCCG
GACGTGTGCTCTTCCG
ACGTGTGCTCTTCCG
ACGTGTGCTCTTCCG
GACGTGTGCTCTTCCG
CGTGTGCTCTTCCG
ACGTGTGCTCTTCCG
CGTGTGCTCTTCCG
GACGTGTGCTCTTCCG
ACGTGTGCTCTTCCG
ACGTGTGCTCTTCCG
GACGTGTGCTCTTCCG
GACGTGTGCTCTTCCG
ACGTGTGCTCTTCCG

ACGTGTGCTCTTCCG
ACGTGTGCTCTTCCG
GACGTGTGCTCTTCCG
ACGTGTGCTCTTCCG
GACGTGTGCTCTTCCG
ACGTGTGCTCTTCCG
ACGTGTGCTCTTCCG
ACGTGTGCTCTTCCG
GACGTGTGCTCTTCCG
ACGTGTGCTCTTCCG
ACGTGTGCTCTTCCG
ACGTGTGCTCTTCCG
ACGTGTGCTCTTCCG
GACGTGTGCTCTTCCG
ACGTGTGCTCTTCCG
ACGTGTGCTCTTCCG
ACGTGTGCTCTTCCG
CGTGTGCTCTTCCG
GACGTGTGCTCTTCCG
ACGTGTGCTCTTCCG
ACGTGTGCTCTTCCG
ACGTGTGCTCTTCCG
GACGTGTGCTCTTCCG
ACGTGTGCTCTTCCG
ACGTGTGCTCTTCCG
GACGTGTGCTCTTCCG
GACGTGTGCTCTTCCG
ACGTGTGCTCTTCCG
ACGTGTGCTCTTCCG
ACGTGTGCTCTTCCG
ACGTGTGCTCTTCCG
GACGTGTGCTCTTCCG
ACGTGTGCTCTTCCG
GACGTGTGCTCTTCCG
GACGTGTGCTCTTCCG
GACGTGTGCTCTTCCG
ACGTGTGCTCTTCCG
GACGTGTGCTCTTCCG
GACGTGTGCTCTTCCG
GACGTGTGCTCTTCCG
GACGTGTGCTCTTCCG
ACGTGTGCTCTTCCG
GACGTGTGCTCTTCCG
GACGTGTGCTCTTCCG
GACGTGTGCTCTTCCG
ACGTGTGCTCTTCCG
GACGTGTGCTCTTCCG
GACGTGTGCTCTTCCG
ACGTGTGCTCTTCCG
CGTGTGCTCTTCCG
GACGTGTGCTCTTCCG
GACGTGTGCTCTTCCG
GACGTGTGCTCTTCCG

GACGTGTGCTCTTCCG
GACGTGTGCTCTTCCG
ACGTGTGCTCTTCCG
GACGTGTGCTCTTCCG
GACGTGTGCTCTTCCG
GACGTGTGCTCTTCCG
GACGTGTGCTCTTCCG
ACGTGTGCTCTTCCG
GACGTGTGCTCTTCCG
GACGTGTGCTCTTCCG
ACGTGTGCTCTTCCG
GACGTGTGCTCTTCCG
ACGTGTGCTCTTCCG
ACGTGTGCTCTTCCG
GACGTGTGCTCTTCCG
GACGTGTGCTCTTCCG
CGTGTGCTCTTCCG
ACGTGTGCTCTTCCG
GACGTGTGCTCTTCCG
ACGTGTGCTCTTCCG
GACGTGTGCTCTTCCG
GACGTGTGCTCTTCCG
GACGTGTGCTCTTCCG
ACGTGTGCTCTTCCG
ACGTGTGCTCTTCCG
ACGTGTGCTCTTCCG
ACGTGTGCTCTTCCG
ACGTGTGCTCTTCCG
ACGTGTGCTCTTCCG
GACGTGTGCTCTTCCG
ACGTGTGCTCTTCCG
ACGTGTGCTCTTCCG
ACGTGTGCTCTTCCG
ACGTGTGCTCTTCCG
GACGTGTGCTCTTCCG
ACGTGTGCTCTTCCG
ACGTGTGCTCTTCCG
ACGTGTGCTCTTCCG
ACGTGTGCTCTTCCG
ACGTGTGCTCTTCCG
GACGTGTGCTCTTCCG
ACGTGTGCTCTTCCG
GACGTGTGCTCTTCCG
ACGTGTGCTCTTCCG
ACGTGTGCTCTTCCG
ACGTGTGCTCTTCCG
GACGTGTGCTCTTCCG
ACGTGTGCTCTTCCG
ACGTGTGCTCTTCCG
ACGTGTGCTCTTCCG
GACGTGTGCTCTTCCG
ACGTGTGCTCTTCCG
CGTGTGCTCTTCCG

GACGTGTGCTCTTCCG
GACGTGTGCTCTTCCG
ACGTGTGCTCTTCCG
GACGTGTGCTCTTCCG
CGTGTGCTCTTCCG
GACGTGTGCTCTTCCG
ACGTGTGCTCTTCCG
GACGTGTGCTCTTCCG
GACGTGTGCTCTTCCG
ACGTGTGCTCTTCCG
GACGTGTGCTCTTCCG
ACGTGTGCTCTTCCG
GACGTGTGCTCTTCCG
ACGTGTGCTCTTCCG
ACGTGTGCTCTTCCG
ACGTGTGCTCTTCCG
GACGTGTGCTCTTCCG
ACGTGTGCTCTTCCG
GACGTGTGCTCTTCCG
ACGTGTGCTCTTCCG
CGTGTGCTCTTCCG
ACGTGTGCTCTTCCG
ACGTGTGCTCTTCCG
ACGTGTGCTCTTCCG
GACGTGTGCTCTTCCG
GACGTGTGCTCTTCCG
CGTGTGCTCTTCCG
GACGTGTGCTCTTCCG
ACGTGTGCTCTTCCG
ACGTGTGCTCTTCCG
GACGTGTGCTCTTCCG
GACGTGTGCTCTTCCG
ACGTGTGCTCTTCCG
GACGTGTGCTCTTCCG
ACGTGTGCTCTTCCG
ACGTGTGCTCTTCCG
GACGTGTGCTCTTCCG
GACGTGTGCTCTTCCG
GACGTGTGCTCTTCCG
ACGTGTGCTCTTCCG
ACGTGTGCTCTTCCG
ACGTGTGCTCTTCCG
GACGTGTGCTCTTCCG
ACGTGTGCTCTTCCG
CGTGTGCTCTTCCG
GACGTGTGCTCTTCCG
ACGTGTGCTCTTCCG
ACGTGTGCTCTTCCG
ACGTGTGCTCTTCCG
ACGTGTGCTCTTCCG
GACGTGTGCTCTTCCG
ACGTGTGCTCTTCCG
GACGTGTGCTCTTCCG

ACGTGTGCTCTTCCG
ACGTGTGCTCTTCCG
ACGTGTGCTCTTCCG
GACGTGTGCTCTTCCG
ACGTGTGCTCTTCCG
ACGTGTGCTCTTCCG
ACGTGTGCTCTTCCG
ACGTGTGCTCTTCCG
ACGTGTGCTCTTCCG
CGTGTGCTCTTCCG
GACGTGTGCTCTTCCG
ACGTGTGCTCTTCCG
CGTGTGCTCTTCCG
ACGTGTGCTCTTCCG
GACGTGTGCTCTTCCG
ACGTGTGCTCTTCCG
ACGTGTGCTCTTCCG
ACGTGTGCTCTTCCG
GACGTGTGCTCTTCCG
GACGTGTGCTCTTCCG
GACGTGTGCTCTTCCG
ACGTGTGCTCTTCCG
GACGTGTGCTCTTCCG
ACGTGTGCTCTTCCG
ACGTGTGCTCTTCCG
ACGTGTGCTCTTCCG
GACGTGTGCTCTTCCG
ACGTGTGCTCTTCCG
GACGTGTGCTCTTCCG
GACGTGTGCTCTTCCG
ACGTGTGCTCTTCCG
GACGTGTGCTCTTCCG
ACGTGTGCTCTTCCG
GACGTGTGCTCTTCCG
GACGTGTGCTCTTCCG
GACGTGTGCTCTTCCG
ACGTGTGCTCTTCCG
GACGTGTGCTCTTCCG
GACGTGTGCTCTTCCG
ACGTGTGCTCTTCCG
GACGTGTGCTCTTCCG
ACGTGTGCTCTTCCG
GACGTGTGCTCTTCCG
GACGTGTGCTCTTCCG
ACGTGTGCTCTTCCG
GACGTGTGCTCTTCCG
GACGTGTGCTCTTCCG
GACGTGTGCTCTTCCG
ACGTGTGCTCTTCCG
ACGTGTGCTCTTCCG
ACGTGTGCTCTTCCG
ACGTGTGCTCTTCCG
GACGTGTGCTCTTCCG

GACGTGTGCTCTTCCG
GACGTGTGCTCTTCCG
GACGTGTGCTCTTCCG
ACGTGTGCTCTTCCG
GACGTGTGCTCTTCCG
GACGTGTGCTCTTCCG
GACGTGTGCTCTTCCG
GACGTGTGCTCTTCCG
GACGTGTGCTCTTCCG
GACGTGTGCTCTTCCG
GACGTGTGCTCTTCCG
ACGTGTGCTCTTCCG
ACGTGTGCTCTTCCG
ACGTGTGCTCTTCCG
GACGTGTGCTCTTCCG
GACGTGTGCTCTTCCG
ACGTGTGCTCTTCCG
ACGTGTGCTCTTCCG
ACGTGTGCTCTTCCG
ACGTGTGCTCTTCCG
GACGTGTGCTCTTCCG
CGTGTGCTCTTCCG
ACGTGTGCTCTTCCG
ACGTGTGCTCTTCCG
ACGTGTGCTCTTCCG
GACGTGTGCTCTTCCG
ACGTGTGCTCTTCCG
GACGTGTGCTCTTCCG
ACGTGTGCTCTTCCG
GACGTGTGCTCTTCCG
ACGTGTGCTCTTCCG
GACGTGTGCTCTTCCG
ACGTGTGCTCTTCCG
GACGTGTGCTCTTCCG
GACGTGTGCTCTTCCG
ACGTGTGCTCTTCCG
GACGTGTGCTCTTCCG
ACGTGTGCTCTTCCG
ACGTGTGCTCTTCCG
ACGTGTGCTCTTCCG

NNNNNNNNNNNNNNNAGATCGGAAGAGCACACGTCTG
NNNNNNNNNNNNNNNNNNNNNNNNNNNNNNNNNNNNNNNNNNNNNNNNNNNNNNNNNNNNNNN

GTAGACAGCTCTAGCACCGCTTAAACGCACGTACGCGCTGTCCCCCGCGTTTTAACCGCCAAG

NNNNNAGATCGGAAGAGCACACGTCTG

GGGATTACTCCCTAGTCTCCAGGCACGTNNNNNNNNNNNNNNNNNNNNNNNNNNNNNNNNNNNN

NNNNNNNNNNNNNNNNNNNNNNNNNNNNNNNNNNAGATCGGAAGAGCACACGTCTG
